# Metabolic Potential of Microbial Communities in the Hypersaline Sediments of the Bonneville Salt Flats

**DOI:** 10.1101/2021.10.18.464844

**Authors:** Julia M. McGonigle, Jeremiah A. Bernau, Brenda B. Bowen, William J. Brazelton

## Abstract

The Bonneville Salt Flats (BSF) appear to be entirely desolate when viewed from above, but in reality they host rich microbial communities just below the surface salt crust. In this study, we investigate the metabolic potential of the BSF microbial ecosystem. The predicted and measured metabolic activities provide new insights into the ecosystem functions of evaporite landscapes and are an important analog for potential subsurface microbial ecosystems on ancient and modern Mars. Hypersaline and evaporite systems have been investigated previously as astrobiological analogs for Mars and other salty celestial bodies. Still, these studies have generally focused on aquatic systems and cultivation-dependent approaches. Here, we present an ecosystem-level examination of metabolic pathways within the shallow subsurface of evaporites. We detected aerobic and anaerobic respiration as well as methanogenesis in BSF sediments. Metagenome-assembled genomes (MAGs) of diverse bacteria and archaea encoded a remarkable diversity of metabolic pathways, including those associated with carbon fixation, carbon monoxide oxidation, acetogenesis, methanogenesis, sulfide oxidation, denitrification, and nitrogen fixation. These results demonstrate the potential for multiple energy sources and metabolic pathways in BSF and highlight the possibility for vibrant microbial ecosystems in the shallow subsurface of evaporites.

## 1. Introduction

Evaporite deposits are valuable analogs for investigating the potential for ancient and modern life in saline conditions of extraterrestrial settings, including on Mars (Barbieri and Stivaletta, 2011). Jezero Crater, the landing site for the Mars Perseverance Rover, contains sedimentary deposits expected to be the remnants of an ancient, now desiccated lake (Goudge et al., 2015; Horgan et al., 2020). Similarly, the Bonneville Salt Flats (BSF) in Utah, USA, are the remnants of Lake Bonneville, the largest lake in western North America before it began to desiccate ~15,000 years ago (Oviatt, 2015). BSF is famous for its other-worldly, barren appearance and for hosting vehicle land speed records. In addition, the extensive salt deposits are an economically significant source of potash.

Despite the sterile appearance of BSF, the salt deposits host remarkably robust microbial communities that are visibly evident just below the top layer of crust. The archaeal and bacterial taxa composition varies with sediment depth (spanning a few centimeters) but is remarkably consistent across a horizontal transect of several kilometers (McGonigle et al., 2019). Similar microbial diversity results have been reported for hypersaline sediments in the nearby Pilot Valley basin, another remnant of Lake Bonneville (Lynch et al., 2019b, 2019a).

The seasonal availability of water may stimulate the density and diversity of BSF microbial communities. The salt flats are typically flooded during the winter and spring months and then rapidly desiccate during the summer (Bowen et al., 2017). This annual cycle of flooding, evaporation, and desiccation contrasts with the desert salt flats of Death Valley (Hunt et al., 1966), Tunisia (Bryant and Rainey, 2002), and the Arabian Peninsula (Schulz et al., 2015) and to the hyperarid soils of the Atacama Desert (Kampf and Tyler, 2006; Azua-Bustos et al., 2015; Wierzchos et al., 2015), where flooding events are rare.

Microbial diversity studies of these hyperarid, hypersaline systems have confirmed the presence of resident microbial communities with both cultivation-dependent and -independent studies (Lynch, 2015; McGonigle et al., 2019; Myers and King, 2020). A few studies have investigated these systems for their ability to support specific metabolic activities, such as carbon monoxide oxidation (King, 2015; Myers and King, 2020), perchlorate reduction (Lynch et al., 2019a), and hypolithic photosynthesis (Lacap-Bugler et al., 2017). However, the metabolic activity, nutrient cycling, and genomic content of these microbial communities have not been studied at the level of the whole ecosystem. Here, we report our exploration of the metabolic potential of the BSF microbial ecosystem with incubation experiments and metagenome sequencing.

## 2. Materials and Methods

### 2.1. Sampling and DNA Extraction

Sampling location, elemental analysis, and DNA extraction methods are described in detail by McGonigle et al., 2019. Briefly, sediment samples were collected from the upper 30 cm of sediments at eight sites distributed across BSF in September 2016. Layers were sampled via visual identification of color and textural characteristics at each site. Samples were transported on ice within 12 hours to the University of Utah, where they were stored at –80°C. Microbiological analyses, including DNA extraction, were performed in the following six months. DNA extraction was performed using a protocol from Brazelton et al., 2010 modified for highly saline material. The modified protocol was published by McGonigle et al., 2019, archived as DOI: 10.5281/zenodo.2827066, and is summarized here. A sterile mortar and pestle were used to crush collected sediments. A 0.25 g subsample was extracted using a buffer containing 0.1 M Tris, 0.1 M EDTA, 0.1 M KH2PO4, 0.8 M guanidium HCl, and 0.5% Triton X-100. Samples were then subjected to one freeze-thaw cycle, incubation at 65°C for 30 min, and beating with 0.1 mm glass beads in a Mini-Beadbeater-16 (Biospec Products) to lyse. Purification was performed via extraction with phenol-chloroform-isoamyl alcohol, precipitation in ethanol, washing in Amicon 30K Ultra centrifugal filters, and final cleanup with 2× SPRI beads (Rohland and Reich, 2012). DNA quantification was performed with a Qubit fluorometer (Thermo Fisher).

### 2.2. Stable Isotope Incubation Experiments

Incubation samples were collected alongside DNA sequencing samples at sites 12B, 33, 41, and 56. Sediment from individual layers was placed into a sterile plastic tube containing 40 mL of either oxic or anoxic 18% artificial seawater solution. The anoxic solution was made in an anoxic chamber (Coy Laboratory Products) with 0.05% dithiothreitol (DTT) added to help maintain anoxic conditions in the field. Additionally, sterile tubes were filled with liquid and kept wrapped with parafilm to minimize oxygen introduction into the tube. Collected samples were placed in a foam cooler inside a cooler filled with ice to prevent freezing and transported back to the University of Utah, where incubations were set up on the same day as sampling.

Anaerobic incubations were prepared in the anoxic chamber; aerobic incubations were prepared on the lab bench. Sediment samples were vortexed to break apart sediments and to create a sediment slurry. 2 mL of the sediment slurry was pipetted into an exetainer (LabCo) tube containing 2 mL of anoxic or oxic saltwater media amended with 0.07 M solution of bicarbonate, glucose, or acetate, each of which was 25% labeled with ^13^C (i.e. mixed 1:4 with 25% H^13^CO_3_, ^13^C_12_H_6_O_12_, or CH_3_^13^CO_2_, respectively; Cambridge Isotope Laboratories). All saltwater media was adapted from the Halohandbook and contained NaCl, MgCl_2_·6 H_2_O, MgSO_4_·7 H_2_O, KCl, TrisHCl and included B vitamins (thiamine and biotin) and the following trace elements: MnCl_2_·4H_2_O, ZnSO_4_·7 H_2_O, FeSO_4_·7 H_2_O, and CuSO_4_·5 H_2_O (Dyall-Smith, 2009). Around 10 mL of hydrogen gas was added to half of the bicarbonate incubations using a syringe. Incubations were conducted at room temperature in either light (aerobic) or dark (anaerobic) conditions for 60 days. After this time, around 7 mL of the headspace from each exetainer tube was sampled with a syringe and transferred into two clean exetainers flushed with nitrogen gas in the anoxic chamber. Exetainers containing the sampled headspace were sent to the UC Davis Stable Isotope Facility for measurements of ^13^CH_4_ and ^13^CO_2_ (ThermoScientific GasBench system interfaced to a ThermoScientific Delta V Plus isotope ratio mass spectrometer).

### 2.3. Metagenome Sequencing and Analysis

Metagenomic DNA was fragmented to ~500-700 bp using a Qsonica Q800R sonicator. According to the manufacturer’s instructions, 500 ng of fragmented DNA was used to construct metagenome libraries using the NEBNext Ultra DNA library preparation kit for Illumina. The University of Utah High-Throughput Genomics Core Facility performed quality control and sequencing of the metagenomic libraries. Quality was evaluated with a Bioanalyzer DNA 1000 chip (Agilent Technologies), and then paired-end sequencing (2 × 125 bp) was performed on an Illumina HiSeq2500 platform with HiSeq (v4) chemistry. The 24 metagenome libraries were multiplexed with eight libraries per one Illumina lane, yielding a total of 845 million paired reads for all 24 libraries before quality filtering. Demultiplexing and conversion of the raw sequencing base-call data were performed through the CASAVA (v1.8) pipeline.

Raw sequences were processed by the Brazelton lab to trim adapter sequences with BBDuk (part of the BBTools suite, v35.85 (Bushnell et al., 2017)), to remove artificial replicates, and to trim the reads based on quality. Removal of replicates and quality trimming were performed with our seq-qc package (https://github.com/Brazelton-Lab/seq-qc). At this stage, 8 of the libraries were determined to be of very low quality and were not included in further analyses. The remaining 16 libraries had 600 million paired reads after quality filtering. Paired-end reads were assembled with MegaHit v1.1.1 (Li et al., 2016), using kmers of 27 to 141. Prodigal v2.6.3 (Hyatt et al., 2010) was run in the anonymous gene prediction mode to identify open reading frames. Functional annotation was performed using the blastp function of Diamond v0.9.14 (Buchfink et al., 2021) with both the prokaryotes and T10000 (addendum annotations) databases from KEGG release 83.2 with an E value of 1e-6. Annotations were selected by the highest quality alignment, as determined by the bit score. Coverage was determined through read mapping with bowtie2 v2.3.2 (Langmead and Salzberg, 2012) and bedtools v2.25.0 (Quinlan and Kindlon, 2019). ANI was performed with fastANI v0.1.2 (https://github.com/ParBLiSS/FastANI) through the online data platform KBase (accessed July, 2020).

### 2.4. Metagenome Binning

Binning of individual samples resulted in poor representation of diverse taxa identified through previous 16S rRNA studies and recovered quality genomes of only a few taxa. Because our previous work showed a striking similarity in microbial community composition along a horizontal transect (McGonigle et al., 2019), we pooled contigs from multiple assemblies to improve metagenomic binning efforts. Metagenomic binning was performed on the pooled contigs using BinSanity (Graham et al., 2017). Bins were evaluated using CheckM (Parks et al., 2015) and taxonomy was assigned using GTDB-Tk Classify through KBase (Chaumeil et al., 2020). As with unbinned contigs, functional annotation was performed using the blastp function of Diamond v0.9.14 (Buchfink et al., 2021) with both the prokaryotes and T10000 (addendum annotations) databases from KEGG release 83.2. The online KEGG mapper tool was used to identify bins with more complete metabolic pathways of interest. MAGs were assigned taxonomic classifications with GTDB-Tk classify (kb_gtdbtk v.0.1.4) (Chaumeil et al., 2020) as implemented by the KBase platform (Arkin et al., 2018).

### 2.5. Data availability

All unassembled sequences related to this study are available at the NCBI Sequence Read Archive (BioProject accession number PRJNA522308). All metagenome assembled genomes (MAGs) may be found under NCBI BioSample accession numbers SAMN19270115 - SAMN19270142. All supplementary materials, solution recipes, and protocols are archived at https://doi.org/10.5281/zenodo.5569980. All custom software and scripts are available at https://github.com/Brazelton-Lab.

## 3. Results

### 3.1. Stable Isotope Incubation Experiments

The capacity for BSF sediments to support respiration was measured as the generation of ^13^C-labeled carbon dioxide from ^13^C-labeled glucose or acetate during incubation of sediment samples at room temperature for 60 days. Production of CO_2_ from both glucose and acetate was detected in all incubations, including all layers from all four locations. Levels of CO_2_ production were roughly equivalent across all incubations (***Table 1***). Our experiment was designed as an activity assay and was not conducive to calculating a precise quantity of carbon released per gram of sediment or per unit time.

**Table 1.**
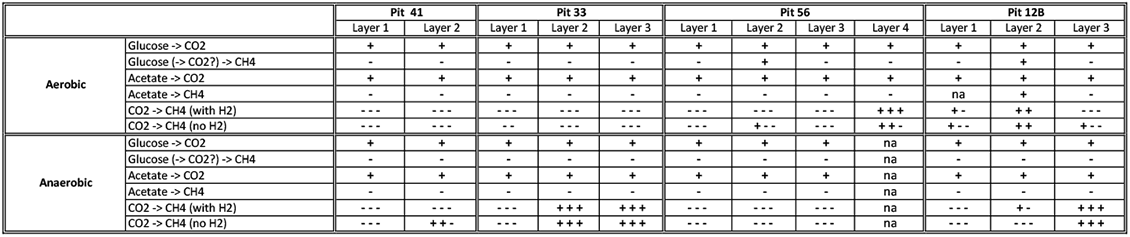
Headspace results for presence/absence (indicated by +/- respectively) of ^13^C conversions in aerobic and anaerobic incubations. Multiple +/- values indicate replicate treatments, na indicates data is not available.

Methanogenesis in BSF sediments was measured during replicate incubations as the generation of ^13^C-labeled methane from ^13^C-labeled bicarbonate, glucose, or acetate. Production of methane from bicarbonate was detected in all sites (***Figure S1***). Evidence of methane production from carbon originating in glucose or acetate was only seen in aerobic incubations from the lower horizons sampled at sites 12B and 56 (***Table 1***). Methane production from bicarbonate was highest in aerobic incubations, with no added hydrogen, from sediments sampled from the layer just beneath the surface halite at site 12B. At this site, methanogenesis occurred mostly in these second layer aerobic incubations, but also in anaerobic incubations from the next deepest layer. Site 56 had methane production only in aerobic incubations, while site 33 had methane production mostly in anaerobic incubations. Incubations from site 33 also had the most consistent methane production trend across all replicates compared to other sites (***Figure S1***).

### 3.2. BSF Metagenome-Assembled Genomes (MAGs)

A summary of metagenome sequencing and assembly results is reported in ***Table S1***. Sediment samples were grouped according to their mineralogy (group 1: surface halite, group 2: upper gypsum, group 3: lower halite, and group 4: halite mixed with gypsum) as described in detail in McGonigle et al., 2019. The metagenomes constructed from all samples in group 4 were among those with poor read quality and are not reported here. Sequencing results for the 16 remaining libraries were broadly similar across sites and mineralogical groups, ranging from 25 to 55 million paired reads per library. Assemblies included 700,000 to 1.6 million predicted proteins, approximately 30% of which could be annotated with the KEGG database. Sample-specific assemblies were pooled for binning purposes, resulting in 43 MAGs with <15% contamination and >45% completion after automated binning. Of those, 28 with the most complete pathways involved in biogeochemical cycles are reported here (***Figure 1, Table S2***).

**Figure 1.**
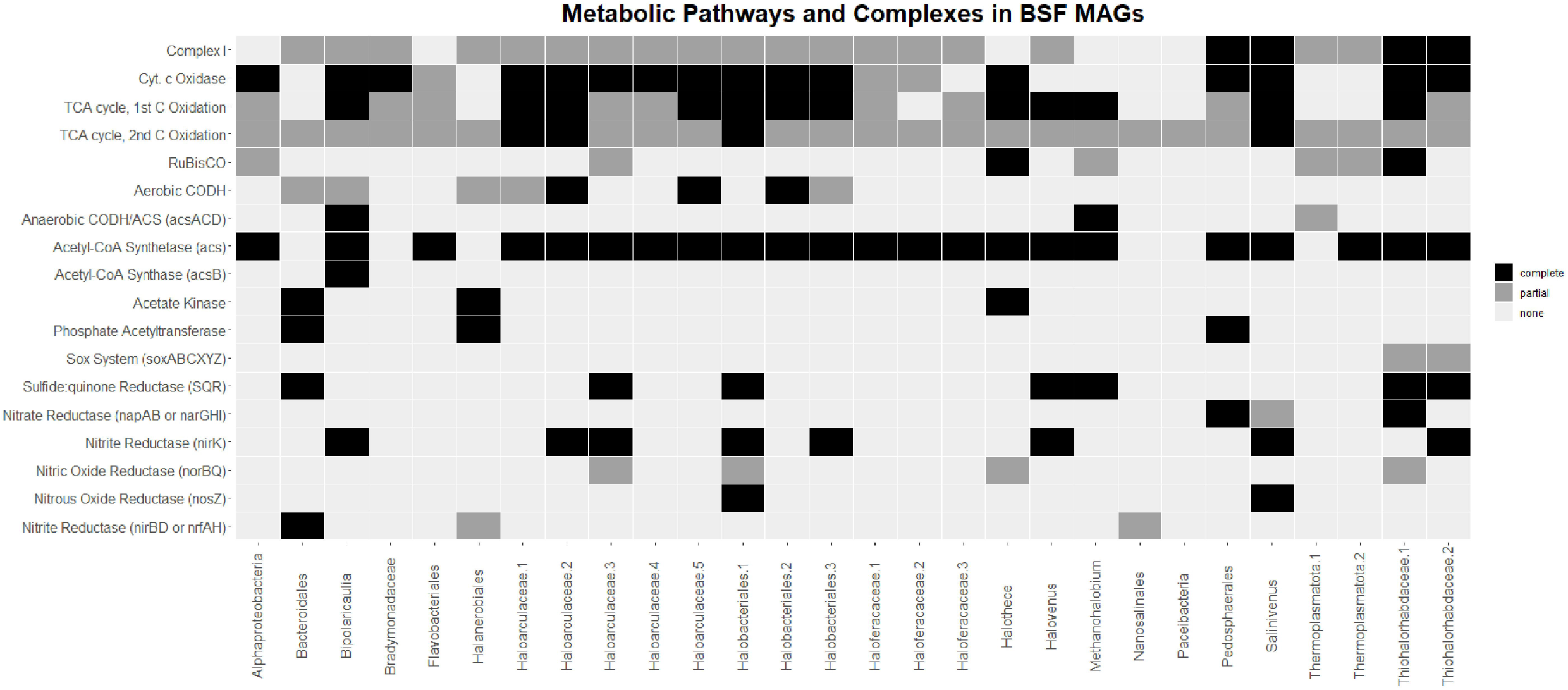
Completion of various metabolic pathways and complexes in the BSF MAGs. Black indicates a complete pathway/complex, dark grey indicates a partial pathway/complex, and light grey indicates an absence of any genes in the pathway/complex. Pathways/complexes correspond to the following KEGG modules and IDs: Complex I (M00144), Cyt. c Oxidase (M00155), TCA cycle, 1st C Oxidation (M00010), TCA cycle, 2nd C Oxidation (M00011), RuBisCO (K01601, K01602), Aerobic CODH (K03518, K03519, K03520), Anaerobic CODH/ACS (K00194, K00197, K00198), Acetate Kinase (K00925), Phosphate Acetyltransferase (K00625)

### 3.3. Carbon Fixation Pathways

We found evidence for multiple carbon fixation pathways in the BSF metagenomes, including the reductive pentose phosphate pathway and the Wood-Ljungdahl pathway (***Figure S2***, ***Figure 2***). We also found evidence for carbon monoxide oxidation, which can serve as a carbon source for autotrophy and/or supplemental electrons for energy conservation.

**Figure 2.**
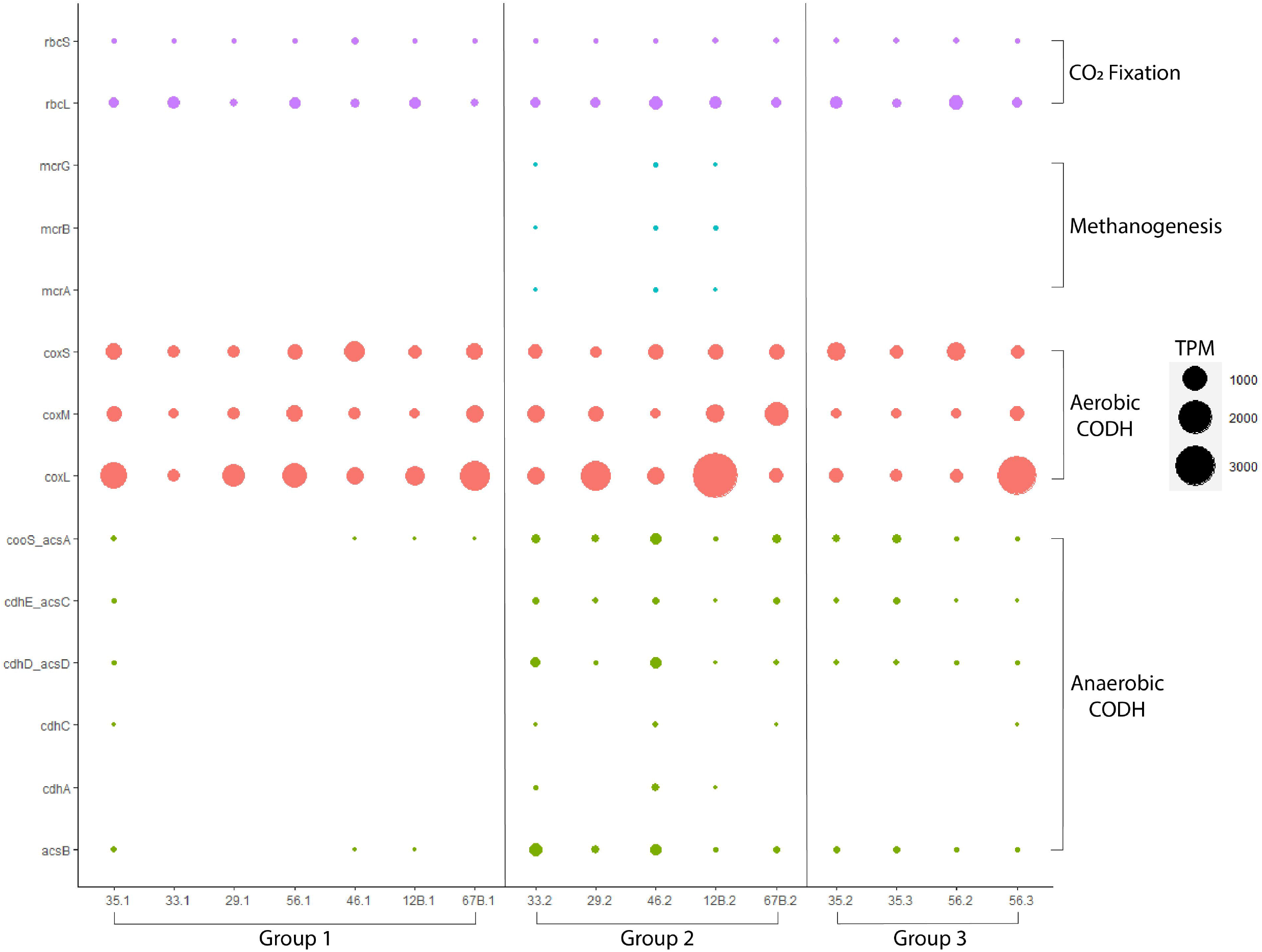
Abundance of key carbon cycling genes in BSF metagenomes (values in transcripts per million - *TPM*).

The reductive pentose phosphate pathway (i.e., the Benson-Calvin cycle, RPP) features two key enzymes: ribulose-1,5-bisphosphate carboxylase/oxygenase (RuBisCO) and phosphoribulokinase. Both small and large subunits are distributed across all mineralogical groups, but the large subunit is generally more abundant (***Figure 2***). The small subunit of RuBisCO appears heterogeneously distributed across all samples, but the large subunit is slightly more abundant with coverage greater than 144 TPM in surface halite crust at site 33 (group 1) and in lower layers from sites 46, 12B, 56, and 35 (groups 2 and 3) (***Figure 2, Table S3***). A complete KEGG module for the RPP pathway is found in all of our metagenomes. Genes encoding both small and large subunits of RuBisCO in addition to phosphoribulokinase are present in only two of our MAGs: the cyanobacteria *Halothece*, which contains a complete RPP pathway and *Thiohalorhabdaceae*.1, which is only missing fructose-1,6-bisphosphatase (***Figure 1***). The large subunit is found in numerous MAGs, but many are missing the small subunit and phosphoribulokinase, another key enzyme in the RPP.

A key enzyme of the Wood-Ljungdahl (WL) pathway is anaerobic CO dehydrogenase/acetyl-CoA synthase (CODH/ACS), a five-subunit complex. Genes *acsCD* encode the corrinoid iron-sulfur protein component of the complex responsible for reactions in the methyl branch of the WL pathway, and acsAB encode the catalytic component responsible for CO_2_ reduction and acetyl-coA synthesis. Although not very abundant in most samples, these genes are present in our metagenomes, mostly from mineralogical groups 2 and 3 (***Figure 2***). The metagenome constructed from group 2 at site 46, located just under the salt crust, is the only sample with a complete suite of CODH/ACS genes. Most of these genes have the highest abundance in this sample. The group 2 metagenome from site 33 has a comparably high abundance of anaerobic CODH/ACS genes and is only missing *cdhB* (the epsilon subunit) (***Table S3***). All metagenomes constructed from groups 2 and 3 contain a complete KEGG module for the WL pathway. In contrast, only one metagenome from group 1, the salt crust samples, contained a complete module for the pathway (site 35); the others are missing methyltransferase (*acsE*) and *acsCD*. Additionally, a commonly used marker gene for the WL pathway, acetyl-CoA synthase (*acsB*), is found in group 1 only at sites 12B, 46, and 35, but is present in all metagenomes from groups 2 and 3.

Some MAGs captured a partial anaerobic CODH/ACS complex, but only the *Bipolaricaulia* MAG contained genes encoding all crucial parts of the complex, including both catalytic units (***Figure 1***). This *Bipolaricaulia* MAG also contained the most complete WL pathway, only missing genes encoding formate dehydrogenase. The *Thermoplasmatota*.1 MAG contained the bacterial form of the catalytic subunits of CODH/ACS, but is missing additional parts of the complex. This MAG contained only two other genes in the WL pathway: methylenetetrahydrofolate dehydrogenase (*folD*) and formate-tetrahydrofolate ligase (*fhs*). This may be due to the incomplete representation of the genome in our data as the *Thermoplasmatota*.1 MAG is only 52% complete (***Table S2***).

#### 3.3.1. Methanogenesis

Genes for methanogenesis were not abundant across the BSF samples, except for *mttB*, which encodes a trimethylamine methyltransferase used in methanogenesis from trimethylamines (***Figure S2, Table S3***). In all metagenomes, there was an absence of methylenetetrahydromethanopterin dehydrogenase (*hmd*), which is present in some methanogens reliant on hydrogen gas as an electron donor (Thauer et al., 2010). A complete suite of genes for methanogenesis from carbon dioxide (excepting *hmd*) was only found in two of the BSF samples: group 2 metagenomes from sites 33 and 46. However, the metagenome from group 2 located just under the salt crust at site 12B had a near-complete pathway for methanogenesis from acetate, -methylamines, and methanol with only *mtrCDEH* (tetrahydromethanopterin S-methyltransferase) and *mtd* (methylenetetrahydromethanopterin dehydrogenase) missing. The key enzyme for methanogenesis, methyl-coenzyme M reductase (*mcr*), was not found in any other metagenomes (***Figure 2***).

We obtained only one MAG, classified as genus *Methanohalobium*, with a near-complete pathway for methanogenesis. The MAG contained genes required for methanogenesis from CO_2_, methylamines, methanol, and acetate, but it did not include any hydrogenases.

#### 3.3.2. CO Oxidation

The enzyme aerobic carbon monoxide dehydrogenase (CODH) enables the use of carbon monoxide (CO) as an energy and/or carbon source in oxic conditions. The three genes encoding aerobicCODH (*coxSML*) were among the most abundant genes in all mineralogical groups (***Figure S2***). The gene encoding the catalytic subunit (*coxL*) was particularly abundant in mineralogical group 1 at sites 35 and 67B, but at the highest abundance just under the surface halite at site 12B (group 2) (***Figure 2***). In group 3, *coxL* was most abundant at site 56. Of the 28 MAGs obtained, 8 contain at least one of the three aerobic CODH genes (***Figure 1***), but the catalytic large subunit was only found in bins belonging to haloarchaea. The bacterial *Bacteroidales* and *Bipolaricaulia* bins both contained genes encoding the small subunit, and the *Haloanerobiales* bin contained genes encoding the small and medium subunits.

The genes encoding form I CODH typically appear in an operon as *coxMSL*, while form II CODH genes are either organized as *coxSLM* or not contained in an operon (King and Weber, 2007). The complete form I CODH operon is found in our *Haloarculaceae*.5 and *Haloarculaceae*.2 MAGs (***Table S4***). These two MAGs plus a third (*Haloarculaceae*.2) encode additional sets of genes annotated as *coxSLM* or *coxLM*, matching the operon structure of form II CODH. However, these genes lack the AYRGAGR active site motif characteristic of previously reported form II CODH genes (King and Weber, 2007). Interestingly, none of our MAGs containing an aerobic CODH encoded additional genes with annotations associated with carbon fixation.

CO oxidation by *Archaea* has recently been described with isolates cultured from BSF (King, 2015; Myers and King, 2020). We calculated average nucleotide identity (ANI) among our haloarchaea bins and the available genomes for these CO-oxidizing *Archaea* (***Table S5***). The best match between the three recently characterized isolates and our MAGs was 79.93% (for *Halovenus carboxidivorans*; ***Table S5***), indicating that our MAGs probably represent distinct species or genera compared to the previously described CO-oxidizing species (King, 2015; Myers and King, 2020).

### 3.4. Nitrogen Fixation

Genomic evidence for nitrogen fixation was present in four of the seven sites (12B, 67B, 33, and 35). Site 12B had the highest abundance of nitrogenase genes (*nifDHK*) (***Table S3***). Unfortunately, no nitrogenase genes were present in a high-quality MAG, so we investigated individual nitrogenase sequences present in the metagenomes by querying them against the NCBI nr database. Six nitrogenase sequences from the group 1 surface halite sample collected at site 12B had high sequence similarity (~55%-75% amino acid identities) to *nifDH* genes previously identified in bacterial *Chloroflexi* (*Roseiflexus*) species. However, all nifK genes found in this sample had the highest sequence similarity to metagenomic sequences representing an unclassified archaeon (***Table S6***). In lower sediment layers (groups 2 and 3), nitrogenase genes are most similar to those previously described in purple non-sulfur bacteria or sulfate-reducing bacteria (*Deltaproteobacteria*). Sites 12B and 35 had one *nifH* and *nifK* gene (respectively) with sequence similarity to *Halorhodospira* (purple sulfur bacteria) species.

#### 3.4.1. Nitrification

Hydroxylamine dehydrogenase (*hao*), involved in the first step of nitrification, was highly abundant in the deepest sample sequenced from site 35 (group 3). It is found elsewhere only at reduced levels in the lower layers at sites 35 and 56 (group 3) (***Table S3***). Evidence for other genes associated with nitrification is limited. Genes annotated as methane/ammonia monooxygenase (*pmoBC/amoBC*) were also detected at low levels in the deeper layers of sites 35 and 56. Still, additional evidence is required to distinguish whether these genes are involved in the oxidation of methane or ammonia, as KEGG orthology alone is not sufficient. These genes were only found in one MAG (*Thiohalorhabdaceae*.2), which is 59% complete and lacked the alpha subunit (***Table S2***).

KEGG orthology alone cannot distinguish nitrite oxidoreductase (*nxr*; involved in nitrification of nitrite to nitrate) from nitrate reductase (*nar*; the reverse reaction involved in denitrification). Genes annotated as the alpha and beta subunits of these enzymes (*nxrAB*/*narGH*) were present in all samples and highly abundant in mineralogical groups 2 and 3 below the surface halite (***Figure 3***). Again, the only MAG to contain both of these genes represents a *Thiohalorhabdaceae* (***Figure 4***), and this MAG also contains the gamma subunit of the enzyme specific to nitrate reductase (*narI*). The type species of this group is *Thiohalorhabdus denitrificans*, an extremely halophilic chemolithoautotroph capable of denitrification but not nitrification (Sorokin et al., 2008).

**Figure 3.**
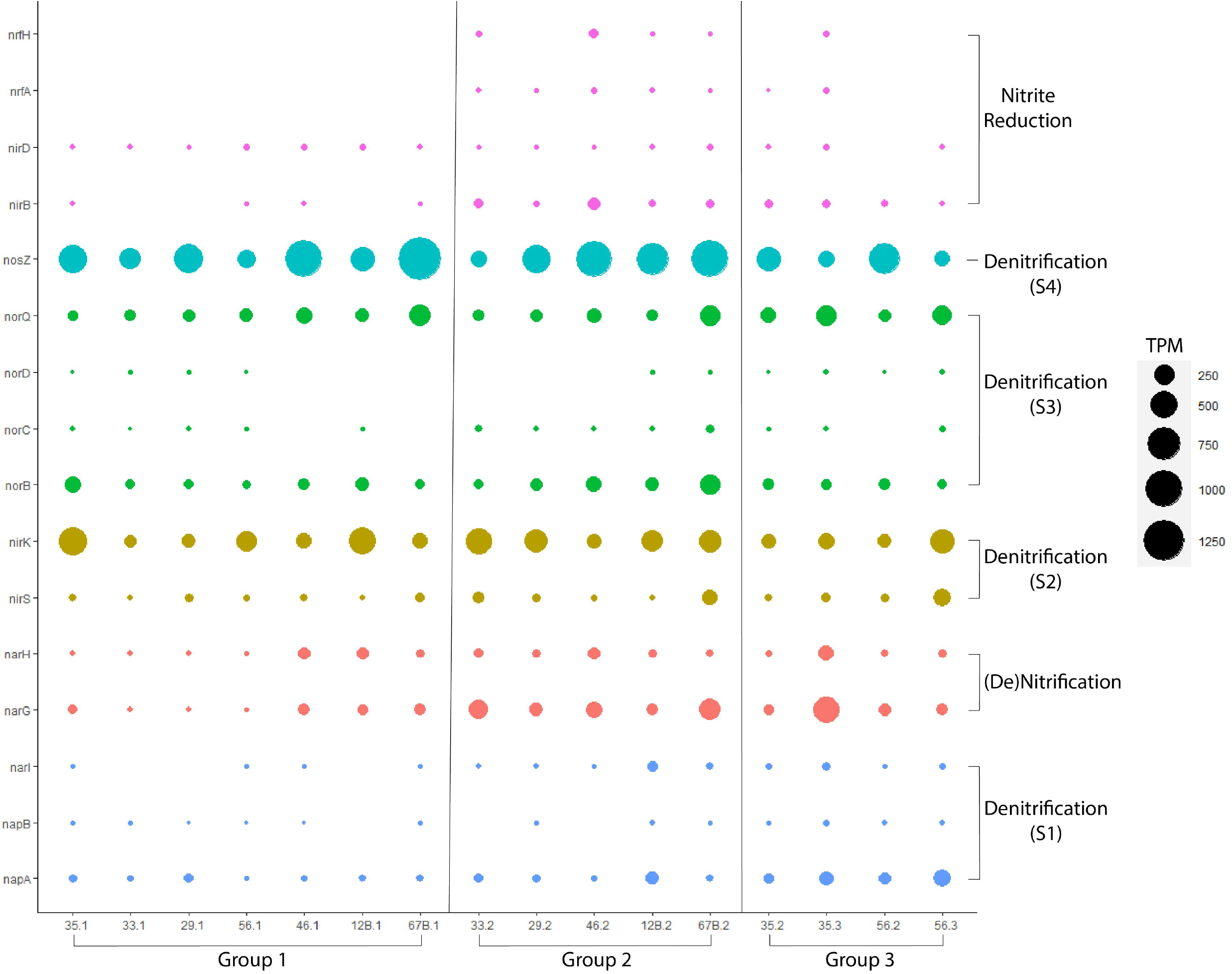
Abundance of nitrogen cycling genes in BSF metagenomes (values in transcripts per million - *TPM*). The *nifDHK*, *hao*, and *pmo/amoBC* genes were removed from this plot for readability as the abundance and processes are discussed in text and insignificant in the overall BSF nitrogen cycle.

**Figure 4.**
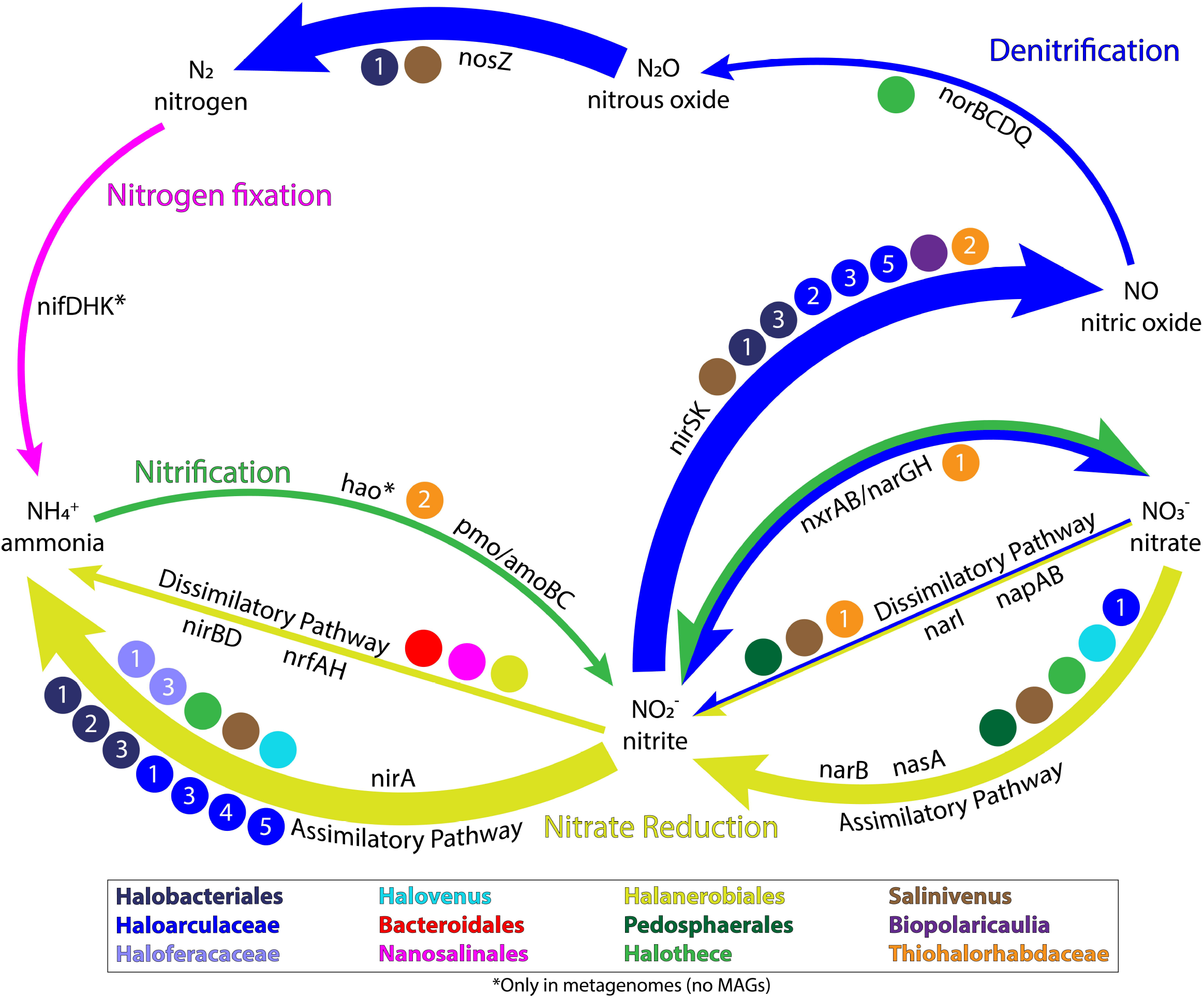
Overview of the BSF nitrogen cycle and associated genes and species. Colored circles represent MAGs containing at least 1 gene in the indicated pathway. Arrows are colored by process. The size of the arrow corresponds to the gene abundance in the sample metagenomes, where larger arrows indicate more abundant enzymes responsible for that reaction in the nitrogen cycle.

#### 3.4.2. Denitrification

Denitrification is a complex microbial process that can be broken into four steps carried out by one or multiple organisms. The first reaction in the process (nitrate to nitrite) overlaps with complete nitrate reduction, and the genes *narGH* are challenging to distinguish from *nxrAB* involved in the reverse reaction, nitrification (discussed above). Nitrate reductase catalyzes this step in denitrification, with two distinct dissimilatory types: *napA* represents a periplasmic version of the enzyme, and *narGH* represents the respiratory version (Sparacino-Watkins et al., 2014). These three nitrate reductase genes were present in all samples but never at high abundance when compared to other nitrogen-cycling enzymes (***Table S3***). While the respiratory version seems more abundant than the periplasmic type (under the assumption that all annotated narGH are nitrate reductase), both types appear most abundant in mineralogical group 3 samples (***Figure 3***). Only two MAGs contained genes involved in this first step. The *Salinivenus* MAG contained the periplasmic version of nitrate reductase (*napA*), and, as previously mentioned, the *Thiohalorhabdaceae*.1 MAG contained *narGHI*.

Two genes involved in the second step of denitrification (*nirSK*; nitrite reductase) are widespread and abundant across BSF (***Table S3***). While *nirK* has a fairly even and high abundance pattern between mineralogical groups, *nirS* is generally much less abundant with the highest counts in groups 2 and 3 and a particularly high abundance at sites 56 and 67B. No MAGs contained *nirS* annotations, but *nirK* was found in eight MAGs (*Salinivenus*, two *Halobacteriales*, three *Haloarculaceae*, *Bipolaricaulia*, and *Thiohalorhabdaceae*.2).

Genes encoding nitric oxide reductase (*norBC*) catalyze the third step of denitrification. These genes were only present at low abundances, but *norB* was widespread across BSF. A functional nitric oxide reductase is believed to require the accessory proteins norQD (Collman et al., 2008). In the BSF metagenomes, *norQ* was as widespread and abundant as *norB*. The only MAG including catalytic nitric oxide reductase genes was *Halothece*, which contained *norB* but is missing the *norQD* accessory protein genes. Three MAGs contained *norQ* genes, but not *norB*: *Halobacteriales*.1, *Haloarculaceae*.3, and *Thiohalorhabdaceae*.1 (***Figure 1***).

The last step in denitrification is carried out by nitrous oxide reductase, encoded by the gene *nosZ*. This gene is the most abundant nitrogen cycling gene at BSF and is relatively evenly distributed between mineralogical groups (***Figure 4***). Curiously, it has a roughly inverse abundance distribution with *nirK*; i.e., samples with a lower abundance of *nosZ* have a higher abundance of *nirK*. The *Salinivenus* and *Halobacteriales*.1 MAGs are the only two MAGs that encode *nosZ*.

#### 3.4.3. Dissimilatory Nitrate Reduction to Ammonia

The last step in dissimilatory nitrate reduction to ammonia is mediated by nitrite reductase. Genes encoding the large subunit of the NADH-dependent form of the enzyme (*nirB*) were less abundant or absent from metagenomes constructed from the surface halite (***Figure 3***). Genes encoding the cytochrome c form of the enzyme (*nrfAH*) were completely absent from all surface halite metagenomes (group 1) and all metagenomes constructed from samples collected at site 56. Generally, both *nir* and *nrf* genes are more abundant in mineralogical group 2 (upper gypsum). Three MAGs contained genes encoding nitrite reductase. The *Halanerobiales* MAG contained only the large subunit of the NADH-dependent form of the enzyme, while the *Nanosalinales* MAG only contained the small subunit (*nirD*). The Bacteroidales MAG contained both genes encoding the cytochrome c form of nitrite reductase (*nrfAH*).

### 3.5. Sulfur Cycle Genes

The gene encoding sulfide-quinone reductase (*sqr*) is among the most abundant genes in our BSF metagenomes (***Figure S2, Table S3, Figure 5***) and was present in seven MAGs (*Halobacteriales*.1, *Haloarculaceae*.3, *Haloarculaceae*.5, *Bacteroidales, Thiohalorhadbaceae*.1, *Thiohalorhadbaceae*.2, and *Methanohalobium*) and all metagenomes.

**Figure 5.**
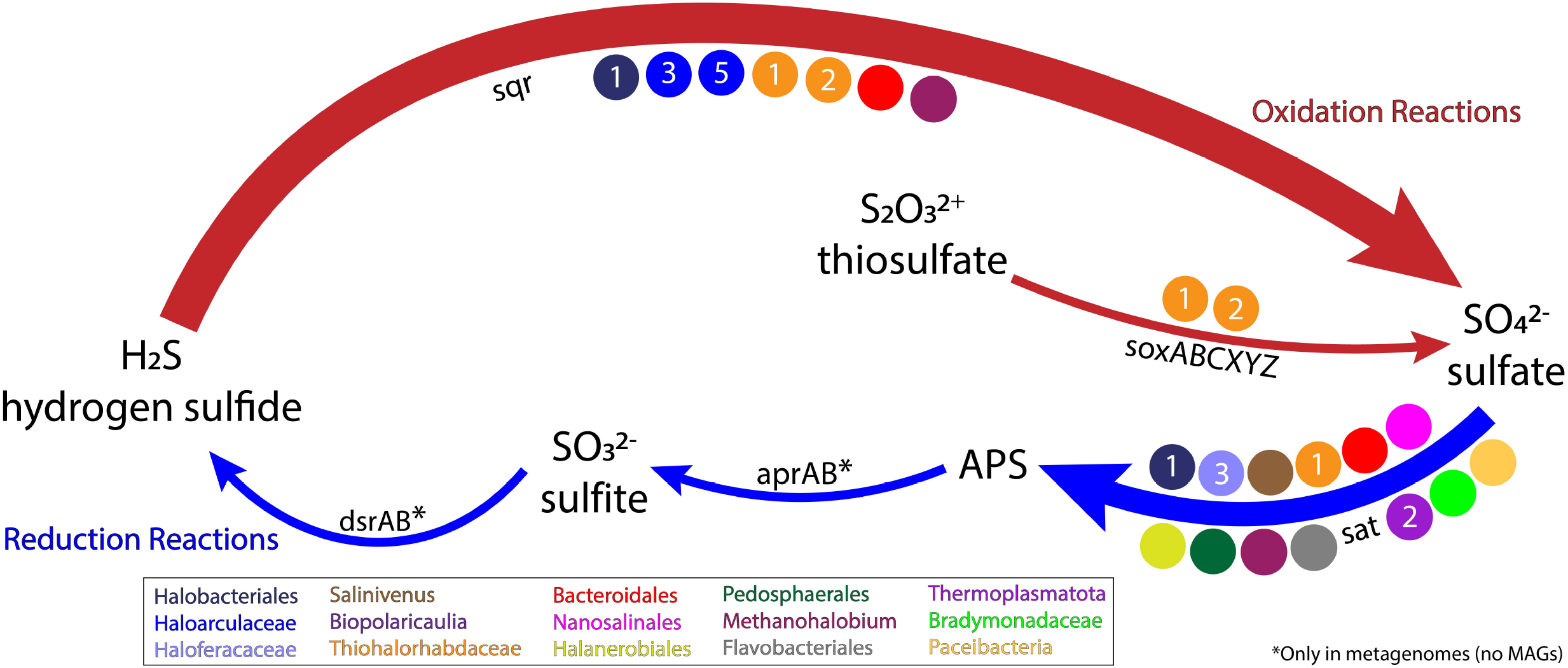
Overview of the BSF sulfur cycle and associated genes and species. Colored circles represent MAGs containing at least 1 gene in the indicated pathway. Arrows are colored by process. The size of the arrow corresponds to the gene abundance in the sample metagenomes, where larger arrows indicate more abundant enzymes responsible for that reaction in the sulfur cycle.

Other genes involved in the oxidation and reduction of sulfur species are widespread in BSF sediments but generally present at low abundances (***Table S3***). Genes encoding enzymes involved in dissimilatory sulfate reduction (*dsrAB*, *aprAB*) are found at most sites below the surface salt crust, but are most abundant in group 2 samples from sites 33 and 46 (***Figure 6***). Sulfate-reducing genes were only present in the surface halite at site 35. Unfortunately, no MAGs contained *dsrAB* genes, so we investigated individual *dsrAB* genes from metagenomes by querying them against the NCBI nr database (***Table S7***). A total of 11 *dsrA* and 14 *dsrB* genes had high sequence similarity to *Desulfovermiculus halophilus*, which were abundant in mineralogical group 2 in our previous survey of 16S rRNA genes (McGonigle et al., 2019). Other notable BLAST hits were to uncultured taxa sequenced from other salty environments, including the Great Salt Lake.

**Figure 6.**
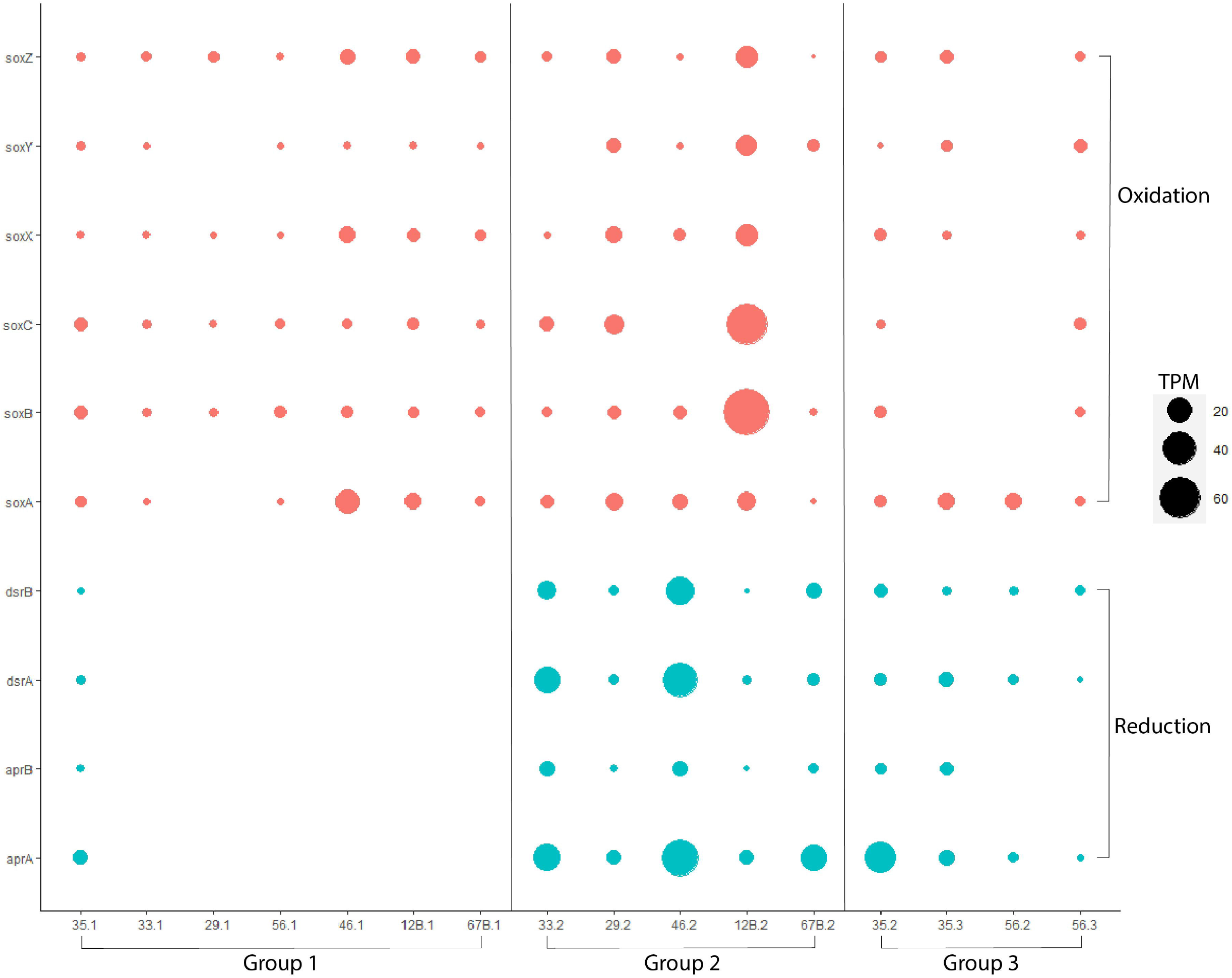
Abundance of sulfur cycling genes in BSF metagenomes (values in transcripts per million - *TPM*). The SQR gene is removed for readability; the high abundance of this gene compared to other BSF metabolic genes is shown in ***Figure S2***. The *sat* gene is also removed for readability as this gene is highly abundant and involved in sulfur assimilation and reduction.

Genes encoding the SOX system, associated with thiosulfate oxidation, were most abundant at site 12B, group 2, and were included in two MAGs (*Thiohalorhadbaceae*.1 and *Thiohalorhadbaceae*.2). However, these MAGs were missing the catalytic component (SoxABC) and only included predicted sequences for SoxYZ. Genes associated with sulfur oxidation and sulfur reduction had generally opposite abundance patterns; samples where SOX genes were more abundant had a lower abundance of genes involved in both dissimilatory and assimilatory sulfate reduction (***Figure 6***).

## 4. Discussion

### 4.1. BSF has an Active Microbial Ecosystem

Even though the Bonneville Salt Flats (BSF) appear to be utterly desolate at the surface, the layers of sediment just below the surface host a diverse, abundant, and active microbial ecosystem. The quantity of DNA extracted from these samples indicates the presence of dense microbial communities comparable to that of some soil communities (McGonigle et al., 2019). Our incubation experiments demonstrated the potential for both aerobic and anaerobic respiration in all sediment layers and mineralogical groups (***Table 1***), from the surface halite crust down to the lower gypsum layers 10-30 cm below the surface. Methanogenesis occurred during incubations of lower sediment layers. In addition, metagenomes reconstructed from the BSF sediments contained multiple carbon fixation pathways encoded by several taxa. Enzymes associated with many steps of the nitrogen and sulfur cycles are also widespread in the BSF sediment metagenomes. Below, we discuss some of these key results and their astrobiological implications.

### 4.2. Carbon Fixation Occurs at BSF via Multiple Pathways

As expected, the dominant carbon-fixation pathway at BSF is the reductive pentose phosphate (RPP) pathway. We would expect genes encoding the RuBisCO enzyme to be present in genomes of abundant cyanobacteria and algae (*Dunaliella salina*) that are known to be important primary producers in such salty ecosystems as BSF (Oren, 2014). Indeed, we found evidence for *Halothece*-like cyanobacterial species with a complete RPP pathway. The only other MAG containing a complete RPP pathway was classified as *Thiohalorhabdaceae* (***Figure 1***). An autotrophic lifestyle and the presence of RuBisCO have been previously described for the type genus for this family, *Thiohalorhabdus* (Tourova et al., 2010). Therefore, these taxa likely represent one of the primary producers at BSF.

Our previous survey of microbial diversity at BSF identified 16S rRNA gene sequences classified as putative acetogenic bacteria (*Acetothermia*). Indeed, the most complete Wood-Ljungdahl (WL) pathway was found in a MAG assigned to class *Bipolaricaulia* (within the phylum *Bipolaricaulota*, which was previously referred to as *Acetothermia* and candidate division OP1 (Hugenholtz et al., 1998; Hao et al., 2018). This MAG appears to account for most of the CODH genes detected in lower sediment layers and may be one of the primary producers in anaerobic zones of BSF. While the *Bipolaricaulia* MAG contained a near-complete WL pathway, it lacked genes encoding an acetate kinase and phosphate acetyltransferase, which are typically required for an acetogenic lifestyle (***Figure 1***). Both of these enzymes are also absent from 14 publicly available *Bipolaricaulota* MAGs that encode the WL pathway (Youssef et al., 2019). Additional analyses of these MAGs are required to assess their potential for autotrophic growth on CO_2_, heterotrophic homoacetogenic fermentation, and syntrophic growth via acetate oxidation, as proposed for other *Bipolaricaulota* (Youssef et al., 2019). Nevertheless, the presence of a near-complete WL pathway in the *Bipolaricaulia* MAG reported here suggests the possibility of acetogenesis as an important component of the BSF microbial ecosystem.

### 4.3. Methanogens at BSF Have a Diverse Suite of Potential Carbon Sources

The BSF metagenomes encode an incredible amount of metabolic flexibility for methanogenesis, with genes enabling the use of carbon dioxide, acetate, methanol, and mono-, di-, and tri-methylamines. Results from our incubation experiments and genomic analyses indicate that methanogens at BSF do not require hydrogen as an electron source. Methane production was generally higher in the absence of hydrogen (***Figure S1***), and sequences encoding hydrogenases associated with methanogenesis were absent.

All of the methanogenesis genes were included in one MAG classified as *Methanohalobium*. This genus is within the order *Methanosarcinales*, which are known to use a variety of substrates for methanogenesis (Thauer et al., 2008). The ability to use methylated carbon sources in methanogenesis is a known strategy to overcome competition for CO_2_ in saline environments (McGenity, 2010; McGenity and Sorokin, 2019). Furthermore, *Methanohalobium* belongs to a group of methanogens that tend to be more tolerant of oxidizing conditions (Lyu and Lu, 2018).

The production of ^13^CH_4_ from ^13^C-glucose in our aerobic incubations is evidence of heterotrophic carbon turnover, as glucose cannot be converted directly to methane by any known species (***Table 1***). We captured higher methane production under aerobic conditions, and no methane production occurred in anaerobic glucose incubations, even though these experiments captured conversion to ^13^CO_2_. This could indicate methanogens at BSF can utilize carbon released during aerobic respiration, but not the carbon molecules produced during anaerobic respiration. Curiously, acetate supported methane production in only one sample (site 12B, layer 2), suggesting that other organisms might outcompete methanogens for acetate at BSF. In support of the hypothesis that aerobic heterotrophy aids methanogenesis, *mcr* genes are only found in sediments just under the surface halite where limited oxygen is likely present in pockets, but not in either of our deeper metagenomes. Unfortunately, only two of our deepest layers were sequenced successfully, and additional sampling might show the presence of these genes in deeper sediments.

### 4.4. Haloarchaeal CO-Oxidizers at BSF Expand the Diversity of Known Carboxydovores

Interestingly, all but one of our MAGs (*Halanerobiales*) encoding aerobic CO dehydrogenase are classified as *Haloarchaea* (***Figure 1***). This group, whose CO-oxidizing ability has only recently been described, is also known to use the light-driven proton pump bacteriorhodopsin for supplemental energy (Myers and King, 2020). It is likely these BSF haloarchaea represent carboxydovores, which are unable to grow solely on CO, yet use atmospheric concentrations of CO as supplemental energy (Cordero et al., 2019). Multiple mechanisms (e.g. CO oxidation, light-harvesting complexes) to capture supplemental energy could be extremely beneficial for these heterotrophs living in an extreme environment that imposes additional costs for osmoregulation.

Known CO-oxidizing haloarchaea have previously been isolated from salty environments, including BSF (Myers and King, 2020). Our haloarchaea MAGs are distinct from the three BSF isolates (*Halobacterium bonnevillei, Halobaculum saliterrae*, and *Halovenus carboxidivorans*) that have been previously reported, and they are most likely new species or genera of haloarchaea (average nucleotide identities <80%). Furthermore, our results indicate that these novel CO-oxidizing haloarchaea encode form I aerobic CODH. These MAGs encode additional genes annotated as aerobic CODH but lack significant sequence similarity with any characterized forms of CODH. Further work should explore whether these sequences represent members of form II CODH, a distinct form of CODH, or another enzyme entirely.

The only non-archaeal MAG (*Halanerobiales*) containing aerobic CODH genes is missing *coxL*, which encodes the catalytic subunit of the enzyme. A functional enzyme might still be present in this species because the *coxL* gene family is diverse and is known to be frequently mis-annotated (King and Weber, 2007). Nevertheless, the absence of bacterial *coxL* in these metagenomes suggests that CO oxidation in BSF may be carried out only by archaea.

### 4.5. Denitrification is the Center of the BSF Nitrogen Cycle

Denitrification is the most widespread and abundant portion of the BSF nitrogen cycle (***Figure 4***). The taxonomy of MAGs containing denitrification genes have been previously described as highly abundant organisms at BSF through 16S rRNA analyses (eg. *Salinivenus, Halobacteriales, Nanosalinales, Halanerobiales*, and *Thiohalorhabdaceae*) (Lynch et al., 2019b; McGonigle et al., 2019). While numerous MAGs contained genes for denitrification, none contained genes for all four steps in the process. Two MAGs (*Salinivenus* and *Halobacteriales*.1,) encode multiple steps of denitrification. Curiously, the *Salinivenus* MAG has three genes involved in the process: *napA* (step 1), *nirK* (step 2), and *nosZ* (step 4). Atypical *nosZ* genes, encoding a functional complex, are previously described in the type species for the *Salinibacteraceae* family, *Salinibacter ruber (S. ruber*) (Sanford et al., 2012). However, culture work shows *S. ruber* is unable to grow with nitrate as an electron acceptor, and denitrifying species in this family have not been identified (Bagheri et al., 2019).

Interestingly, we found *nosZ* to be a highly abundant gene (***Figure S2***). This is in apparent contrast to other environmental studies, which have shown species containing *nosZ* are often less abundant than microbes with genes encoding the preceding reductases in the denitrification pathway, such as *nirK* (Jones et al., 2013). It is to be noted that while *nosZ* is more abundant overall, the BSF samples with the highest abundances of *nirK* seem to have a lower abundance of *nosZ* (***Table S3***).

Denitrifying haloarchaea have been described, but it is unclear if they participate in partial or complete denitrification (Torregrosa-Crespo et al., 2018). Our MAGs suggest the BSF haloarchaea are involved in partial denitrification, particularly in reducing nitrite to nitric oxide (*nirSK*). A functional NirK enzyme has been extracted from a cultured member of *Halobacteriales (Haloferax mediterranei*) (Esclapez et al., 2013). Interestingly, the same haloarchaea MAGs encode aerobic CODH, and denitrification linked with CO oxidation has been previously reported for bacteria (King, 2006).

The abundance of genes associated with denitrification contrasts sharply with the sparse distribution of nitrogen fixation genes. Nitrogenase-encoding sequences were relatively rare or absent in all BSF metagenomes. For example, *Halothece* cyanobacteria are commonly known as nitrogen fixers (Li et al., 2019). Yet, the only nitrogen-associated gene in our *Halothece* MAG was *norB*, which catalyzes the reduction of nitric oxide during denitrification. The few nitrogenase sequences in BSF had similarities with those encoded by photoautotrophic and/or sulfur-cycling bacteria. Nitrogenase sequences were mostly found in lower layers of the sediment, suggesting that further sampling could yield additional nitrogenase genes encoded by organisms inhabiting anoxic zones in deeper sediments.

### 4.6 BSF Contains a Dynamic Sulfur Cycle with Surprising Players

Sulfide oxidation is abundant and widespread across BSF, including the surface salt crust. This process may occur through aerobic or anaerobic pathways, using alternative electron acceptors such as those used in denitrification (Shao et al., 2010). The gene encoding sulfide-quinone reductase (SQR) is one of the most abundant genes in the BSF metagenomes. This enzyme can be involved in energy conservation, by linking sulfide oxidation to ATP production, or in sulfide detoxification, where the oxidation of sulfide only serves to prevent damage to the cell (Schütz et al., 1999; Pham et al., 2008; Marcia et al., 2009; van der Meer et al., 2010).

We identified diverse taxa encoding SQR, including *Thiohalorhadbaceae, Halobacteriales*, *Haloarculaceae*, *Bacteroidales*, and *Methanohalobium*. The SQR gene is not typically associated with haloarchaea or methanogens, but it has been identified in a *Bacteroidales*-like species through metagenomic studies of a Siberian soda lake (Vavourakis et al., 2019). Interestingly, the *Bacteroidales* MAG has the genomic capacity to couple sulfide oxidation to nitrite reduction, thereby linking the two macronutrient cycles. Supporting this, the genome lacks a cytochrome c oxidase, along with other genes typically associated with aerobic metabolism. It does contain *nrfA*, which has been shown to use nitrite as a respiratory electron acceptor in *Escherichia coli* (Clarke et al., 2008).

Our results indicate *Thiohalorhadbaceae*-like species are important players in the BSF sulfur cycle. They likely play a role in sulfite (and possibly sulfate) assimilation for amino acid synthesis and sulfide oxidation for energy conservation (via SQR). Even though the type species for this family, *Thiohalorhabdus denitrificans*, is known to oxidize thiosulfate (Sorokin et al., 2008), neither of our *Thiohalorhadbaceae* MAGs contain catalytic *sox* genes. While other *sox* genes are present in the metagenomes, the *Thiohalorhadbaceae* MAGs only contain *soxYZ* genes, recently shown to encode a protein that acts as a carrier in the multistep process of thiosulfate oxidation (Grabarczyk and Berks, 2017). This is consistent with the absence of *soxB* genes in salt marsh sediments where *Thiohalorhabdus*-like species were also described (Thomas et al., 2014). Not much is known about this largely uncultured family, but these results suggest that some members of *Thiohalorhadbaceae* are unable to oxidize thiosulfate or else do so with novel enzymes.

The inclusion of SQR in our *Methanohalobium* MAG is consistent with its presence in the genome of *Methanolobus* species, who belong to the same family of methanogens (*Methanosarcinaceae*) and are known to be capable of sulfide oxidation (Stetter and Gaag, 1983). Metagenomic sequences predicted to encode SQR have also been associated with the archaeal methanotrophs ANME-1, who have been proposed to be capable of using SQR in the reverse direction, to produce sulfide instead of or in addition to oxidizing sulfide (Milucka et al., 2012; Vigneron et al., 2019). Environmental surveys indicate that SQR sequences are diverse and mostly uncharacterized, indicating a potentially widespread but unknown role in sulfur-rich environments (Pham et al., 2008).

## 5. Conclusions

This metagenomic study has produced an inventory of potential energy sources and metabolic pathways that are feasible in the hypersaline sediments of BSF. Furthermore, laboratory incubation experiments confirmed that BSF microbial communities are capable of aerobic and anaerobic respiration and methanogenesis from multiple carbon sources. Analyses of individual genomes highlighted reduced sulfur and nitrate as a likely redox couple for multiple species of BSF archaea and bacteria. In particular, genomes of haloarchaea (*Halobacteriales* and *Haloarculaceae*) encode the potential for sulfide oxidation, CO oxidation, and denitrification, suggesting that they may play a central role in the cycling of carbon, nitrogen, and sulfur in BSF.

As an ecosystem that has persisted in a shallow subsurface, mineral-rich environment during thousands of years of desiccation, BSF is an important natural laboratory for investigating the past, present, and future potential for life on Mars. The diverse microbial communities of BSF are capable of maintaining an active and sustainable ecosystem, even though the surface salt crust appears to be completely devoid of life. Similarly, Mars contains expansive evaporite deposits, and recent missions have begun to explore their underlying shallow subsurface environments (Ehlmann et al., 2011; Farley et al., 2020). If hypersaline sediments on Mars are not habitable, then further study of BSF and similar environments on Earth will be important for understanding which requirements for life are lacking on Mars.

## Acknowledgments

The authors gratefully acknowledge Betsy Kleba and Emily Dart for their critical roles in collecting samples for this study.

---

***Table S1*** Overview of the BSF metagenomic assemblies used in this study, including quality measurements, number of contigs, and number of predicted proteins per assembly.

***Table S2*** Summary table with completion and contamination of the BSF MAGs used in this study (as determined through checkM).

***Table S3*** Functional read abundance for genes of interest in each assembly, normalized as transcripts per million (*TPM*).

***Table S4*** Genomic organization of aerobic CODH (*cox*) genes in the BSF MAGs and their associated enzyme form.

***Table S5*** Table of average nucleotide identity (ANI) results for the haloarchaeal MAGs compared against reference genomes of cultured haloarchaea carboxydovores.

***Table S6*** BLAST results for nitrogenase genes in the BSF metagenomes. Best BLAST hit reported here was determined by highest bit score and lowest e-value.

***Table S7*** BLAST results for *dsrAB* genes in the BSF metagenomes. Best BLAST hit reported here was determined by highest bit score and lowest e-value.

***Figure S1*** Recovered ^13^CH_4_ from the headspace of ^13^CO_2_ incubations for 4 sites at the BSF: 12B, 33, 41, and 56. Incubations were done aerobically and anaerobically with and without the addition of hydrogen gas, in triplicate. The x-axis represents layers in the sediment column at each site. Anaerobic incubations for layer 4 at site 56 failed.

***Figure S2*** Heatmap of normalized gene abundance (as scaled *TPM* values) for genes and complexes of interest at the BSF. Names on the left represent KEGG pathways associated with these genes, but overlaps with other functions are possible.

## References

Arkin, A.P., Cottingham, R.W., Henry, C.S., Harris, N.L., Stevens, R.L., Maslov, S., Dehal, P., Ware, D., Perez, F., Canon, S., Sneddon, M.W., Henderson, M.L., Riehl, W.J., Murphy-Olson, D., Chan, S.Y., Kamimura, R.T., Kumari, S., Drake, M.M., Brettin, T.S., Glass, E.M., Chivian, D., Gunter, D., Weston, D.J., Allen, B.H., Baumohl, J., Best, A.A., Bowen, B., Brenner, S.E., Bun, C.C., Chandonia, J.M., Chia, J.M., Colasanti, R., Conrad, N., Davis, J.J., Davison, B.H., Dejongh, M., Devoid, S., Dietrich, E., Dubchak, I., Edirisinghe, J.N., Fang, G., Faria, J.P., Frybarger, P.M., Gerlach, W., Gerstein, M., Greiner, A., Gurtowski, J., Haun, H.L., He, F., Jain, R., Joachimiak, M.P., Keegan, K.P., Kondo, S., Kumar, V., Land, M.L., Meyer, F., Mills, M., Novichkov, P.S., Oh, T., Olsen, G.J., Olson, R., Parrello, B., Pasternak, S., Pearson, E., Poon, S.S., Price, G.A., Ramakrishnan, S., Ranjan, P., Ronald, P.C., Schatz, M.C., Seaver, S.M.D., Shukla, M., Sutormin, R.A., Syed, M.H., Thomason, J., Tintle, N.L., Wang, D., Xia, F., Yoo, H., Yoo, S., Yu, D. (2018) KBase: The United States department of energy systems biology knowledgebase. Nature Biotechnology, 36, 566–569. https://doi.org/10.1038/nbt.4163

Azua-Bustos, A., Caro-Lara, L., Vicuña, R. (2015) Discovery and microbial content of the driest site of the hyperarid Atacama Desert, Chile. Environmental Microbiology Reports, 7, 388–394. https://doi.org/10.1111/1758-2229.12261

Bagheri, M., Marashi, S.A., Amoozegar, M.A. (2019) A genome-scale metabolic network reconstruction of extremely halophilic bacterium Salinibacter ruber. PLoS ONE, 14, 1–17. https://doi.org/10.1371/journal.pone.0216336

Barbieri, R., Stivaletta, N. (2011) Continental evaporites and the search for evidence of life on Mars. Geological Journal, 46, 513–524. https://doi.org/10.1002/gj.1326

Bowen, B.B., Kipnis, E.L., Raming, L.W. (2017) Temporal dynamics of flooding, evaporation, and desiccation cycles and observations of salt crust area change at the Bonneville Salt Flats, Utah. Geomorphology, 299, 1–11. https://doi.org/10.1016/j.geomorph.2017.09.036

Brazelton, W.J., Ludwig, K.A., Sogin, M.L., Andreishcheva, E.N., Kelley, D.S., Shen, C.C., Edwards, R.L., Baross, J.A. (2010) Archaea and bacteria with surprising microdiversity show shifts in dominance over 1,000-year time scales in hydrothermal chimneys. Proceedings of the National Academy of Sciences of the United States of America, 107, 1612–1617. https://doi.org/10.1073/pnas.0905369107

Bryant, R.G., Rainey, M.P. (2002) Investigation of flood inundation on playas within the Zone of Chotts, using a time-series of AVHRR. Remote Sensing of Environment, 82, 360–375. https://doi.org/10.1016/S0034-4257(02)00053-6

Buchfink, B., Reuter, K., Drost, H.G. (2021) Sensitive protein alignments at tree-of-life scale using DIAMOND. Nature Methods, 18, 366–368. https://doi.org/10.1038/s41592-021-01101-x

Bushnell, B., Rood, J., Singer, E. (2017) BBMerge – Accurate paired shotgun read merging via overlap. PLoS ONE, 12, 1–15. https://doi.org/10.1371/journal.pone.0185056

Chaumeil, P.A., Mussig, A.J., Hugenholtz, P., Parks, D.H. (2020) GTDB-Tk: A toolkit to classify genomes with the genome taxonomy database. Bioinformatics, 36, 1925–1927. https://doi.org/10.1093/bioinformatics/btz848

Clarke, T.A., Mills, P.C., Poock, S.R., Butt, J.N., Cheesman, M.R., Cole, J.A., Hinton, J.C.D., Hemmings, A.M., Kemp, G., Söderberg, C.A.G., Spiro, S., Van Wonderen, J., Richardson, D.J. (2008) Escherichia coli Cytochrome c Nitrite Reductase NrfA. Methods in Enzymology, 437, 63–77. https://doi.org/10.1016/S0076-6879(07)37004-3

Collman, J.P., Yang, Y., Dey, A., Decréau, R.A., Ghosh, S., Ohta, T., Solomon, E.I. (2008) A functional nitric oxide reductase model. Proceedings of the National Academy of Sciences of the United States of America, 105, 15660–15665. https://doi.org/10.1073/pnas.0808606105

Cordero, P.R.F., Bayly, K., Man Leung, P., Huang, C., Islam, Z.F., Schittenhelm, R.B., King, G.M., Greening, C. (2019) Atmospheric carbon monoxide oxidation is a widespread mechanism supporting microbial survival. ISME Journal, 13, 2868–2881. https://doi.org/10.1038/s41396-019-0479-8

Dyall-Smith, M. (2009) The halohandbook: protocols for halobacterial genetics, version 7.2

Ehlmann, B.L., Mustard, J.F., Murchie, S.L., Bibring, J.P., Meunier, A., Fraeman, A.A., Langevin, Y. (2011) Subsurface water and clay mineral formation during the early history of Mars. Nature, 479, 53–60. https://doi.org/10.1038/nature10582

Esclapez, J., Zafrilla, B., Martínez-Espinosa, R.M., Bonete, M.J. (2013) Cu-NirK from Haloferax mediterranei as an example of metalloprotein maturation and exportation via Tat system. Biochimica et Biophysica Acta - Proteins and Proteomics, 1834, 1003–1009. https://doi.org/10.1016/j.bbapap.2013.03.002

Farley, K.A., Williford, K.H., Stack, K.M., Bhartia, R., Chen, A., de la Torre, M., Hand, K., Goreva, Y., Herd, C.D.K., Hueso, R., Liu, Y., Maki, J.N., Martinez, G., Moeller, R.C., Nelessen, A., Newman, C.E., Nunes, D., Ponce, A., Spanovich, N., Willis, P.A., Beegle, L.W., Bell, J.F., Brown, A.J., Hamran, S.-E., Hurowitz, J.A., Maurice, S., Paige, D.A., Rodriguez-Manfredi, J.A., Schulte, M., Wiens, R.C. (2020) Mars 2020 Mission Overview. Space Science Reviews 2020 216:8, 216, 1–41. https://doi.org/10.1007/S11214-020-00762-Y

Goudge, T.A., Mustard, J.F., Head, J.W., Fassett, C.I., Wiseman, S.M. (2015) Assessing the mineralogy of the watershed and fan deposits of the Jezero crater paleolake system, Mars. Journal of Geophysical Research: Planets, 120, 775–808. https://doi.org/10.1038/175238c0

Grabarczyk, D.B., Berks, B.C. (2017) Intermediates in the Sox sulfur oxidation pathway are bound to a sulfane conjugate of the carrier protein SoxYZ. PLoS ONE, 12, 1–15. https://doi.org/10.1371/journal.pone.0173395

Graham, E.D., Heidelberg, J.F., Tully, B.J. (2017) Binsanity: Unsupervised clustering of environmental microbial assemblies using coverage and affinity propagation. PeerJ, 2017, 1–19. https://doi.org/10.7717/peerj.3035

Hao, L., McIlroy, S.J., Kirkegaard, R.H., Karst, S.M., Fernando, W.E.Y., Aslan, H., Meyer, R.L., Albertsen, M., Nielsen, P.H., Dueholm, M.S. (2018) Novel prosthecate bacteria from the candidate phylum Acetothermia. ISME Journal, 12, 2225–2237. https://doi.org/10.1038/s41396-018-0187-9

Horgan, B.H.N., Anderson, R.B., Dromart, G., Amador, E.S., Rice, M.S. (2020) The mineral diversity of Jezero crater: Evidence for possible lacustrine carbonates on Mars. Icarus, 339, 113526. https://doi.org/10.1016/j.icarus.2019.113526

Hugenholtz, P., Pitulle, C., Hershberger, K.L., Pace, N.R. (1998) Novel division level bacterial diversity in a Yellowstone hot spring. Journal of Bacteriology, 180, 366–376. https://doi.org/10.1128/jb.180.2.366-376.1998

Hydrologic Basin, Death Valley, California (1966, Washington D.C.). 1966. Ed. Hunt, C.B., Robinson, T., Bowles, W.A., Washburn, A. Washington D.C.,

Hyatt, D., Chen, G.-L., LoCascio, P.F., Land, M.L., Larimer, F.W., Hauser, L.J. (2010) Prodigal: prokaryotic gene recognition and translation initiation site identification. BMC Bioinformatics, 11, 119. https://doi.org/10.1186/1471-2105-11-119

Jones, C.M., Graf, D.R.H., Bru, D., Philippot, L., Hallin, S. (2013) The unaccounted yet abundant nitrous oxide-reducing microbial community: A potential nitrous oxide sink. ISME Journal, 7, 417–426. https://doi.org/10.1038/ismej.2012.125

Kampf, S.K., Tyler, S.W. (2006) Spatial characterization of land surface energy fluxes and uncertainty estimation at the Salar de Atacama, Northern Chile. Advances in Water Resources, 29, 336–354. https://doi.org/10.1016/j.advwatres.2005.02.017

King, G.M. (2006) Nitrate-dependent anaerobic carbon monoxide oxidation by aerobic CO-oxidizing bacteria. FEMS Microbiology Ecology, 56, 1–7. https://doi.org/10.1111/j.1574-6941.2006.00065.x

King, G.M. (2015) Carbon monoxide as a metabolic energy source for extremely halophilic microbes: Implications for microbial activity in Mars regolith. Proceedings of the National Academy of Sciences, 112, 4465–4470. https://doi.org/10.1073/pnas.1424989112

King, G.M., Weber, C.F. (2007) Distribution, diversity and ecology of aerobic CO-oxidizing bacteria. Nature Reviews Microbiology, 5, 107–118. https://doi.org/10.1038/nrmicro1595

Lacap-Bugler, D.C., Lee, K.K., Archer, S., Gillman, L.N., Lau, M.C.Y., Leuzinger, S., Lee, C.K., Maki, T., McKay, C.P., Perrott, J.K., de los Rios-Murillo, A., Warren-Rhodes, K.A., Hopkins, D.W., Pointing, S.B. (2017) Global diversity of desert hypolithic cyanobacteria. Frontiers in Microbiology, 8, 1–13. https://doi.org/10.3389/fmicb.2017.00867

Langmead, B., Salzberg, S.L. (2012) Fast gapped-read alignment with Bowtie 2. Nature Methods, 9, 357–359. https://doi.org/10.1038/nmeth.1923

Li, D., Luo, R., Liu, C.M., Leung, C.M., Ting, H.F., Sadakane, K., Yamashita, H., Lam, T.W. (2016) MEGAHIT v1.0: A fast and scalable metagenome assembler driven by advanced methodologies and community practices. Methods, 102, 3–11. https://doi.org/10.1016/j.ymeth.2016.02.020

Li, Y., Cha, Q.Q., Dang, Y.R., Chen, X.L., Wang, M., McMinn, A., Espina, G., Zhang, Y.Z., Blamey, J.M., Qin, Q.L. (2019) Reconstruction of the Functional Ecosystem in the High Light, Low Temperature Union Glacier Region, Antarctica. Frontiers in Microbiology, 10, 1–14. https://doi.org/10.3389/fmicb.2019.02408

Lynch, K.L. 2015. A Geobiological Investigation of the Hypersaline Sediments of Pilot Valley, Utah: A Terrestrial Analog to Ancient Lake Basins on Mars. s.l.,

Lynch, K.L., Andrew Jackson, W., Rey, K., Spear, J.R., Rosenzweig, F., Munakata-Marr, J. (2019a) Evidence for Biotic Perchlorate Reduction in Naturally Perchlorate-Rich Sediments of Pilot Valley Basin, Utah. Astrobiology, 19, 629–641. https://doi.org/10.1089/ast.2018.1864

Lynch, K.L., Rey, K.A., Bond, R.J., Biddle, J.F., Spear, J.R., Rosenzweig, F., Munakata-Marr, J. (2019b) Discrete Community Assemblages Within Hypersaline Paleolake Sediments of Pilot Valley, Utah. bioRxiv, no. May. https://doi.org/10.1101/634642

Lyu, Z., Lu, Y. (2018) Metabolic shift at the class level sheds light on adaptation of methanogens to oxidative environments. ISME Journal, 12, 411–423. https://doi.org/10.1038/ismej.2017.173

Marcia, M., Ermler, U., Peng, G., Michel, H. (2009) The structure of Aquifex aeolicus sulfide:quinone oxidoreductase, a basis to understand sulfide detoxification and respiration. Proceedings of the National Academy of Sciences of the United States of America, 106, 9625–9630. https://doi.org/10.1073/pnas.0904165106

McGenity, T.J. (2010) Methanogens and Methanogenesis in Hypersaline Environments BT - Handbook of Hydrocarbon and Lipid Microbiology. In: Timmis, K.N. (ed.). Berlin, Heidelberg: Springer Berlin Heidelberg: pp. 665–680.

McGenity, T.J., Sorokin, D.Y. (2019) Methanogens and Methanogenesis in Hypersaline Environments BT - Biogenesis of Hydrocarbons. In: Stams, A.J.M., and Sousa, D.Z. (eds.). Cham: Springer International Publishing: pp. 283–309.

McGonigle, J.M., Bernau, J.A., Bowen, B.B., Brazelton, W. (2019) Robust archaeal and bacterial communities inhabit shallow subsurface sediments of the Bonneville Salt Flats. mSphere, 4, e00378–19. https://doi.org/10.1101/553032

van der Meer, M.T.J., Klatt, C.G., Wood, J., Bryant, D.A., Bateson, M.M., Lammerts, L., Schouten, S., Sinninghe Damste, J.S., Madigan, M.T., Ward, D.M. (2010) Cultivation and genomic, nutritional, and lipid biomarker characterization of Roseiflexus strains closely related to predominant in situ populations inhabiting yellowstone hot spring microbial mats. Journal of Bacteriology, 192, 3033–3042. https://doi.org/10.1128/JB.01610-09

Milucka, J., Ferdelman, T.G., Polerecky, L., Franzke, D., Wegener, G., Schmid, M., Lieberwirth, I., Wagner, M., Widdel, F., Kuypers, M.M.M. (2012) Zero-valent sulphur is a key intermediate in marine methane oxidation. Nature, 491, 541–546. https://doi.org/10.1038/nature11656

Myers, M.R., King, G.M. (2020) Halobacterium bonnevillei sp. Nov., halobaculum saliterrae sp. nov. and halovenus carboxidivorans sp. nov., three novel carbon monoxide-oxidizing halobacteria from saline crusts and soils. International Journal of Systematic and Evolutionary Microbiology, 70, 4261–4268. https://doi.org/10.1099/ijsem.0.004282

Oren, A. (2014) The ecology of Dunaliella in high-salt environments. Journal of Biological Research (Greece), 21, 1–8.https://doi.org/10.1186/s40709-014-0023-y

Oviatt, C.G. (2015) Chronology of Lake Bonneville, 30,000 to 10,000 yr BP. Quaternary Science Reviews, 110, 166–171

Parks, D.H., Imelfort, M., Skennerton, C.T., Hugenholtz, P., Tyson, G.W. (2015) CheckM: Assessing the quality of microbial genomes recovered from isolates, single cells, and metagenomes. Genome Research, 25, 1043–1055. https://doi.org/10.1101/gr.186072.114

Pham, V.H., Yong, J.J., Park, S.J., Yoon, D.N., Chung, W.H., Rhee, S.K. (2008) Molecular analysis of the diversity of the sulfide: Quinone reductase (sqr) gene in sediment environments. Microbiology, 154, 3112–3121. https://doi.org/10.1099/mic.0.2008/018580-0

Quinlan, A.R., Kindlon, N. (2019) bedtools: A powerful toolset for genome arithmetic. 2019

Rohland, N., Reich, D. (2012) Cost-effective, high-throughput DNA sequencing libraries for multiplexed target capture. Genome Research, 22, 939–946. https://doi.org/10.1101/gr.128124.111

Sanford, R.A., Wagner, D.D., Wu, Q., Chee-Sanford, J.C., Thomas, S.H., Cruz-García, C., Rodríguez, G., Massol-Deyá, A., Krishnani, K.K., Ritalahti, K.M., Nissen, S., Konstantinidis, K.T., Löffler, F.E. (2012) Unexpected nondenitrifier nitrous oxide reductase gene diversity and abundance in soils. Proceedings of the National Academy of Sciences of the United States of America, 109, 19709–19714. https://doi.org/10.1073/pnas.1211238109

Schulz, S., Horovitz, M., Rausch, R., Michelsen, N., Mallast, U., Köhne, M., Siebert, C., Schüth, C., Al-Saud, M., Merz, R. (2015) Groundwater evaporation from salt pans: Examples from the eastern Arabian Peninsula. Journal of Hydrology, 531, 792–801. https://doi.org/10.1016/j.jhydrol.2015.10.048

Schütz, M., Maldener, I., Griesbeck, C., Hauska, G. (1999) Sulfide-quinone reductase from Rhodobacter capsulatus: Requirement for growth, periplasmic localization, and extension of gene sequence analysis. Journal of Bacteriology, 181, 6516–6523. https://doi.org/10.1128/jb.181.20.6516-6523.1999

Shao, M.F., Zhang, T., Fang, H.H.P. (2010) Sulfur-driven autotrophic denitrification: Diversity, biochemistry, and engineering applications. Applied Microbiology and Biotechnology, 88, 1027–1042. https://doi.org/10.1007/s00253-010-2847-1

Sorokin, D.Y., Tourova, T.P., Galinski, E.A., Muyzer, G., Kuenen, J.G. (2008) Thiohalorhabdus denitrificans gen. nov., sp. nov., an extremely halophilic, sulfur-oxidizing, deep-lineage gammaproteobacterium from hypersaline habitats. International Journal of Systematic and Evolutionary Microbiology, 58, 2890–2897. https://doi.org/10.1099/ijs.0.2008/000166-0

Sparacino-Watkins, C., Stolz, J.F., Basu, P. (2014) Nitrate and periplasmic nitrate reductases. Chemical Society reviews, 43, 676–706. https://doi.org/10.1039/c3cs60249d.Nitrate

Stetter, K.O., Gaag, G. (1983) Reduction of molecular sulphur by methanogenic bacteria. Nature, 305, 309–311. https://doi.org/10.1038/305309a0

Thauer, R.K., Kaster, A.K., Goenrich, M., Schick, M., Hiromoto, T., Shima, S. (2010) Hydrogenases from methanogenic archaea, nickel, a novel cofactor, and H2 storage. Annual Review of Biochemistry, 79, 507–536. https://doi.org/10.1146/annurev.biochem.030508.152103

Thauer, R.K., Kaster, A.K., Seedorf, H., Buckel, W., Hedderich, R. (2008) Methanogenic archaea: Ecologically relevant differences in energy conservation. Nature Reviews Microbiology, 6, 579–591. https://doi.org/10.1038/nrmicro1931

Thomas, F., Giblin, A.E., Cardon, Z.G., Sievert, S.M. (2014) Rhizosphere heterogeneity shapes abundance and activity of sulfur-oxidizing bacteria in vegetated salt marsh sediments. Frontiers in Microbiology, 5, 1–14. https://doi.org/10.3389/fmicb.2014.00309

Torregrosa-Crespo, J., Bergaust, L., Pire, C., Martínez-Espinosa, R.M. (2018) Denitrifying haloarchaea: Sources and sinks of nitrogenous gases. FEMS Microbiology Letters, 365, 1–6. https://doi.org/10.1093/femsle/fnx270

Tourova, T.P., Kovaleva, O.L., Sorokin, D.Y., Muyzer, G. (2010) Ribulose-1,5-bisphosphate carboxylase/oxygenase genes as a functional marker for chemolithoautotrophic halophilic sulfur-oxidizing bacteria in hypersaline habitats. Microbiology, 156, 2016–2025. https://doi.org/10.1099/mic.0.034603-0

Vavourakis, C.D., Mehrshad, M., Balkema, C., Van Hall, R., Andrei, A.Ş., Ghai, R., Sorokin, D.Y., Muyzer, G. (2019) Metagenomes and metatranscriptomes shed new light on the microbial-mediated sulfur cycle in a Siberian soda lake. BMC Biology, 17, 1–20. https://doi.org/10.1186/s12915-019-0688-7

Vigneron, A., Alsop, E.B., Cruaud, P., Philibert, G., King, B., Baksmaty, L., Lavallee, D., Lomans, B.P., Eloe-Fadrosh, E., Kyrpides, N.C., Head, I.M., Tsesmetzis, N. (2019) Contrasting Pathways for Anaerobic Methane Oxidation in Gulf of Mexico Cold Seep Sediments. mSystems, 4, 1–17. https://doi.org/10.1128/msystems.00091-18

Wierzchos, J., Rodríguez-Ochoa, R., Ascaso, C., Pueyo, J., Buhler, P., Davila, A., Artieda, O., Davila, A., Wierzchos, J., Buhler, P., Rodríguez-Ochoa, R., Pueyo, J., Ascaso, C. (2015) Surface evolution of salt-encrusted playas under extreme and continued dryness. Earth Surface Processes and Landforms, 40, 1939–1950. https://doi.org/10.1002/esp.3771

Youssef, N.H., Farag, I.F., Rudy, S., Mulliner, A., Walker, K., Caldwell, F., Miller, M., Hoff, W., Elshahed, M. (2019) The Wood–Ljungdahl pathway as a key component of metabolic versatility in candidate phylum Bipolaricaulota (Acetothermia, OP1). Environmental Microbiology Reports, 11, 538–547. https://doi.org/10.1111/1758-2229.12753

