## Supplementary figures and images for "Metabolic Potential of Microbial Communities in the Hypersaline Sediments of the Bonneville Salt Flats"

### FigureS1

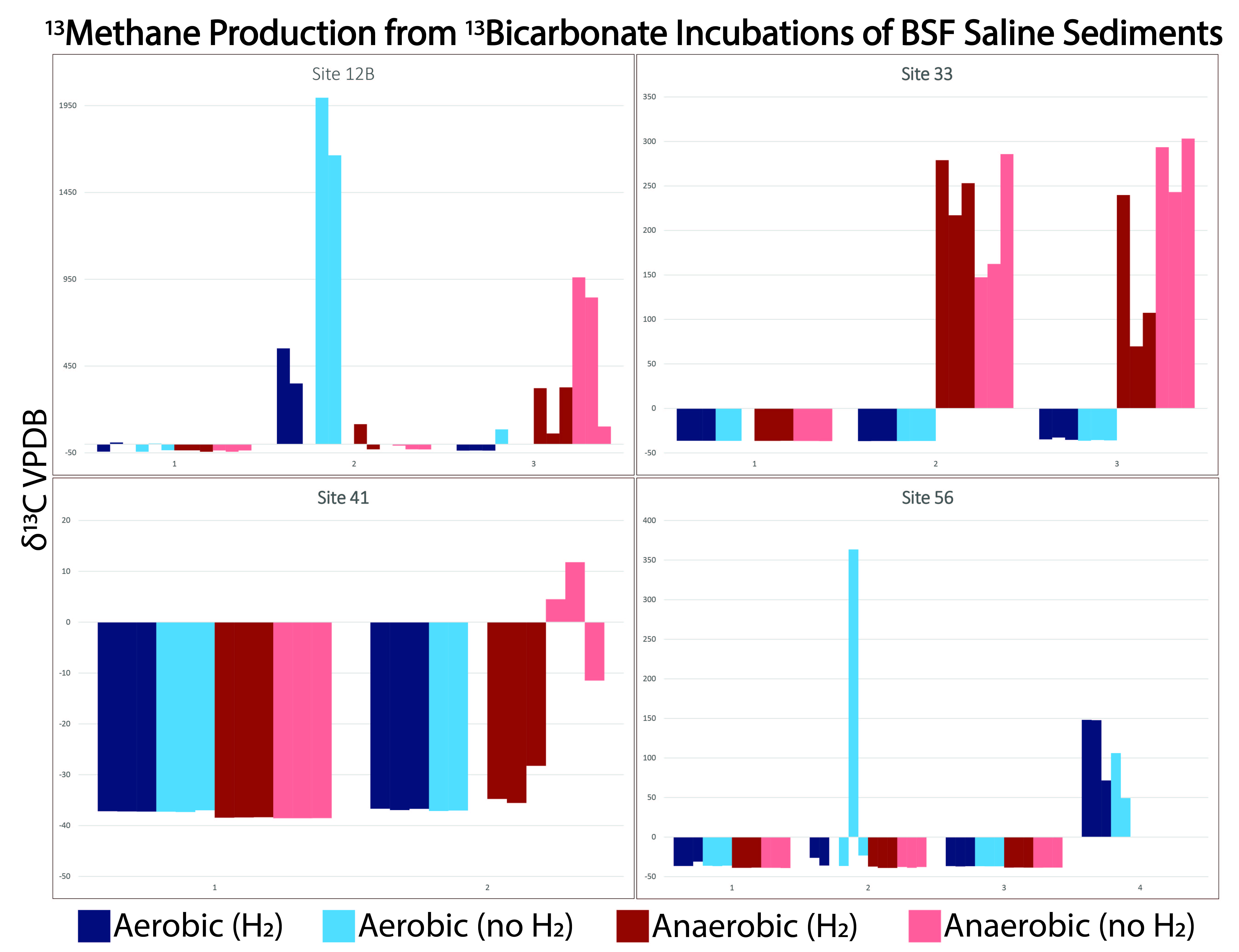

### FigureS2

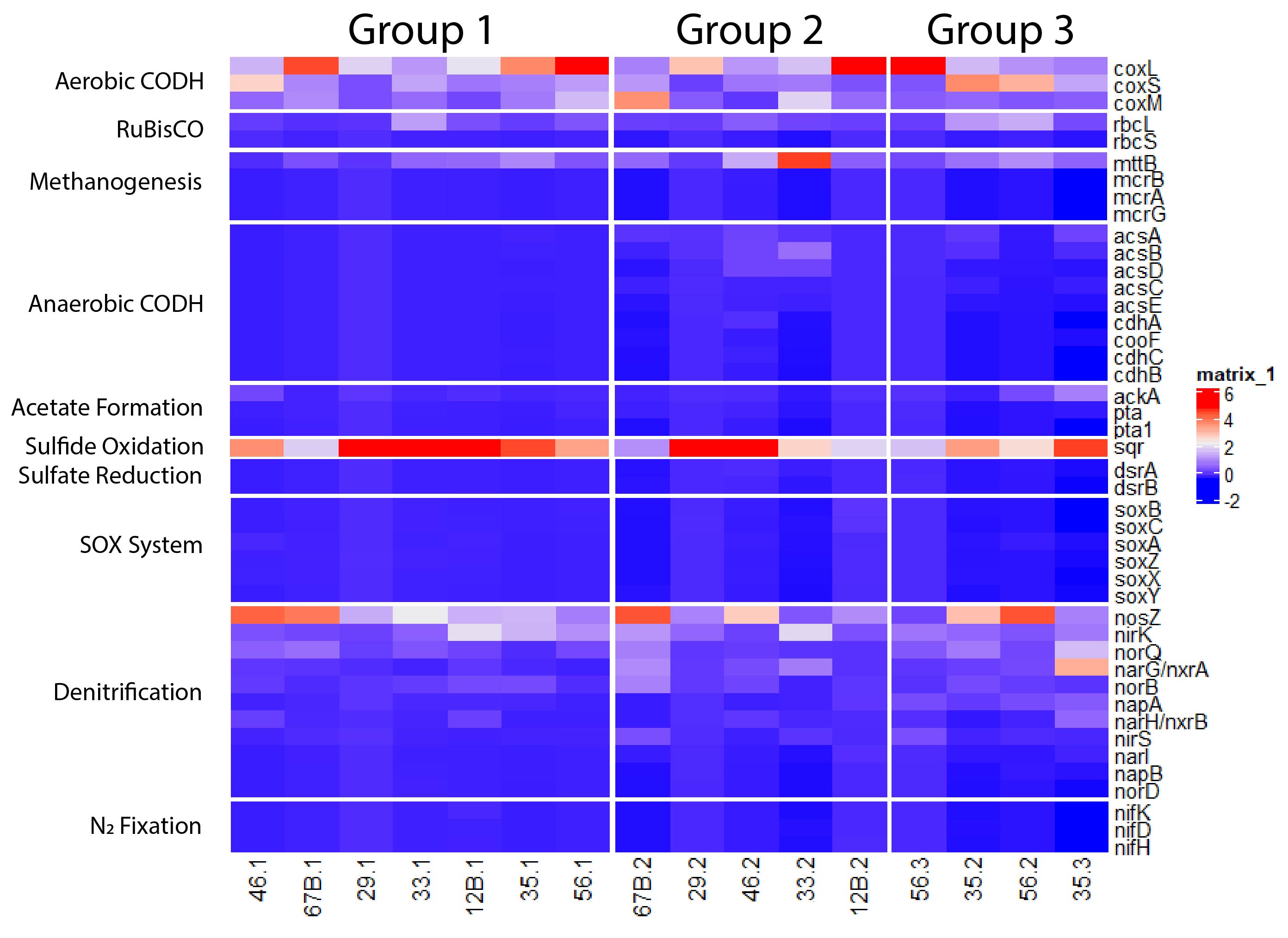
