## Supplementary material for "Metabolic Potential of Microbial Communities in the Hypersaline Sediments of the Bonneville Salt Flats": Tables S1 to S7

*Table S1*

| Sample | Mineralogical Group | Paired Reads | % Reads Mapped | # Contigs | N50 | Max Length (bp) | # Predicted Proteins | # Known Functions | % Unknown Functions |
|---|---|---|---|---|---|---|---|---|---|
| D12B.1 | 1 | 27,028,379 | 73.27 | 480,395 | 1,609 | 152,799 | 772,263 | 221,411 | 71.33 |
| D12B.2 | 2 | 36,398,584 | 76.72 | 529,453 | 1,867 | 232,657 | 873,020 | 265,247 | 69.62 |
| D67B.1 | 1 | 36,944,236 | 79.68 | 551,915 | 1,851 | 110,939 | 926,130 | 274,266 | 70.39 |
| D67B.2 | 2 | 41,065,986 | 71.34 | 718,467 | 1,950 | 248,332 | 1,222,333 | 387,170 | 68.33 |
| D29.1 | 1 | 27,262,857 | 75.68 | 431,625 | 2,162 | 231,711 | 723,205 | 235,225 | 67.47 |
| D29.2 | 2 | 31,153,558 | 68.34 | 514,747 | 1,772 | 231,545 | 831,006 | 271,565 | 67.32 |
| D33.1 | 1 | 31,759,001 | 78.86 | 503,866 | 2,222 | 222,646 | 864,854 | 272,038 | 68.55 |
| D33.2 | 2 | 30,502,979 | 71.02 | 556,953 | 1,880 | 183,134 | 930,075 | 289,647 | 68.86 |
| D35.1 | 1 | 53,706,656 | 75.09 | 795,975 | 2,322 | 223,962 | 1,369,835 | 434,454 | 68.28 |
| D35.2 | 3 | 44,663,651 | 74.56 | 768,295 | 1,461 | 88,995 | 1,212,160 | 325,402 | 73.16 |
| D35.3 | 3 | 54,664,207 | 70.81 | 1,046,983 | 1,281 | 232,405 | 1,610,078 | 369,933 | 77.02 |
| D56.1 | 1 | 35,366,014 | 77.89 | 554,687 | 2,239 | 231,467 | 961,743 | 303,059 | 68.49 |
| D56.2 | 3 | 25,485,574 | 67.47 | 497,380 | 1,467 | 205,540 | 775,528 | 196,824 | 74.62 |
| D56.3 | 3 | 43,499,921 | 72.23 | 733,529 | 1,982 | 206,649 | 1,236,617 | 394,497 | 68.10 |
| D46.1 | 1 | 45,073,753 | 77.47 | 802,472 | 1,481 | 521,643 | 1,298,167 | 357,923 | 72.43 |
| D46.2 | 2 | 32,815,788 | 65.5 | 660,420 | 1,479 | 162,191 | 1,060,402 | 302,249 | 71.50 |

*Table S2*

| BinSanity Bin Id | GTDB-Tk Classify Results | Bin ID | Completeness | Contamination | Strain Heterogeneity |
|---|---|---|---|---|---|
| BinSanityLC-kmean-bin_57-bin_0-refined_5 | d__Archaea; p__Halobacterota; c__Halobacteria; o__Halobacteriales; f__; g__; s__ | Halobacteriales.1 | 92 | 5.46 | 35.29 |
| BinSanityLC-kmean-bin_45-bin_0-refined_15 | d__Archaea; p__Halobacterota; c__Halobacteria; o__Halobacteriales; f__; g__; s__ | Halobacteriales.2 | 62 | 6.07 | 40.74 |
| BinSanityLC-kmean-bin_57-bin_0-refined_8 | d__Archaea; p__Halobacterota; c__Halobacteria; o__Halobacteriales; f__; g__; s__ | Halobacteriales.3 | 47 | 0 | 0 |
| BinSanityLC-kmean-bin_35-bin_0-refined_4 | d__Archaea; p__Halobacterota; c__Halobacteria; o__Halobacteriales; f__Haloarculaceae; g__; s__ | Haloarculaceae.1 | 90 | 13.91 | 15.79 |
| BinSanityLC-kmean-bin_36-bin_0-refined_3 | d__Archaea; p__Halobacterota; c__Halobacteria; o__Halobacteriales; f__Haloarculaceae; g__; s__ | Haloarculaceae.2 | 71 | 11.17 | 13.95 |
| BinSanityLC-kmean-bin_44-bin_0-refined_5 | d__Archaea; p__Halobacterota; c__Halobacteria; o__Halobacteriales; f__Haloarculaceae; g__; s__ | Haloarculaceae.3 | 67 | 6.81 | 25 |
| BinSanityLC-kmean-bin_61-bin_0-refined_12 | d__Archaea; p__Halobacterota; c__Halobacteria; o__Halobacteriales; f__Haloarculaceae; g__Halovenus; s__ | Halovenus | 67 | 8.27 | 46.43 |
| BinSanityLC-kmean-bin_53-bin_0-refined_1 | d__Archaea; p__Halobacterota; c__Halobacteria; o__Halobacteriales; f__Haloarculaceae; g__QS-1-69-70; s__ | Haloarculaceae.4 | 87 | 3.97 | 18.18 |
| BinSanityLC-kmean-bin_18-bin_0-refined_0 | d__Archaea; p__Halobacterota; c__Halobacteria; o__Halobacteriales; f__Haloarculaceae; g__SW-9-67-25; s__ | Haloarculaceae.5 | 90 | 5.48 | 52.38 |
| BinSanityLC-kmean-bin_53-bin_0-refined_2 | d__Archaea; p__Halobacterota; c__Halobacteria; o__Halobacteriales; f__Haloferacaceae; g__; s__ | Haloferacaceae.1 | 80 | 2.31 | 60 |
| BinSanityLC-kmean-bin_45-bin_0-refined_6 | d__Archaea; p__Halobacterota; c__Halobacteria; o__Halobacteriales; f__Haloferacaceae; g__; s__ | Haloferacaceae.2 | 49 | 0.38 | 0 |
| BinSanityLC-kmean-bin_45-bin_0-refined_10 | d__Archaea; p__Halobacterota; c__Halobacteria; o__Halobacteriales; f__Haloferacaceae; g__; s__ | Haloferacaceae.3 | 48 | 1.2 | 0 |
| BinSanityLC-kmean-bin_14-bin_0-refined_8 | d__Archaea; p__Halobacterota; c__Methanosarcinia; o__Methanosarcinales; f__Methanosarcinaceae; g__Methanohalobium; s__ | Methanohalobium | 99 | 0.65 | 0 |
| BinSanityLC-kmean-bin_57-bin_0-refined_3 | d__Archaea; p__Nanohaloarchaeota; c__Nanosalinia; o__Nanosalinales; f__; g__; s__ | Nanosalinales | 56 | 0.93 | 0 |
| BinSanityLC-kmean-bin_14-bin_0-refined_6 | d__Archaea; p__Thermoplasmatota; c__; o__; f__; g__; s__ | Thermoplasmatota.1 | 52 | 1.6 | 0 |
| BinSanityLC-kmean-bin_14-bin_0-refined_12 | d__Archaea; p__Thermoplasmatota; c__SW-10-69-26; o__; f__; g__; s__ | Thermoplasmatota.2 | 57 | 0.8 | 0 |
| BinSanityLC-kmean-bin_44-bin_0-refined_1 | d__Bacteria; p__Bacteroidota; c__Bacteroidia; o__Bacteroidales; f__UBA12077; g__; s__ | Bacteroidales | 76 | 8.46 | 48 |
| BinSanityLC-kmean-bin_57-bin_0-refined_4 | d__Bacteria; p__Bacteroidota; c__Bacteroidia; o__Flavobacteriales; f__PHOS-HE28; g__; s__ | Flavobacteriales | 58 | 5.25 | 88.24 |
| BinSanityLC-kmean-bin_85-bin_0 | d__Bacteria; p__Bacteroidota; c__Rhodothermia; o__Rhodothermales; f__Salinibacteraceae; g__Salinivenus; s__ | Salinivenus | 96 | 4.55 | 15.79 |
| BinSanityLC-kmean-bin_28-bin_0-refined_2 | d__Bacteria; p__Bipolaricaulota; c__Bipolaricaulia; o__RBG-16-55-9; f__RBG-16-55-9; g__; s__ | Bipolaricaulia | 83 | 11.76 | 100 |
| BinSanityLC-kmean-bin_24-bin_0-refined_1 | d__Bacteria; p__Cyanobacteria; c__Cyanobacteriia; o__Cyanobacteriales; f__Rubidibacteraceae; g__Halothece; s__ | Halothece | 87 | 2.9 | 83.33 |
| BinSanityLC-kmean-bin_44-bin_0-refined_9 | d__Bacteria; p__Firmicutes_F; c__Halanaerobiia; o__Halanaerobiales; f__; g__; s__ | Halanerobiales | 64 | 3.02 | 0 |
| BinSanityLC-kmean-bin_13-bin_0-refined_3 | d__Bacteria; p__Myxococcota; c__Bradimonadia; o__Bradymonadales; f__Bradymonadaceae; g__; s__ | Bradymonadaceae | 53 | 1.94 | 0 |
| BinSanityLC-kmean-bin_24-bin_0-refined_5 | d__Bacteria; p__Patescibacteria; c__Paceibacteria; o__SW-4-49-11; f__SW-4-49-11; g__SW-4-49-11; s__ | Paceibacteria | 80 | 2.59 | 100 |
| BinSanityLC-kmean-bin_73-bin_0-refined_4 | d__Bacteria; p__Proteobacteria; c__Alphaproteobacteria; o__UBA6615; f__; g__; s__ | Alphaproteobacteria | 47 | 1.3 | 100 |
| BinSanityLC-kmean-bin_55-bin_0-refined_6 | d__Bacteria; p__Proteobacteria; c__Gammaproteobacteria; o__Thiohalorhabdales; f__Thiohalorhabdaceae; g__Thiohalorhabdus; s__ | Thiohalorhabdaceae.1 | 80 | 0.89 | 0 |
| BinSanityLC-kmean-bin_92-bin_0-refined_3 | d__Bacteria; p__Proteobacteria; c__Gammaproteobacteria; o__Thiohalorhabdales; f__Thiohalorhabdaceae; g__Thiohalorhabdus; s__ | Thiohalorhabdaceae.2 | 59 | 3.41 | 66.67 |
| BinSanityLC-kmean-bin_92-bin_0-refined_6 | d__Bacteria; p__Verrucomicrobiota; c__Verrucomicrobiae; o__Pedosphaerales; f__AV2; g__; s__ | Pedosphaerales | 74 | 3.38 | 0 |

**Completeness, Contamination, and Strain Heterogeneity Determined by CheckM*

*Table S3*

| KEGG_ID | Name | Definition | Metabolism | 12B.1 | 12B.2 | 67B.1 | 67B.2 | 29.1 | 29.2 | 33.1 | 33.2 | 35.1 | 35.2 | 35.3 | 56.1 | 56.2 | 56.3 | 46.1 | 46.2 |
|---|---|---|---|---|---|---|---|---|---|---|---|---|---|---|---|---|---|---|---|
| K00192 | cdhA | anaerobic carbon-monoxide dehydrogenase, CODH/ACS complex subunit alpha | Carbon Metabolism | 0 | 0.935414579 | 0 | 0 | 0 | 0 | 0 | 2.185944648 | 0 | 0 | 0 | 0 | 0 | 0 | 0 | 50.61576593 |
| K00193 | cdhC | acetyl-CoA decarbonylase/synthase, CODH/ACS complex subunit beta | Carbon Metabolism | 0 | 0 | 0 | 0.963611375 | 0 | 0 | 0 | 0.929096702 | 0.307160551 | 0 | 0 | 0 | 0 | 0.51350565 | 0 | 12.97963333 |
| K00194 | cdhD_acsD | acetyl-CoA decarbonylase/synthase, CODH/ACS complex subunit delta | Carbon Metabolism | 0 | 0.538442983 | 0 | 7.784968174 | 0 | 4.158399345 | 0 | 98.28689317 | 2.652937789 | 7.58725181 | 10.58678273 | 0 | 2.296986436 | 2.001699215 | 0 | 124.9395321 |
| K00195 | cdhB | anaerobic carbon-monoxide dehydrogenase, CODH/ACS complex subunit epsilon | Carbon Metabolism | 0 | 0 | 0 | 0 | 0 | 0 | 0 | 0 | 0 | 0 | 0 | 0 | 0 | 0 | 0 | 0.618446008 |
| K00196 | cooF | anaerobic carbon-monoxide dehydrogenase iron sulfur subunit | Carbon Metabolism | 0 | 1.162128863 | 0 | 3.961984015 | 0 | 0 | 0 | 1.775037695 | 0 | 0 | 6.499610805 | 0 | 0 | 0.183751872 | 0 | 1.936733407 |
| K00197 | cdhE_acsC | acetyl-CoA decarbonylase/synthase, CODH/ACS complex subunit gamma | Carbon Metabolism | 0 | 1.291922914 | 0 | 23.47220044 | 0 | 9.967936592 | 0 | 30.20872608 | 4.95282723 | 12.81289452 | 18.46734418 | 0 | 0.793250796 | 0.586962324 | 0 | 20.47170479 |
| K00198 | cooS_acsA | anaerobic carbon-monoxide dehydrogenase catalytic subunit | Carbon Metabolism | 0.375775832 | 4.581165689 | 0.242601133 | 63.83678833 | 0 | 43.67182779 | 0 | 61.12756575 | 15.67962822 | 40.14547597 | 64.44255786 | 0 | 4.894988429 | 3.784149588 | 0.934403097 | 119.5487705 |
| K00297 | metF_MTHFR | methylenetetrahydrofolate reductase (NADPH) | Carbon Metabolism | 1122.268425 | 539.0172148 | 342.554246 | 309.8658433 | 356.6670654 | 769.7316959 | 220.5766966 | 743.8391725 | 267.7077216 | 405.1184348 | 924.4721429 | 815.3326892 | 396.1817699 | 1026.842155 | 2254.288659 | 756.7009766 |
| K00625 | pta | phosphate acetyltransferase | Carbon Metabolism | 0.622963467 | 5.165148077 | 6.086834169 | 23.67923094 | 1.053407525 | 2.185432799 | 0.22458495 | 8.903856221 | 0.423717073 | 1.00605456 | 13.87314845 | 7.580180643 | 0.929817162 | 3.096566118 | 3.774871478 | 20.15340686 |
| K00656 | pflD | pyruvate formate lyase | Carbon Metabolism | 0 | 11.07832831 | 0 | 3.885172197 | 0 | 2.887412681 | 0 | 7.349479378 | 0.20794603 | 0.793749288 | 6.745458584 | 0 | 0 | 0.341934538 | 0.907535836 | 55.60637695 |
| K00925 | ackA | acetate kinase | Carbon Metabolism | 27.49234652 | 32.90509058 | 7.948359056 | 31.29523186 | 39.96239271 | 20.78118927 | 6.280419655 | 14.01641238 | 18.10700632 | 12.80527491 | 151.4739714 | 15.62263996 | 73.5485566 | 33.74773607 | 94.45534767 | 40.55197187 |
| K01491 | folD | methylenetetrahydrofolate dehydrogenase (NADP+) / methenyltetrahydrofolate cyclohydrolase | Carbon Metabolism | 311.646744 | 1845.112593 | 321.1860743 | 361.0539997 | 492.6048427 | 508.2944989 | 839.9126764 | 1027.167336 | 512.8890023 | 619.5681565 | 324.9805403 | 1827.099826 | 1088.224955 | 628.0690658 | 2322.10778 | 308.1263068 |
| K01601 | rbcL | ribulose-bisphosphate carboxylase large chain | Carbon Metabolism | 110.6067392 | 144.3320862 | 44.73015366 | 91.37150017 | 33.37458904 | 104.9098722 | 166.1190644 | 97.58963196 | 91.72723877 | 176.5609547 | 73.47539198 | 108.0355469 | 241.9828171 | 95.08241625 | 66.80455684 | 206.0447548 |
| K01602 | rbcS | ribulose-bisphosphate carboxylase small chain | Carbon Metabolism | 5.450925647 | 12.14696142 | 6.807411081 | 9.233737956 | 5.80147289 | 3.668554337 | 1.900053199 | 2.010844352 | 4.965524421 | 9.209863215 | 10.80267814 | 5.519086227 | 12.76499755 | 4.217196015 | 26.41252967 | 3.542787076 |
| K01938 | fhs | formate--tetrahydrofolate ligase | Carbon Metabolism | 528.6083888 | 852.3408805 | 848.3042456 | 1033.747333 | 571.9332281 | 764.9493035 | 279.0044087 | 365.5287948 | 443.3054431 | 261.8666532 | 416.4782697 | 783.8852756 | 155.3160189 | 205.2263504 | 332.3724006 | 849.1088933 |
| K03518 | coxS | aerobic carbon-monoxide dehydrogenase small subunit | Carbon Metabolism | 211.8322553 | 279.9939847 | 369.7167636 | 315.3490208 | 142.6428744 | 125.154831 | 174.4750253 | 224.194873 | 336.1913497 | 475.2162286 | 212.0985496 | 284.2600316 | 485.680917 | 202.8549563 | 632.6754702 | 301.1200753 |
| K03519 | coxM | aerobic carbon-monoxide dehydrogenase medium subunit | Carbon Metabolism | 92.16690733 | 455.0417024 | 390.2510302 | 829.6159472 | 142.7894031 | 279.2013368 | 94.12554247 | 440.0772534 | 313.8457701 | 104.0937756 | 103.37306 | 359.2063958 | 89.86530316 | 248.1318141 | 172.1223263 | 78.18218935 |
| K03520 | coxL | aerobic carbon-monoxide dehydrogenase large subunit | Carbon Metabolism | 512.6548717 | 3737.721826 | 1505.566521 | 257.4317567 | 775.5454277 | 1520.507043 | 156.4082256 | 397.4896783 | 1206.672374 | 224.5002992 | 151.6277956 | 1007.569238 | 195.5860745 | 2740.596039 | 382.1272539 | 425.3953543 |
| K05299 | fdhA | formate dehydrogenase (NADP+) alpha subunit | Carbon Metabolism | 22.84139375 | 6.078876176 | 32.76014464 | 238.8448648 | 2.276605185 | 87.67367831 | 1.009404547 | 49.70956727 | 47.25016002 | 17.72461977 | 11.88562163 | 1.716084961 | 54.46023044 | 53.65816772 | 2.127172191 | 171.4876071 |
| K13788 | pta | phosphate acetyltransferase | Carbon Metabolism | 0 | 0 | 0 | 0 | 0 | 0 | 0 | 0 | 0 | 0 | 0.17304618 | 0 | 0.738828472 | 0 | 0.622936976 | 2.291385407 |
| K14138 | acsB | acetyl-CoA synthase | Carbon Metabolism | 0.956695369 | 3.752834643 | 0 | 26.89996292 | 0 | 39.67909503 | 0 | 195.4347724 | 9.473839237 | 30.64794642 | 31.34653964 | 0 | 2.983845866 | 3.634935885 | 0.500545358 | 124.8516045 |
| K15022 | fdhB | formate dehydrogenase (NADP+) beta subunit | Carbon Metabolism | 0 | 2.679708989 | 0 | 1.460691663 | 0 | 2.505220454 | 0 | 20.78384797 | 1.016764476 | 1.626525588 | 0.495397086 | 0 | 0.54589232 | 1.766096597 | 0 | 18.48374157 |
| K15023 | acsE | 5-methyltetrahydrofolate corrinoid/iron sulfur protein methyltransferase | Carbon Metabolism | 0 | 0.230358978 | 0 | 7.334569591 | 0 | 4.165177539 | 0 | 25.11579482 | 0.624893729 | 5.292392568 | 8.641828363 | 0 | 0.273576953 | 0.423176653 | 0 | 16.51008656 |
| K00244 | frdA | fumarate reductase flavoprotein subunit | Electron Transport | 24.42933874 | 31.04211238 | 44.3465055 | 136.992168 | 5.757648128 | 11.10833005 | 15.85459609 | 27.24682823 | 36.5405962 | 38.37000269 | 114.858414 | 29.32093162 | 101.0703173 | 102.1767846 | 11.6352047 | 45.85184826 |
| K00245 | frdB | fumarate reductase iron-sulfur subunit | Electron Transport | 0 | 0.25607334 | 0 | 0.595625625 | 0 | 0 | 0 | 0 | 0 | 0 | 1.470152384 | 0 | 0 | 0.734247789 | 0 | 0 |
| K00246 | frdC | fumarate reductase subunit C | Electron Transport | 0 | 0.234280052 | 0 | 0.237613233 | 0 | 0 | 0 | 0 | 0.129057349 | 0 | 0 | 0 | 0 | 0 | 0 | 0 |
| K00247 | frdD | fumarate reductase subunit D | Electron Transport | 0 | 0 | 0 | 0 | 0 | 0 | 0 | 0 | 0 | 0 | 0.155196082 | 0 | 0 | 0 | 0 | 0 |
| K00334 | nuoE | NADH-quinone oxidoreductase subunit E | Electron Transport | 165.0048948 | 128.9307763 | 181.4075166 | 83.43294781 | 273.2316495 | 159.1090635 | 580.1402507 | 206.1361157 | 138.8485814 | 343.5655628 | 220.1634833 | 106.306534 | 164.1672575 | 119.8204012 | 461.4716018 | 212.4562759 |
| K00335 | nuoF | NADH-quinone oxidoreductase subunit F | Electron Transport | 897.3787746 | 228.7188654 | 613.8422338 | 661.4326296 | 128.974846 | 600.3875596 | 216.5414276 | 754.5351285 | 111.9770437 | 729.6715073 | 299.678097 | 116.1667796 | 253.070557 | 245.9575932 | 200.5855263 | 272.9848402 |
| K00336 | nuoG | NADH-quinone oxidoreductase subunit G | Electron Transport | 696.7962991 | 615.028355 | 506.4285186 | 732.1898279 | 790.7477806 | 133.0785396 | 678.3903302 | 232.1825926 | 354.4424544 | 600.5295717 | 282.3707167 | 503.0538109 | 1659.536156 | 108.8246136 | 75.59950147 | 118.7925931 |
| K02274 | coxA | cytochrome c oxidase subunit I | Electron Transport | 860.1585892 | 1195.299983 | 2365.483285 | 7721.808832 | 1729.151408 | 5773.311566 | 469.3350872 | 614.2001526 | 1732.823297 | 672.2687128 | 1181.304264 | 1385.957933 | 1854.436354 | 1408.744373 | 3663.127343 | 2272.656827 |
| K02275 | coxB | cytochrome c oxidase subunit II | Electron Transport | 1403.827001 | 2142.843355 | 542.1315715 | 1399.869365 | 545.3241792 | 337.4267753 | 877.2819339 | 1045.314598 | 533.7987659 | 1341.16601 | 1782.534512 | 451.0049707 | 430.4222745 | 747.078544 | 2418.806248 | 1047.30571 |
| K02276 | coxC | cytochrome c oxidase subunit III | Electron Transport | 623.5569422 | 185.6767906 | 206.0077182 | 672.0134217 | 189.2988814 | 293.9386366 | 517.7624266 | 209.6662891 | 482.0244934 | 631.1645752 | 503.5833456 | 403.9120299 | 313.2736682 | 196.8817688 | 467.6049123 | 454.7583785 |
| K02277 | coxD | cytochrome c oxidase subunit IV | Electron Transport | 286.111366 | 174.7239895 | 472.68333 | 191.8402135 | 199.6425398 | 369.4823604 | 168.9556747 | 172.9477139 | 138.476155 | 194.3094249 | 478.3746051 | 186.6790076 | 491.3586454 | 155.7996737 | 239.2170969 | 204.1127455 |
| K05576 | ndhE | NAD(P)H-quinone oxidoreductase subunit 4L | Electron Transport | 0.898619166 | 0.87948339 | 1.323842535 | 1.732755087 | 0.675205672 | 1.443702575 | 0.375265337 | 0.245880004 | 0.707547274 | 2.408725427 | 0.346092359 | 0.361992531 | 2.649235406 | 0.154398681 | 5.596485978 | 0.255380934 |
| K05577 | ndhF | NAD(P)H-quinone oxidoreductase subunit 5 | Electron Transport | 1069.193166 | 411.1396559 | 565.7347126 | 184.2423568 | 968.5124579 | 956.8550346 | 265.2647611 | 327.0501353 | 1346.765945 | 184.4425531 | 135.3657693 | 772.1356925 | 828.4958308 | 517.7294083 | 652.1287875 | 302.1288447 |
| K05578 | ndhG | NAD(P)H-quinone oxidoreductase subunit 6 | Electron Transport | 20.8682992 | 6.638573451 | 7.544976868 | 7.451175245 | 6.748654151 | 4.015085733 | 3.415578325 | 2.373227776 | 3.875891357 | 23.27352178 | 4.300310415 | 4.942161303 | 13.39602736 | 1.093011155 | 14.9142417 | 1.028089531 |
| K15408 | coxAC | cytochrome c oxidase subunit I+III | Electron Transport | 472.4476208 | 916.6020116 | 329.2842773 | 1447.014535 | 370.7500236 | 1980.640968 | 749.6873263 | 515.2236943 | 1496.087125 | 1374.154742 | 1497.152851 | 183.4708162 | 1494.81925 | 1369.488546 | 424.9898855 | 619.5291159 |
| K08738 | CYC | cytochrome c | Energy | 109.7113696 | 84.67912557 | 91.77928874 | 10.86616781 | 17.02917134 | 52.15797589 | 16.00382646 | 48.22199974 | 23.11442409 | 57.69975051 | 173.7519291 | 9.784218877 | 84.20558051 | 21.74704182 | 13.54431889 | 39.61498333 |
| K00436 | hoxH | NAD-reducing hydrogenase large subunit | Energy (from H2) | 48.4646211 | 19.2101252 | 13.54282663 | 62.60417042 | 15.97331933 | 41.46462193 | 0.160024757 | 32.48136662 | 4.007846176 | 32.7644755 | 11.50187377 | 0.809310603 | 10.05154565 | 0.82978496 | 16.44603042 | 38.87111579 |
| K06281 | hyaB_hybC | hydrogenase large subunit | Energy (from H2) | 68.11662401 | 358.2673678 | 137.1230101 | 138.7395272 | 38.34233474 | 39.17201082 | 88.09348772 | 317.1638027 | 21.74552665 | 60.17090454 | 96.5452189 | 9.76195472 | 25.71526459 | 129.942441 | 137.9945135 | 138.7257813 |
| K06282 | hyaA_hybO | hydrogenase small subunit | Energy (from H2) | 32.20152175 | 57.82290575 | 95.3308959 | 264.0612689 | 38.68343666 | 23.46942096 | 24.20891263 | 104.9462764 | 26.46290219 | 92.07984407 | 146.0354221 | 7.170474833 | 30.05997654 | 154.4015806 | 126.830989 | 275.9751781 |
| K06441 | E1.12.7.2G | ferredoxin hydrogenase gamma subunit | Energy (from H2) | 0 | 0 | 0 | 0 | 0 | 0 | 0 | 0 | 0 | 0 | 0 | 0 | 0 | 0 | 0 | 0 |
| K17992 | hndB | NADP-reducing hydrogenase subunit HndB | Energy (from H2) | 0 | 2.512922945 | 0 | 0.300638466 | 0 | 0.463466537 | 0 | 4.823131195 | 0 | 0.490806679 | 1.688208911 | 0 | 0 | 0 | 0 | 39.37933735 |
| K17993 | hydA | sulfhydrogenase subunit alpha | Energy (from H2) | 0 | 0 | 0 | 0 | 0 | 0 | 0 | 0 | 0 | 0 | 0.305145895 | 0 | 0 | 0 | 0 | 1.085082606 |
| K17994 | hydD | sulfhydrogenase subunit delta | Energy (from H2) | 0 | 0 | 0 | 0 | 0 | 0 | 0 | 0.719255713 | 0 | 0 | 0 | 0 | 0 | 0 | 0 | 1.384995675 |
| K18005 | hoxF | NiFe hydrogenase diaphorase moiety large subunit | Energy (from H2) | 0 | 8.808928189 | 1.737039941 | 1.717746732 | 0 | 2.389494091 | 0 | 4.587609291 | 0 | 0 | 15.58660834 | 0 | 0 | 5.18628286 | 0 | 7.703961008 |
| K18006 | hoxU | NiFe hydrogenase diaphorase moiety small subunit | Energy (from H2) | 0 | 0.31762958 | 0 | 0.475225973 | 0 | 0 | 0 | 1.869037124 | 0 | 0 | 0 | 0 | 0 | 0.270359707 | 0 | 1.941257682 |
| K18007 | hoxY | NAD-reducing hydrogenase small subunit | Energy (from H2) | 4.712175424 | 4.530530868 | 1.977639702 | 6.336691438 | 3.386848939 | 6.067891872 | 0 | 4.543280102 | 0.767633778 | 6.247975046 | 5.990420462 | 0.148303079 | 2.153793487 | 0.263600727 | 6.35089973 | 7.092128755 |
| K18016 | mbhL | membrane-bound hydrogenase subunit alpha | Energy (from H2) | 0 | 0 | 0 | 0.488165503 | 0 | 3.47599338 | 0 | 39.703946 | 0 | 0.192579878 | 5.376023085 | 0 | 0 | 0 | 0 | 31.76536625 |
| K18017 | mbhK | membrane-bound hydrogenase subunit beta | Energy (from H2) | 0 | 0 | 0 | 0 | 0 | 0.586856778 | 0 | 1.688788177 | 0 | 0 | 0.415575907 | 0 | 0 | 0 | 0 | 4.375957092 |
| K18023 | mbhJ | membrane-bound hydrogenase subunit mbhJ | Energy (from H2) | 0 | 0 | 0 | 0.220110552 | 0 | 0.413809489 | 0 | 3.779301933 | 0 | 0 | 1.209003439 | 0 | 0 | 0 | 0 | 3.951802317 |
| K18330 | hndA | NADP-reducing hydrogenase subunit HndA | Energy (from H2) | 0.450749675 | 0 | 0 | 0 | 0 | 0 | 0 | 1.256493929 | 0 | 0.226909348 | 1.796893236 | 0 | 1.155695511 | 0 | 0 | 0.553178145 |
| K18331 | hndC | NADP-reducing hydrogenase subunit HndC | Energy (from H2) | 7.417401019 | 1.97714191 | 0 | 4.327277727 | 0 | 7.995557489 | 0 | 41.63903823 | 1.004041447 | 6.483139145 | 7.750515068 | 0 | 0.547153906 | 1.302217697 | 0 | 6.674272793 |
| K18332 | hndD | NADP-reducing hydrogenase subunit HndD | Energy (from H2) | 0 | 3.588542627 | 0 | 2.608377265 | 0 | 0 | 0 | 145.9869475 | 0 | 0.389471098 | 1.853039041 | 0 | 0 | 0 | 0 | 66.83025587 |
| K16161 | mmoC | methane monooxygenase component C | Methane Metabolism | 0 | 0 | 0 | 0 | 0 | 0 | 0 | 0 | 0 | 0 | 0 | 0 | 0 | 0.239835076 | 0 | 0 |
| K00123 | fdoG, fdhF, fdwA | formate dehydrogenase major subunit | Methanogenesis | 982.8720377 | 1215.015465 | 891.7294672 | 1376.203101 | 1516.022592 | 387.6057445 | 308.7077709 | 1084.456656 | 2310.264966 | 1255.017457 | 902.9963455 | 1559.963933 | 893.1257458 | 1077.551424 | 1990.591818 | 785.9073298 |
| K00124 | fdoH, fdsB | formate dehydrogenase iron-sulfur subunit | Methanogenesis | 26.63991717 | 35.94472137 | 29.49165086 | 27.79395604 | 27.30104475 | 36.01939754 | 11.13043599 | 65.52031132 | 25.53202092 | 78.1829297 | 8.533393189 | 10.87170681 | 20.83736976 | 26.55714918 | 32.91199233 | 32.5033986 |
| K00125 | fdhB | formate dehydrogenase (coenzyme F420) beta subunit | Methanogenesis | 0 | 0 | 0 | 2.728325779 | 0 | 0 | 0 | 0.195330876 | 0.334535547 | 0 | 0 | 0 | 0 | 0 | 0 | 1.645701026 |
| K00127 | fdoI, fdsG | formate dehydrogenase subunit gamma | Methanogenesis | 7.294786952 | 12.89170124 | 4.617633163 | 16.64034174 | 3.22208164 | 6.189236605 | 2.669496958 | 19.61489649 | 16.45301265 | 5.722952669 | 4.907823621 | 1.997004236 | 5.486085796 | 16.36411738 | 12.33309069 | 11.68573864 |
| K00200 | fwdA, fmdA | formylmethanofuran dehydrogenase subunit A | Methanogenesis | 0 | 0 | 0 | 1.9759506 | 0 | 0 | 0 | 1.131771909 | 0.327365326 | 0 | 1.341048448 | 0 | 0 | 1.452997187 | 0 | 34.60838547 |
| K00201 | fwdB, fmdB | formylmethanofuran dehydrogenase subunit B | Methanogenesis | 0 | 0 | 0 | 0.856122163 | 0 | 0 | 0 | 1.112678034 | 0.106677443 | 0 | 0 | 0 | 0 | 2.555557709 | 0 | 7.624682282 |
| K00202 | fwdC, fmdC | formylmethanofuran dehydrogenase subunit C | Methanogenesis | 0 | 0 | 0 | 1.110746509 | 0 | 0 | 0 | 2.67688272 | 0 | 0 | 0 | 0 | 0 | 0.804413412 | 0 | 5.409082477 |
| K00203 | fwdD, fmdD | formylmethanofuran dehydrogenase subunit D | Methanogenesis | 0 | 0 | 0 | 0 | 0 | 0 | 0 | 1.007334642 | 0 | 0 | 0 | 0 | 0 | 0 | 0 | 13.74868014 |
| K00205 | fwdF, fmdF | 4Fe-4S ferredoxin | Methanogenesis | 0 | 0.103488389 | 0 | 0.235907783 | 0 | 0 | 0 | 1.324365303 | 0 | 0 | 0 | 0 | 0 | 0 | 0 | 12.72696136 |
| K00319 | mtd | methylenetetrahydromethanopterin dehydrogenase | Methanogenesis | 0 | 0 | 0 | 0 | 0 | 0 | 0 | 0.607582328 | 0 | 0 | 0 | 0 | 0 | 0 | 0 | 18.56287641 |
| K00320 | mer | 5,10-methylenetetrahydromethanopterin reductase | Methanogenesis | 104.9898186 | 130.1387006 | 54.16705929 | 349.9348001 | 422.0470455 | 85.77578403 | 586.5000132 | 74.23299791 | 161.5743385 | 117.2605644 | 61.30757892 | 176.8363241 | 177.6881404 | 181.9526662 | 44.51107418 | 240.1039124 |
| K00399 | mcrA | methyl-coenzyme M reductase alpha subunit | Methanogenesis | 0 | 0.636483601 | 0 | 0 | 0 | 0 | 0 | 0.253158919 | 0 | 0 | 0 | 0 | 0 | 0 | 0 | 2.783548189 |
| K00401 | mcrB | methyl-coenzyme M reductase beta subunit | Methanogenesis | 0 | 1.556350421 | 0 | 0 | 0 | 0 | 0 | 0.864727026 | 0 | 0 | 0 | 0 | 0 | 0 | 0 | 3.58528009 |
| K00402 | mcrG | methyl-coenzyme M reductase gamma subunit | Methanogenesis | 0 | 0.469225728 | 0 | 0 | 0 | 0 | 0 | 0.614863551 | 0 | 0 | 0 | 0 | 0 | 0 | 0 | 1.809430221 |
| K00441 | frhB | coenzyme F420 hydrogenase subunit beta | Methanogenesis | 409.946136 | 323.6862686 | 195.2285878 | 412.7075927 | 69.52237467 | 225.2845703 | 205.0822267 | 356.6852756 | 77.97551659 | 413.0616163 | 548.1446772 | 278.9500585 | 349.7056599 | 1764.409993 | 420.3720307 | 409.1918953 |
| K00442 | frhD | coenzyme F420 hydrogenase subunit delta | Methanogenesis | 0 | 0 | 0 | 1.70016467 | 0 | 3.208066439 | 0 | 9.990767914 | 1.123037011 | 1.905486781 | 2.417026519 | 0 | 0.771718547 | 0.36473027 | 0 | 5.999116677 |
| K00577 | mtrA | tetrahydromethanopterin S-methyltransferase subunit A | Methanogenesis | 0 | 0.88190034 | 0.615390915 | 23.49142928 | 0.459433703 | 3.765663261 | 0 | 3.671134094 | 0.672264586 | 0.342597977 | 4.5964381 | 0 | 0 | 2.067863382 | 0.459642734 | 20.47098539 |
| K00578 | mtrB | tetrahydromethanopterin S-methyltransferase subunit B | Methanogenesis | 0 | 0 | 0 | 0 | 0 | 0.459227153 | 0 | 0 | 0 | 0 | 0 | 0 | 0 | 0 | 0 | 0 |
| K00579 | mtrC | tetrahydromethanopterin S-methyltransferase subunit C | Methanogenesis | 0 | 0 | 0 | 0 | 0 | 0 | 0 | 0.268154652 | 0 | 0.30820327 | 0 | 0 | 0 | 0 | 0 | 1.472987228 |
| K00580 | mtrD | tetrahydromethanopterin S-methyltransferase subunit D | Methanogenesis | 0 | 0 | 0 | 0 | 0 | 0 | 0 | 0.640447496 | 0 | 0 | 0 | 0 | 0 | 0 | 0 | 1.441253356 |
| K00581 | mtrE | tetrahydromethanopterin S-methyltransferase subunit E | Methanogenesis | 0 | 0 | 0 | 0 | 0 | 0 | 0 | 1.557001193 | 0 | 0 | 0 | 0 | 0 | 0 | 0 | 2.792460851 |
| K00582 | mtrF | tetrahydromethanopterin S-methyltransferase subunit F | Methanogenesis | 0 | 0 | 0 | 0 | 0 | 0 | 0 | 0 | 0 | 0 | 0 | 0 | 0 | 0 | 0 | 0 |
| K00583 | mtrG | tetrahydromethanopterin S-methyltransferase subunit G | Methanogenesis | 0 | 0 | 0 | 0 | 0 | 0 | 0 | 0 | 0 | 0 | 0 | 0 | 0 | 0 | 0 | 0 |
| K00584 | mtrH | tetrahydromethanopterin S-methyltransferase subunit H | Methanogenesis | 160.2045888 | 41.89030745 | 16.56311709 | 75.01663274 | 2.864305042 | 25.29388481 | 3.226870328 | 56.16031082 | 14.10333102 | 39.77582056 | 124.6831173 | 15.3749583 | 36.18855896 | 24.902006 | 13.99990242 | 52.55249135 |
| K00672 | ftr | formylmethanofuran--tetrahydromethanopterin N-formyltransferase | Methanogenesis | 0 | 0.400891569 | 0 | 4.0897917 | 0 | 0 | 0 | 11.56399236 | 0.935923232 | 0 | 28.90333594 | 0 | 0 | 4.254349628 | 0 | 12.71944755 |
| K01499 | mch | methenyltetrahydromethanopterin cyclohydrolase | Methanogenesis | 149.8061133 | 337.4386571 | 110.1790689 | 822.358293 | 405.239217 | 98.46155042 | 167.2584068 | 62.3758634 | 213.4737374 | 34.048484 | 25.29892261 | 1848.128973 | 54.76692017 | 164.705329 | 150.072235 | 97.37902379 |
| K01895 | ACSS1_2, acs | acetyl-CoA synthetase | Methanogenesis | 5612.60783 | 4693.584129 | 1255.959549 | 3839.861163 | 1931.833009 | 1002.163495 | 5712.915535 | 2122.121391 | 4774.955826 | 2190.955957 | 3362.032016 | 2172.56816 | 2484.246045 | 2873.562285 | 1187.457188 | 3438.065298 |
| K03388 | hdrA2 | heterodisulfide reductase subunit A2 | Methanogenesis | 0.556415474 | 18.3269751 | 2.609335793 | 244.0124921 | 0 | 98.64305539 | 0 | 138.3278487 | 101.5952446 | 17.04136003 | 58.70665133 | 0 | 11.45220203 | 66.55029738 | 1.034552883 | 577.1607995 |
| K03389 | hdrB2 | heterodisulfide reductase subunit B2 | Methanogenesis | 0 | 2.191606278 | 0 | 38.67871172 | 0 | 6.450814637 | 0 | 51.95804224 | 2.468725171 | 6.600408059 | 41.31109297 | 0 | 1.091785825 | 1.769711439 | 0 | 81.16677202 |
| K03390 | hdrC2 | heterodisulfide reductase subunit C2 | Methanogenesis | 0 | 0.181104406 | 0 | 9.34866725 | 0 | 2.048678972 | 0 | 2.198866299 | 0 | 0 | 12.09886302 | 0 | 0 | 3.451912781 | 0 | 23.54909105 |
| K04480 | mtaB | methanol---5-hydroxybenzimidazolylcobamide Co-methyltransferase | Methanogenesis | 0 | 3.9325588 | 0 | 2.127168165 | 0 | 0 | 0 | 4.261651224 | 0 | 0 | 4.812041023 | 0 | 0 | 0.486921099 | 0 | 37.28706001 |
| K08264 | hdrD | heterodisulfide reductase subunit D | Methanogenesis | 0 | 0.315506078 | 0 | 0 | 0 | 2.597012262 | 0 | 0.441033871 | 0 | 0 | 0 | 0 | 0 | 0 | 0 | 2.578524993 |
| K08265 | hdrE | heterodisulfide reductase subunit E | Methanogenesis | 0 | 0 | 0 | 0.143745435 | 0 | 0 | 0 | 0 | 0 | 0 | 0 | 0 | 0 | 0 | 0 | 3.172803699 |
| K11260 | fwdG | 4Fe-4S ferredoxin | Methanogenesis | 0 | 0.52938245 | 0 | 0 | 0 | 0 | 0 | 0.574331588 | 0 | 0 | 0 | 0 | 0 | 0 | 0 | 1.042621566 |
| K11261 | fwdE, fmdE | formylmethanofuran dehydrogenase subunit E | Methanogenesis | 8.188290792 | 21.02636103 | 1.221251485 | 21.10483449 | 2.149547892 | 11.90808162 | 0.917640497 | 38.69638047 | 6.81178457 | 9.059976111 | 29.95210231 | 1.414329154 | 0 | 9.061267426 | 1.956329551 | 235.2023481 |
| K14080 | mtaA | methyl-Co(III) methanol-specific corrinoid protein:coenzyme M methyltransferase | Methanogenesis | 0 | 3.093023899 | 0 | 5.306321478 | 0 | 0 | 0 | 4.680023336 | 0 | 0 | 2.07655524 | 0 | 0 | 0 | 0 | 25.83880584 |
| K14081 | mtaC | methanol corrinoid protein | Methanogenesis | 0 | 0.193858633 | 0 | 0 | 0 | 0 | 0 | 1.406589792 | 0 | 0 | 0 | 0 | 0 | 0 | 0 | 3.786898089 |
| K14082 | mtbA | methyl-Co(III) methylamine-specific corrinoid protein:coenzyme M methyltransferase | Methanogenesis | 0 | 0 | 0 | 0 | 0 | 0 | 0 | 1.181988568 | 0 | 0 | 0 | 0 | 0 | 0 | 0 | 3.356612361 |
| K14083 | mttB | trimethylamine---corrinoid protein Co-methyltransferase | Methanogenesis | 182.2728708 | 377.5980102 | 157.5951223 | 189.6150917 | 36.19089889 | 86.52063222 | 86.24058803 | 970.8169334 | 361.9374881 | 119.2557538 | 112.8186195 | 108.2644202 | 185.9482697 | 145.2122473 | 31.19002578 | 498.9867634 |
| K14084 | mttC | trimethylamine corrinoid protein | Methanogenesis | 0 | 0.964197752 | 0.220875573 | 11.92686017 | 0 | 3.800209456 | 0 | 13.98208805 | 0.767190853 | 5.56111426 | 9.344219637 | 0 | 0 | 0 | 0 | 19.07117791 |
| K14126 | mvhA, vhuA, vhcA | F420-non-reducing hydrogenase large subunit | Methanogenesis | 0 | 0.688197515 | 0 | 59.0724736 | 0 | 17.51997426 | 0 | 19.20054171 | 5.886033854 | 6.844496495 | 35.01806146 | 0 | 0 | 1.782542193 | 0 | 74.16741488 |
| K14127 | mvhD, vhuD, vhcD | F420-non-reducing hydrogenase iron-sulfur subunit | Methanogenesis | 0 | 0 | 0 | 1.533401328 | 0 | 3.306433095 | 0 | 8.998440549 | 3.11292675 | 4.837037425 | 8.012341057 | 0 | 2.85442967 | 0 | 0 | 7.883421251 |
| K14128 | mvhG, vhuG, vhcG | F420-non-reducing hydrogenase small subunit | Methanogenesis | 0 | 0 | 0 | 16.8417022 | 0 | 4.217542852 | 0 | 16.5912764 | 2.240227408 | 2.632016505 | 17.47209128 | 0 | 0 | 1.234623058 | 0 | 152.8149811 |
| K16176 | mtmB | methylamine---corrinoid protein Co-methyltransferase | Methanogenesis | 0 | 3.07765232 | 0 | 4.758629898 | 0 | 0.308660874 | 0 | 14.9610243 | 0.459228606 | 0.515023989 | 9.241688277 | 0 | 0 | 0.446780409 | 0 | 57.80256874 |
| K16177 | mtmC | monomethylamine corrinoid protein | Methanogenesis | 0 | 2.095550421 | 0 | 0 | 0 | 0 | 0 | 1.754420004 | 0 | 0 | 0.570140443 | 0 | 0 | 0 | 0 | 15.66550194 |
| K16178 | mtbB | dimethylamine---corrinoid protein Co-methyltransferase | Methanogenesis | 0 | 0.541088965 | 0 | 0.230935402 | 0 | 0.233892123 | 0 | 10.08104246 | 0 | 0 | 0 | 0 | 0 | 0 | 0 | 5.415389284 |
| K16179 | mtbC | dimethylamine corrinoid protein | Methanogenesis | 0 | 0 | 0 | 0.131129344 | 0 | 0 | 0 | 3.50959422 | 0 | 0 | 0 | 0 | 0 | 0 | 0 | 9.106422303 |
| K00362 | nirB | nitrite reductase (NADH) large subunit | Nitrogen Metabolism | 0 | 13.09144368 | 0.631649475 | 21.29066397 | 0 | 9.122997069 | 0 | 31.083381 | 3.246520392 | 19.63228805 | 19.22249063 | 1.755735555 | 17.07203961 | 5.342958215 | 2.913428973 | 64.30457537 |
| K00363 | nirD | nitrite reductase (NADH) small subunit | Nitrogen Metabolism | 7.514794729 | 3.947396339 | 3.223778853 | 7.896446137 | 2.220374795 | 1.560897547 | 3.220621563 | 0.639560143 | 4.291908913 | 4.337796464 | 7.126769084 | 10.7280945 | 0 | 5.076339482 | 8.17744739 | 2.214240923 |
| K00366 | nirA | ferredoxin-nitrite reductase | Nitrogen Metabolism | 755.1387687 | 354.9359413 | 909.96661 | 899.248998 | 1096.28126 | 372.5317945 | 765.6851037 | 182.8455906 | 1100.872292 | 752.3007024 | 714.3288928 | 1201.697957 | 790.1779756 | 154.3344064 | 1985.890639 | 1732.509553 |
| K00367 | narB | ferredoxin-nitrate reductase | Nitrogen Metabolism | 75.53178426 | 75.39836916 | 21.13410854 | 37.36045153 | 94.85479221 | 84.60089709 | 28.11791433 | 13.73081235 | 55.81596555 | 86.16473113 | 22.71370241 | 93.39105693 | 170.8341165 | 9.306187443 | 133.7961193 | 2.190748266 |
| K00368 | nirK | nitrite reductase (NO-forming) | Nitrogen Metabolism | 499.2158899 | 266.8361988 | 127.020845 | 309.768713 | 92.60966252 | 345.526717 | 77.45061982 | 450.8301601 | 532.883387 | 103.6564309 | 144.4663077 | 251.5645789 | 89.08205383 | 384.3180725 | 116.7947059 | 112.886256 |
| K00370 | narG, narZ, nxrA | nitrate reductase / nitrite oxidoreductase, alpha subunit | Nitrogen Metabolism | 49.72226378 | 60.63956054 | 56.80758624 | 281.4850625 | 3.734763028 | 83.41621943 | 2.958097097 | 228.0998435 | 29.32224315 | 50.03567634 | 443.723555 | 2.164053395 | 70.79972107 | 57.55572088 | 53.86992357 | 148.0133345 |
| K00371 | narH, narY, nxrB | nitrate reductase / nitrite oxidoreductase, beta subunit | Nitrogen Metabolism | 72.86180153 | 19.33256931 | 20.71814008 | 18.69452413 | 2.73694483 | 25.05122547 | 4.556559305 | 36.47499556 | 5.399554271 | 6.801492674 | 115.5565805 | 0.601081251 | 15.80583392 | 25.59385189 | 68.44168439 | 76.24786196 |
| K00372 | nasA | assimilatory nitrate reductase catalytic subunit | Nitrogen Metabolism | 58.71130562 | 54.50865663 | 52.165169 | 187.4708296 | 300.5620704 | 146.4481505 | 521.4970032 | 171.1845512 | 494.695847 | 83.63303773 | 47.17460853 | 596.5830982 | 14.7623307 | 181.6482077 | 113.2610101 | 46.61458027 |
| K00374 | narI, narV | nitrate reductase gamma subunit | Nitrogen Metabolism | 0 | 42.77411822 | 1.547230701 | 18.01520375 | 0 | 4.485740921 | 0 | 3.168959805 | 1.410879377 | 7.01703367 | 23.32981135 | 0.578080621 | 1.929290445 | 8.848581357 | 1.138059302 | 1.891930295 |
| K00376 | nosZ | nitrous-oxide reductase | Nitrogen Metabolism | 374.4613737 | 711.8439814 | 1332.937799 | 987.9136692 | 596.0026809 | 521.6498002 | 275.1958336 | 136.5092746 | 534.9614447 | 387.4242413 | 156.2452274 | 206.7524395 | 661.4452871 | 126.2362618 | 969.7684604 | 915.2464395 |
| K02164 | norE | nitric oxide reductase NorE protein | Nitrogen Metabolism | 0 | 0.332998957 | 0 | 0 | 0 | 0 | 0 | 0 | 0.339250487 | 0 | 0 | 0 | 0 | 0 | 0 | 0 |
| K02305 | norC | nitric oxide reductase subunit C | Nitrogen Metabolism | 2.146336839 | 3.637740491 | 0 | 26.67306262 | 5.753597455 | 2.899681212 | 0.224156229 | 15.54007433 | 4.700839665 | 2.561430743 | 4.972819729 | 1.414822025 | 0 | 10.1708015 | 0 | 3.030680714 |
| K02448 | norD | nitric oxide reductase NorD protein | Nitrogen Metabolism | 0 | 2.117528697 | 0 | 1.143667268 | 1.464505072 | 0 | 1.953801973 | 0 | 0.134219643 | 0.48623387 | 2.935430061 | 0.3062262 | 0.23882826 | 5.741798962 | 0 | 0 |
| K02567 | napA | nitrate reductase (cytochrome) | Nitrogen Metabolism | 18.51436788 | 84.67967613 | 18.0657002 | 18.19482083 | 34.00867791 | 23.71479158 | 7.312581686 | 27.84157818 | 24.62933923 | 43.11968077 | 96.98719243 | 2.359547422 | 74.76749725 | 160.6048035 | 9.664023595 | 8.608272451 |
| K02568 | napB | nitrate reductase (cytochrome), electron transfer subunit | Nitrogen Metabolism | 0 | 3.709021236 | 1.161329444 | 1.623806456 | 0.195142418 | 2.344381445 | 0.799812521 | 0 | 1.938403368 | 0.90327334 | 10.39179441 | 0.392733497 | 4.688337121 | 6.04084495 | 0.446633772 | 0 |
| K02586 | nifD | nitrogenase molybdenum-iron protein alpha chain | Nitrogen Metabolism | 4.511837651 | 1.131005818 | 0.790769971 | 0 | 0 | 0 | 0 | 4.30748114 | 0 | 1.065085654 | 0 | 0 | 0 | 0 | 0 | 0 |
| K02588 | nifH | nitrogenase iron protein NifH | Nitrogen Metabolism | 4.736233207 | 6.43508721 | 0 | 0 | 0 | 0 | 0 | 1.665007414 | 0 | 0 | 0 | 0 | 0 | 0 | 0 | 0 |
| K02591 | nifK | nitrogenase molybdenum-iron protein beta chain | Nitrogen Metabolism | 14.84954048 | 0.41457679 | 0 | 0 | 0 | 0 | 0 | 0 | 0 | 0.1636024 | 0 | 0 | 0 | 0 | 0 | 0 |
| K03385 | nrfA | nitrite reductase (cytochrome c-552) | Nitrogen Metabolism | 0 | 4.538034974 | 0 | 2.089736024 | 0 | 0.577850818 | 0 | 3.401287853 | 0 | 0.42603374 | 10.67831892 | 0 | 0 | 0 | 0 | 8.363993576 |
| K04561 | norB | nitric oxide reductase subunit B | Nitrogen Metabolism | 93.9725537 | 80.20914507 | 30.10205906 | 255.5566984 | 34.87453574 | 77.41827098 | 29.1459066 | 28.51390421 | 137.3780695 | 63.75994916 | 42.84202213 | 24.25898009 | 51.0312047 | 36.6775942 | 59.19082483 | 122.3200888 |
| K04748 | norQ | nitric oxide reductase NorQ protein | Nitrogen Metabolism | 81.34831134 | 59.67773569 | 272.3119819 | 249.2274519 | 71.58279366 | 73.21435983 | 61.95438804 | 60.19165036 | 38.91541383 | 132.7732534 | 246.0194768 | 81.60861581 | 67.72465011 | 224.6638564 | 156.941732 | 97.44137246 |
| K10535 | hao | hydroxylamine dehydrogenase | Nitrogen Metabolism | 0 | 0 | 0 | 0.415197481 | 0 | 0 | 0 | 0 | 0 | 0 | 27.34126281 | 0 | 0 | 2.144451631 | 0 | 0 |
| K10944 | pmoA-amoA | methane/ammonia monooxygenase subunit A | Nitrogen Metabolism | 0 | 0 | 0 | 0 | 0 | 0 | 0 | 0 | 0 | 0 | 5.678980777 | 0 | 0 | 0.455912717 | 0 | 0 |
| K10945 | pmoB-amoB | methane/ammonia monooxygenase subunit B | Nitrogen Metabolism | 0 | 0 | 0 | 0 | 0 | 0 | 0 | 0 | 0 | 0 | 3.735407575 | 0 | 0 | 1.019704293 | 0 | 0 |
| K10946 | pmoC-amoC | methane/ammonia monooxygenase subunit C | Nitrogen Metabolism | 0 | 0 | 0 | 0 | 0 | 0 | 0 | 0 | 0 | 0 | 5.009331293 | 0 | 0 | 1.452707226 | 0 | 0 |
| K15864 | nirS | nitrite reductase (NO-forming) / hydroxylamine reductase | Nitrogen Metabolism | 4.778826548 | 6.12380171 | 29.7853305 | 123.2617906 | 19.03365582 | 19.47525725 | 3.281823046 | 59.14860609 | 15.81246889 | 16.58590572 | 29.26401018 | 6.605146532 | 24.5585148 | 173.2277531 | 12.07800557 | 9.122648943 |
| K15876 | nrfH | cytochrome c nitrite reductase small subunit | Nitrogen Metabolism | 0 | 1.40927986 | 0 | 1.268561045 | 0 | 0 | 0 | 9.118498769 | 0 | 0 | 9.39567489 | 0 | 0 | 0 | 0 | 30.74388728 |
| K00380 | cysJ | sulfite reductase (NADPH) flavoprotein alpha-component | Sulfur Metabolism | 1.244544971 | 65.00548557 | 4.81650141 | 18.501792 | 9.227785954 | 55.06288384 | 1.774778387 | 22.02483438 | 57.62789899 | 21.76149908 | 26.20431839 | 18.97178101 | 31.17337043 | 23.82895863 | 8.840015352 | 1.102540233 |
| K00381 | cysI | sulfite reductase (NADPH) hemoprotein beta-component | Sulfur Metabolism | 11.01351466 | 54.36853962 | 8.521733582 | 47.21069565 | 4.911606896 | 31.81337762 | 3.167201672 | 31.30302597 | 35.82482897 | 7.979348542 | 62.75482559 | 25.27820261 | 35.91172084 | 20.86454686 | 3.521396032 | 10.52421498 |
| K00390 | cysH | phosphoadenosine phosphosulfate reductase | Sulfur Metabolism | 2556.022334 | 557.9740283 | 944.2447727 | 1105.608957 | 282.5122703 | 1334.566122 | 795.7778391 | 605.9592332 | 337.5308295 | 2188.179496 | 891.091225 | 857.5949311 | 1482.262621 | 376.4635242 | 700.0969089 | 5687.996835 |
| K00392 | sir | sulfite reductase (ferredoxin) | Sulfur Metabolism | 116.6802514 | 33.96898888 | 10.66937598 | 51.15216398 | 62.77691175 | 57.08135466 | 28.28758238 | 22.20015016 | 30.43276752 | 45.99921454 | 88.75922679 | 42.2827499 | 118.0005618 | 22.34059108 | 171.6716191 | 5.786124043 |
| K00394 | aprA | adenylylsulfate reductase, subunit A | Sulfur Metabolism | 0 | 5.505580118 | 0 | 21.68120662 | 0 | 5.502010591 | 0 | 24.35319424 | 5.594847403 | 33.10281746 | 6.343024348 | 0 | 1.992014358 | 0.692029532 | 0 | 49.30676369 |
| K00395 | aprB | adenylylsulfate reductase, subunit B | Sulfur Metabolism | 0 | 0.493774009 | 0 | 2.000193729 | 0 | 1.06935799 | 0 | 6.64657698 | 1.048167877 | 2.477818581 | 4.439624157 | 0 | 0 | 0 | 0 | 6.224387203 |
| K00860 | cysC | adenylylsulfate kinase | Sulfur Metabolism | 36.19983916 | 66.46556542 | 63.27268185 | 99.60735997 | 87.63543574 | 48.17608824 | 25.45760845 | 121.8198435 | 34.21405742 | 146.947292 | 110.6585821 | 121.1527044 | 313.595876 | 53.87515487 | 56.78152983 | 91.18812102 |
| K00955 | cysNC | bifunctional enzyme CysN/CysC | Sulfur Metabolism | 169.3411088 | 653.8482004 | 525.6347524 | 160.0253593 | 438.0848938 | 1015.945823 | 433.5579727 | 299.4636809 | 661.830005 | 213.1256842 | 377.89658 | 808.211558 | 434.1679405 | 70.59959657 | 105.5521183 | 123.8955916 |
| K00956 | cysN | sulfate adenylyltransferase subunit 1 | Sulfur Metabolism | 7.919445527 | 14.8909976 | 1.620909587 | 2.718321852 | 6.9544988 | 13.06173021 | 3.725714384 | 2.330052993 | 11.15898532 | 26.63716166 | 141.8756712 | 14.91772163 | 16.12579103 | 12.44133765 | 18.53997243 | 19.07517461 |
| K00957 | cysD | sulfate adenylyltransferase subunit 2 | Sulfur Metabolism | 186.1768697 | 159.1767818 | 619.7469037 | 80.66300885 | 312.2885935 | 412.5397072 | 205.4328388 | 980.1522296 | 115.1841917 | 342.0666918 | 459.5528154 | 244.4717737 | 431.213579 | 84.74387257 | 131.6585742 | 374.7962147 |
| K00958 | sat, met3 | sulfate adenylyltransferase | Sulfur Metabolism | 325.2160466 | 152.9169372 | 358.4129464 | 91.5223726 | 265.210612 | 104.7396642 | 64.82329929 | 251.8706015 | 175.2323082 | 289.9021175 | 369.9361817 | 198.9401564 | 508.5055509 | 97.36972178 | 161.207827 | 444.9504538 |
| K08357 | ttrABC | Involved in alternative thiosulfate oxidation to/from tetrathionate | Sulfur Metabolism | 0 | 0 | 0 | 0 | 0 | 0 | 0 | 0 | 0 | 0 | 0 | 0 | 0 | 0 | 0.539215276 | 0 |
| K08358 | ttrABC | Involved in alternative thiosulfate oxidation to/from tetrathionate | Sulfur Metabolism | 0 | 0 | 0 | 0 | 0 | 0 | 0 | 0 | 0 | 0 | 1.867706496 | 0 | 0 | 0.254351468 | 0 | 0.368359914 |
| K11180 | dsrA | dissimilatory sulfite reductase alpha subunit | Sulfur Metabolism | 0 | 1.413821963 | 0 | 3.531105169 | 0 | 2.047790275 | 0 | 22.3543788 | 1.533289156 | 3.621955069 | 5.402487334 | 0 | 2.049455011 | 0.398297554 | 0 | 40.87546553 |
| K11181 | dsrB | dissimilatory sulfite reductase beta subunit | Sulfur Metabolism | 0 | 0.278999126 | 0 | 6.723388328 | 0 | 2.059350862 | 0 | 9.739107414 | 0.610088952 | 4.201269771 | 1.272770037 | 0 | 1.28360728 | 1.858719664 | 0 | 26.5595724 |
| K13811 | PAPSS | 3''-phosphoadenosine 5''-phosphosulfate synthase | Sulfur Metabolism | 0 | 0 | 0 | 0 | 0 | 0 | 0 | 0 | 0 | 0 | 0 | 0 | 0 | 0 | 0 | 0 |
| K16937 | doxAD | Involved in alternative thiosulfate oxidation to/from tetrathionate | Sulfur Metabolism | 212.0600823 | 241.3349023 | 983.4124166 | 347.1954301 | 200.3984306 | 196.6587513 | 135.096649 | 311.9443485 | 240.9801346 | 647.4423565 | 341.2398583 | 208.2911023 | 804.165586 | 838.6432554 | 814.8520701 | 748.0289166 |
| K16950 | asrA | anaerobic sulfite reductase subunit A | Sulfur Metabolism | 0 | 0 | 0 | 0 | 0 | 0.765898251 | 0 | 0 | 0 | 0 | 0 | 0 | 0 | 0 | 0 | 0.229037824 |
| K17218 | sqr | sulfide:quinone oxidoreductase | Sulfur Metabolism | 1252.411097 | 1238.182946 | 651.1193474 | 293.5765512 | 2395.18072 | 2904.651888 | 659.2153144 | 588.7762853 | 1433.841361 | 444.3756 | 610.2538748 | 713.1838306 | 397.5795833 | 784.4206312 | 849.1474778 | 1725.555278 |
| K17222 | soxA | L-cysteine S-thiosulfotransferase | Sulfur Metabolism | 7.055756088 | 9.667743632 | 2.244266626 | 0.452336751 | 0 | 8.118619364 | 0.657764708 | 4.315538894 | 2.660984264 | 3.570522708 | 7.126769084 | 0.688359012 | 7.378049886 | 1.871811384 | 18.36484996 | 6.340387695 |
| K17223 | soxX | L-cysteine S-thiosulfotransferase | Sulfur Metabolism | 4.55385908 | 14.38486962 | 2.515772673 | 0 | 0.582202213 | 7.042279834 | 0.894348579 | 0.786458301 | 1.001856349 | 3.72779457 | 1.324290285 | 0.728377843 | 0 | 1.412146577 | 7.033825263 | 3.221531579 |
| K17224 | soxB | S-sulfosulfanyl-L-cysteine sulfohydrolase | Sulfur Metabolism | 2.932002554 | 77.81752107 | 2.053426659 | 1.020099788 | 1.360462649 | 4.14445886 | 1.61177607 | 2.003702427 | 4.424601941 | 3.246214403 | 0 | 3.141087314 | 0 | 2.052915915 | 3.232536285 | 4.142932951 |
| K17225 | soxC | sulfane dehydrogenase subunit SoxC | Sulfur Metabolism | 3.487122321 | 62.12056159 | 1.483501702 | 0 | 0.872402309 | 11.23786887 | 1.393392314 | 5.066356835 | 4.676995521 | 1.619965417 | 0 | 1.976224356 | 0 | 3.092912615 | 2.084042543 | 0 |
| K17226 | soxY | sulfur-oxidizing protein SoxY | Sulfur Metabolism | 1.180809657 | 13.64216686 | 0.680026332 | 3.623028882 | 0 | 5.312552541 | 0.728045693 | 0 | 1.667410618 | 0.398390216 | 2.927755103 | 0.585247073 | 0 | 3.894097799 | 0.978167143 | 0.636925992 |
| K17227 | soxZ | sulfur-oxidizing protein SoxZ | Sulfur Metabolism | 5.430983751 | 14.63278589 | 2.842425228 | 0.148956944 | 2.362535815 | 5.281116784 | 2.327670734 | 2.346353205 | 1.251201128 | 3.023147738 | 4.04204838 | 1.209034365 | 0 | 1.898028651 | 6.206597683 | 0.724478707 |
| K19713 | tsdAB | Involved in alternative thiosulfate oxidation to/from tetrathionate | Sulfur Metabolism | 0 | 0 | 0 | 0.869350618 | 0 | 0.88188821 | 0 | 0.913473741 | 0 | 1.526557361 | 2.990617173 | 0 | 0 | 0 | 0 | 0.722732145 |
| K08355 | aoxA | arsenite oxidase small subunit | Arsenic Metabolism | 43.30785483 | 52.43905401 | 44.41969233 | 26.08781564 | 9.992905301 | 21.69022038 | 18.24862471 | 34.89199715 | 20.72033921 | 63.91808214 | 48.0997198 | 44.46265447 | 89.6988686 | 60.22239148 | 42.94974308 | 58.60207047 |
| K08356 | aoxB | arsenite oxidase large subunit | Arsenic Metabolism | 472.6332576 | 164.6829125 | 243.2413236 | 448.0595673 | 105.8289467 | 117.6415797 | 91.52737187 | 311.7357858 | 856.0822623 | 680.5459046 | 171.2463291 | 207.4991102 | 1080.746653 | 1042.794819 | 104.236451 | 114.759114 |
| K00055 | E1.1.1.90 | aryl-alcohol dehydrogenase | Hydrocarbon Degredation | 342.570278 | 61.94988861 | 64.14957713 | 107.333409 | 31.63801257 | 127.6070309 | 61.79229603 | 110.0356674 | 163.4135857 | 161.4908293 | 29.47741407 | 54.31247613 | 92.99012679 | 51.83141621 | 89.97336776 | 57.39770208 |
| K00446 | dmpB, xylE | catechol 2,3-dioxygenase | Hydrocarbon Degredation | 52.8290556 | 137.8652317 | 163.4790741 | 139.8207796 | 15.686743 | 42.88344853 | 70.34911013 | 45.95128322 | 37.112708 | 55.66176603 | 59.08742019 | 74.99905044 | 145.7421819 | 137.2388152 | 151.9678564 | 280.6657162 |
| K00462 | bphC | biphenyl-2,3-diol 1,2-dioxygenase | Hydrocarbon Degredation | 0.392284385 | 0 | 0.717956067 | 0.598840292 | 0.862610247 | 0 | 0.989119553 | 0 | 0.297664972 | 0 | 0 | 0 | 0.473835041 | 0 | 0 | 0 |
| K01055 | pcaD | 3-oxoadipate enol-lactonase | Hydrocarbon Degredation | 0.90149935 | 16.66891459 | 0.920222089 | 4.29720185 | 7.350950521 | 8.204250547 | 2.640308649 | 8.411463598 | 3.477742145 | 2.688130228 | 11.90067906 | 5.358954252 | 1.146375176 | 6.731241038 | 0.49187682 | 11.59197584 |
| K01617 | dmpH, xylI, nahK | 2-oxo-3-hexenedioate decarboxylase | Hydrocarbon Degredation | 0 | 0.552925411 | 0.506804035 | 0 | 0 | 1.222123417 | 0 | 0 | 0 | 0 | 2.716658577 | 0 | 0 | 0 | 0 | 0 |
| K01666 | mhpE | 4-hydroxy 2-oxovalerate aldolase | Hydrocarbon Degredation | 27.20376562 | 3.876429411 | 21.47611323 | 6.661510909 | 25.67119472 | 7.872495614 | 3.688022118 | 6.380648022 | 21.55455204 | 25.99826266 | 4.874708104 | 16.1309199 | 11.3790419 | 17.85987891 | 15.50896664 | 9.562686642 |
| K01821 | praC, xylH | 4-oxalocrotonate tautomerase | Hydrocarbon Degredation | 7.772520535 | 14.04484499 | 10.63047767 | 14.04612479 | 12.04522295 | 17.95031426 | 7.911676132 | 5.789053447 | 8.414804927 | 9.193945538 | 15.42281815 | 4.109459172 | 6.28984033 | 7.126505605 | 15.20871612 | 8.840641132 |
| K01856 | catB | muconate cycloisomerase | Hydrocarbon Degredation | 64.23062626 | 81.26016032 | 14.05746523 | 33.5527962 | 13.77052797 | 20.22876647 | 18.06374584 | 22.76096058 | 40.37079244 | 20.59481168 | 4.287343692 | 30.45951948 | 7.342488566 | 23.04679002 | 18.84187996 | 12.29755491 |
| K02554 | mhpD | 2-keto-4-pentenoate hydratase | Hydrocarbon Degredation | 3.261076659 | 2.034669716 | 1.491488545 | 6.963305077 | 22.50539983 | 5.379521096 | 34.50616041 | 18.15341855 | 15.36374869 | 4.136941482 | 19.66441 | 24.72794368 | 1.438764564 | 9.81405336 | 9.842412739 | 0 |
| K03381 | catA | catechol 1,2-dioxygenase | Hydrocarbon Degredation | 0.305394158 | 0 | 0 | 0 | 0.822730488 | 0 | 0 | 0 | 0 | 0 | 0 | 0 | 0.558767603 | 0 | 0.455849124 | 0 |
| K04102 | pht5 | 4,5-dihydroxyphthalate decarboxylase | Hydrocarbon Degredation | 25.5847874 | 24.64512379 | 25.3562583 | 32.00255732 | 9.495905302 | 10.05047993 | 5.211246614 | 19.80006321 | 18.74507708 | 13.73398 | 46.23969368 | 10.03507304 | 42.59196571 | 11.33895593 | 13.47818045 | 161.9906264 |
| K04112 | bcrC, badD | benzoyl-CoA reductase subunit C | Hydrocarbon Degredation | 0 | 0 | 0 | 3.733920119 | 0 | 0 | 0 | 0 | 0.379867472 | 0 | 22.46411735 | 0 | 0 | 2.103291729 | 0 | 5.436332041 |
| K04113 | bcrB, badE | benzoyl-CoA reductase subunit B | Hydrocarbon Degredation | 0 | 0.993784739 | 0 | 26.3623443 | 0 | 2.80235388 | 0 | 0.30121453 | 0 | 0.639050349 | 10.47818507 | 0 | 0 | 11.39583653 | 0 | 4.965847411 |
| K04114 | bcrA, badF | benzoyl-CoA reductase subunit A | Hydrocarbon Degredation | 0 | 0 | 0 | 1.115489623 | 0 | 0 | 0 | 0.836528736 | 0 | 0 | 1.089724748 | 0 | 0 | 0 | 0 | 0 |
| K04116 | aliA | cyclohexanecarboxylate-CoA ligase | Hydrocarbon Degredation | 22.11515995 | 4.691249763 | 5.400752961 | 27.16788475 | 12.10489112 | 5.489305243 | 5.730939462 | 12.63127915 | 5.266307587 | 5.044113348 | 7.417789158 | 3.803477141 | 10.83464097 | 16.82945574 | 8.615797998 | 10.81125864 |
| K04117 | aliB | cyclohexanecarboxyl-CoA dehydrogenase | Hydrocarbon Degredation | 0 | 0.435798048 | 0 | 6.601457765 | 0 | 0 | 1.192468687 | 0.431553889 | 1.90170701 | 4.103576988 | 29.27755103 | 1.838600141 | 3.747371805 | 10.69360038 | 0.300115117 | 4.475083477 |
| K05784 | benC-xylZ | benzoate/toluate 1,2-dioxygenase reductase component | Hydrocarbon Degredation | 0 | 0.593595132 | 0 | 0.132184465 | 0 | 0 | 0 | 0.330302482 | 0 | 0 | 0 | 0 | 0 | 0 | 0 | 0 |
| K07104 | catE | catechol 2,3-dioxygenase | Hydrocarbon Degredation | 28.84796419 | 102.0547364 | 148.9359756 | 634.162191 | 55.04879071 | 112.6468039 | 125.9830841 | 109.6947329 | 99.17797995 | 73.29072317 | 49.55953239 | 92.67214561 | 26.81444343 | 452.1499955 | 39.96503893 | 251.0972612 |
| K07535 | badH | 2-hydroxycyclohexanecarboxyl-CoA dehydrogenase | Hydrocarbon Degredation | 5.778490281 | 10.23569928 | 4.593672634 | 21.79648105 | 13.77271182 | 13.514137 | 13.61267555 | 16.07117803 | 14.47583117 | 9.588234212 | 50.57910467 | 15.72852218 | 3.181743071 | 15.63733124 | 4.543772246 | 21.01717568 |
| K07536 | badI | 2-ketocyclohexanecarboxyl-CoA hydrolase | Hydrocarbon Degredation | 0 | 0 | 0 | 0 | 0 | 0 | 0 | 0 | 0 | 0 | 0 | 0 | 0 | 0 | 0.136829871 | 0 |
| K07537 | dch | cyclohexa-1,5-dienecarbonyl-CoA hydratase | Hydrocarbon Degredation | 0 | 0 | 0 | 3.008420941 | 0 | 0 | 0 | 0 | 0 | 0 | 1.945376845 | 0 | 0.336530677 | 0.117297744 | 0 | 3.717866925 |
| K07538 | had | 6-hydroxycyclohex-1-ene-1-carbonyl-CoA dehydrogenase | Hydrocarbon Degredation | 0 | 0 | 0 | 0.892390712 | 0 | 0 | 0 | 10.29406884 | 0 | 0 | 10.57297106 | 0 | 0 | 0 | 0 | 31.14315854 |
| K07539 | oah | 6-oxocyclohex-1-ene-carbonyl-CoA hydrolase | Hydrocarbon Degredation | 0 | 0 | 0 | 0.404966752 | 0 | 0 | 0 | 0.400140964 | 0 | 0 | 0 | 0 | 0 | 0 | 0 | 0.538881916 |
| K07543 | bbsE | benzylsuccinate CoA-transferase BbsE subunit | Hydrocarbon Degredation | 0 | 0 | 0 | 0 | 0 | 0 | 0 | 0 | 0 | 0 | 0 | 0 | 0 | 0.354474849 | 0 | 0 |
| K07544 | bbsF | benzylsuccinate CoA-transferase BbsF subunit | Hydrocarbon Degredation | 0 | 0 | 0 | 0.188185748 | 0 | 0.543778344 | 0 | 0 | 1.235495736 | 0 | 3.679023451 | 0 | 0 | 0.305221665 | 0 | 0 |
| K07546 | bbsH | E-phenylitaconyl-CoA hydratase | Hydrocarbon Degredation | 0 | 0 | 0 | 0 | 0 | 0 | 0 | 0 | 0 | 0 | 0 | 0.578080621 | 0 | 0 | 0 | 0 |
| K07548 | bbsD | 2-hydroxy(phenyl)methyl-succinyl-CoA dehydrogenase BbsD subunit | Hydrocarbon Degredation | 0 | 0 | 0 | 0 | 0 | 0 | 0.492142491 | 0 | 0 | 0.377270851 | 0 | 0.129047133 | 0 | 0 | 0 | 0 |
| K07549 | bbsA | benzoylsuccinyl-CoA thiolase BbsA subunit | Hydrocarbon Degredation | 0 | 0 | 0 | 0 | 0 | 0 | 0 | 0 | 0 | 0 | 0 | 0 | 0 | 0 | 0 | 0 |
| K08690 | bphB | cis-2,3-dihydrobiphenyl-2,3-diol dehydrogenase | Hydrocarbon Degredation | 2.698859565 | 1.929364486 | 7.664779513 | 0 | 0 | 0 | 0 | 1.059529184 | 0 | 0 | 0 | 0 | 0 | 0 | 0 | 0 |

| | | | | | | | | | | | | | | | | | | | |
|---|---|---|---|---|---|---|---|---|---|---|---|---|---|---|---|---|---|---|---|
| K10216 | dmpD, xylF | 2-hydroxymuconate-semialdehyde hydrolase | Hydrocarbon Degredation | 2.766241986 | 1.272967201 | 3.875100202 | 3.590980816 | 1.000255647 | 1.542912739 | 1.033630256 | 2.313260234 | 2.901629252 | 2.835710574 | 11.03438926 | 3.274672255 | 4.244045695 | 4.446274521 | 8.68184175 | 1.709384592 |
| K10217 | dmpC, xylG, praB | aminomuconate-semialdehyde/2-hydroxymuconate-6-semialdehyde dehydrogenase | Hydrocarbon Degredation | 0.369118221 | 12.93937956 | 2.092014195 | 8.339350403 | 0 | 6.379477913 | 5.13365387 | 15.05745568 | 22.46040802 | 20.37123356 | 58.34754783 | 20.15829628 | 184.3751374 | 10.46525645 | 1.963284265 | 1.975309645 |
| K10622 | cmtD, dhbB | HCOMODA/2-hydroxy-3-carboxy-muconic semialdehyde decarboxylase | Hydrocarbon Degredation | 0 | 0 | 0 | 2.01243609 | 0.450137584 | 0 | 0 | 0 | 1.395771934 | 0 | 0 | 2.142293439 | 0 | 0 | 0 | 0 |
| K11311 | antC | anthranilate 1,2-dioxygenase reductase component | Hydrocarbon Degredation | 0 | 0 | 0 | 0.295237627 | 0 | 0 | 0 | 0 | 0 | 0 | 0 | 0 | 0 | 0 | 0 | 0 |
| K14578 | nagAb, ndoA, nbzAb | naphthalene 1,2-dioxygenase ferredoxin component | Hydrocarbon Degredation | 0 | 1.114489078 | 0 | 0 | 0 | 1.853866149 | 0 | 0 | 0.112531808 | 0.74555787 | 1.145448078 | 0 | 0 | 0 | 0 | 0 |
| K14581 | nagAa, ndoR, nbzAa | naphthalene 1,2-dioxygenase ferredoxin reductase component | Hydrocarbon Degredation | 1.659396419 | 0.987546917 | 0 | 0 | 0.687283018 | 0 | 0 | 1.59668967 | 0 | 0.877126599 | 0 | 0 | 0 | 0 | 0 | 0 |
| K14584 | nahD | 2-hydroxychromene-2-carboxylate isomerase | Hydrocarbon Degredation | 0 | 0 | 0 | 0 | 0 | 0 | 0 | 0 | 0 | 0.196939756 | 0.242885039 | 0 | 0 | 0 | 0 | 0 |
| K14585 | nahE | trans-o-hydroxybenzylidenepyruvate hydratase-aldolase | Hydrocarbon Degredation | 8.915874679 | 7.86510659 | 24.36427376 | 5.378700801 | 0.216758219 | 2.297694752 | 4.47072101 | 13.10681745 | 13.25269305 | 0.648313914 | 0 | 3.817356832 | 0.485486645 | 3.774750033 | 1.392448472 | 0 |
| K14727 | pcaL | 3-oxoadipate enol-lactonase / 4-carboxymuconolactone decarboxylase | Hydrocarbon Degredation | 0 | 8.130915998 | 0.972380007 | 0.717531536 | 0 | 3.676115789 | 1.181850485 | 0 | 1.709244909 | 0 | 0.152572947 | 1.79982765 | 0 | 0.337163237 | 0 | 0 |
| K15750 | bphAb, bphA2, bphE | biphenyl 2,3-dioxygenase subunit beta | Hydrocarbon Degredation | 0 | 0 | 0.202325587 | 0 | 0 | 0 | 0 | 0 | 0 | 0 | 0 | 0 | 0 | 0 | 0 | 0 |
| K15756 | carC | 2-hydroxy-6-oxo-6-(2'-aminophenyl)hexa-2,4-dienoate hydrolase | Hydrocarbon Degredation | 0 | 1.409879968 | 0 | 0 | 0 | 0 | 0 | 1.469912807 | 1.303599186 | 1.737881477 | 0 | 0.769727991 | 1.053636181 | 0.29229909 | 0 | 8.021075914 |
| K15760 | tmoA, tbuA1, touA | toluene monooxygenase system protein A | Hydrocarbon Degredation | 0 | 0 | 0 | 0 | 0 | 0 | 0 | 0.185446852 | 0 | 0 | 0 | 0 | 0 | 0 | 0 | 0 |
| K15762 | tmoC, tbuB, touC | toluene monooxygenase system ferredoxin subunit | Hydrocarbon Degredation | 0 | 0 | 0 | 0 | 0 | 0 | 0 | 0 | 0 | 0 | 0 | 0 | 0 | 0 | 0 | 0 |
| K15763 | tmoD, tbuV, touD | toluene monooxygenase system protein D | Hydrocarbon Degredation | 0 | 0 | 0 | 0 | 0 | 0 | 0 | 0 | 0 | 0 | 0 | 0 | 0 | 0 | 0 | 0 |
| K15764 | tmoE, tbuA2, touE | toluene monooxygenase system protein E | Hydrocarbon Degredation | 0 | 0 | 0 | 0 | 0 | 0 | 0 | 0 | 0 | 0 | 0.666627166 | 0 | 0 | 0 | 0 | 0 |
| K16246 | dmpP, poxF, tomA5 | phenol/toluene 2-monooxygenase (NADH) P5/A5 | Hydrocarbon Degredation | 0 | 0 | 0 | 0 | 0 | 0 | 0 | 0 | 0 | 0.232640755 | 1.021950056 | 0 | 0.342365955 | 0 | 0 | 0.428601515 |
| K16268 | todC2, bedC2, tcbAb | benzene/toluene/chlorobenzene dioxygenase subunit beta | Hydrocarbon Degredation | 0 | 0 | 0 | 0 | 0 | 0 | 0 | 0 | 0 | 0.211291759 | 0 | 0.345437743 | 0 | 0 | 0 | 0 |
| K16269 | todD, tcbB | cis-1,2-dihydrobenzene-1,2-diol/chlorobenzene dihydrodiol dehydrogenase | Hydrocarbon Degredation | 0 | 0 | 0 | 0 | 0 | 0 | 0 | 0 | 0 | 0 | 0 | 0 | 0 | 0 | 0 | 0 |
| K18087 | bphAc, bphA3, bphF | biphenyl 2,3-dioxygenase ferredoxin component | Hydrocarbon Degredation | 0 | 0 | 0 | 0 | 0 | 0 | 0 | 0 | 0 | 0 | 0 | 0 | 0 | 0 | 0 | 0 |
| K18248 | andAb | anthranilate 1,2-dioxygenase ferredoxin component | Hydrocarbon Degredation | 0 | 0.179334913 | 0 | 0 | 0 | 0 | 0 | 0 | 0.11462869 | 0 | 0 | 0.257507919 | 0 | 0.172595639 | 0 | 0 |
| K18249 | andAa | anthranilate 1,2-dioxygenase ferredoxin reductase component | Hydrocarbon Degredation | 0 | 2.542317237 | 0 | 0 | 0 | 0 | 0 | 0 | 0.670038887 | 0 | 0 | 0 | 0 | 0 | 0 | 0 |
| K18364 | bphH, xylJ, tesE | 2-oxopent-4-enoate/cis-2-oxohex-4-enoate hydratase | Hydrocarbon Degredation | 0 | 0 | 0 | 0 | 0 | 0 | 0 | 0 | 0.105760588 | 0 | 1.019816017 | 0 | 0 | 0 | 0 | 0 |
| K19185 | kdhA | 6-hydroxypseudooxynicotine dehydrogenase subunit alpha | Hydrocarbon Degredation | 0 | 0 | 0 | 0 | 0 | 0 | 0.338495703 | 0 | 0 | 0 | 0 | 0 | 0 | 0 | 0 | 0 |
| K19186 | kdhB | 6-hydroxypseudooxynicotine dehydrogenase subunit beta | Hydrocarbon Degredation | 0 | 0 | 0 | 0 | 0 | 0 | 0 | 0 | 0 | 0 | 0 | 0 | 0 | 0 | 0 | 0 |
| K19187 | kdhL | 6-hydroxypseudooxynicotine dehydrogenase subunit gamma | Hydrocarbon Degredation | 0 | 0 | 0 | 0 | 0 | 0 | 0 | 0 | 0 | 0 | 0 | 0 | 0 | 0 | 0 | 0 |
| K19515 | bamB | benzoyl-CoA reductase subunit BamB | Hydrocarbon Degredation | 0 | 0 | 0 | 0 | 0 | 0 | 0 | 0 | 0 | 0 | 0 | 0 | 0 | 0 | 0 | 0 |
| K19516 | bamC | benzoyl-CoA reductase subunit BamC | Hydrocarbon Degredation | 0 | 0 | 0 | 0.142499012 | 0 | 0 | 0 | 0.426767722 | 0 | 0 | 0 | 0 | 0 | 0 | 0 | 2.761366452 |
| K19818 | ndhA | nicotine dehydrogenase subunit A | Hydrocarbon Degredation | 0 | 0 | 0 | 0 | 0 | 0 | 0 | 0.425195421 | 0 | 0 | 0 | 0 | 0 | 0 | 0 | 0 |
| K19819 | ndhB | nicotine dehydrogenase subunit B | Hydrocarbon Degredation | 0 | 0 | 0 | 0 | 0 | 0 | 0 | 0 | 0 | 0 | 0.703420379 | 0 | 0 | 0 | 0 | 0 |
| K19820 | ndhC | nicotine dehydrogenase subunit C | Hydrocarbon Degredation | 0 | 0 | 0 | 0 | 0 | 0 | 0 | 0 | 0 | 0 | 0 | 0 | 0 | 0 | 0 | 0 |
| K20155 | mao | 4-methylaminobutanoate oxidase (methylamine-forming) | Hydrocarbon Degredation | 0 | 0 | 0 | 0 | 0 | 0 | 0 | 0 | 0 | 0 | 0 | 0 | 0 | 0 | 0 | 0 |
| K20170 | nicA1 | nicotine oxidoreductase | Hydrocarbon Degredation | 2.068144354 | 0 | 0 | 0 | 0 | 0 | 0 | 0 | 1.440821125 | 2.135505035 | 1.948323335 | 0 | 3.3471115 | 0.855847582 | 0 | 0 |

*Values represent transcripts per million (TPM)

***Table S4***

| Bin ID | Operon Order on Contigs | Form |
| --- | --- | --- |
| Halobacteriales.2 | coxSL, coxLM | Infer form II |
| Halobacteriales.3 | none | form II or mis-annotated |
| Haloarculaceae.1 | coxLM | form II |
| Haloarculaceae.2 | coxGSL, coxMS, coxLM | Infer both |
| Haloarculaceae.5 | coxSLM, coxMSL, coxGMSL | form I, form II |
| Halanerobiales | coxMS, coxMSMS, coxMS, coxMS | Infer form I missing coxL |

***Table S5***

| Query | Reference | ANI Estimate | Matches | Total |
|---|---|---|---|---|
| PCN9.gz_assembly | Haloferacaceae.3 | 75.5033 | 51 | 981 |
| Haloferacaceae.2 | Haloarculaceae.1 | 75.5864 | 54 | 416 |
| Haloarculaceae.4 | Halovenus | 75.5912 | 76 | 678 |
| WSH3.gz_assembly | Haloferacaceae.2 | 75.6237 | 81 | 1077 |
| Haloferacaceae.2 | WSH3.gz_assembly | 75.6309 | 59 | 416 |
| WSH3.gz_assembly | Haloferacaceae.3 | 75.6391 | 51 | 1077 |
| Haloarculaceae.3 | Haloferacaceae.1 | 75.6425 | 52 | 604 |
| Haloferacaceae.1 | Haloarculaceae.3 | 75.6458 | 54 | 713 |
| Halovenus | WSA2.gz_assembly | 75.6842 | 58 | 605 |
| Haloferacaceae.3 | Haloarculaceae.4 | 75.6976 | 57 | 347 |
| Haloarculaceae.4 | Haloferacaceae.3 | 75.7592 | 57 | 678 |
| Haloarculaceae.1 | Haloferacaceae.1 | 75.7632 | 92 | 953 |
| Halovenus | Haloarculaceae.2 | 75.7656 | 56 | 605 |
| Haloarculaceae.2 | Halobacteriales.2 | 75.7952 | 66 | 663 |
| Halobacteriales.2 | Haloarculaceae.2 | 75.8023 | 68 | 686 |
| Haloarculaceae.2 | Halovenus | 75.8058 | 61 | 663 |
| Halobacteriales.2 | Haloarculaceae.1 | 75.8343 | 65 | 686 |
| WSA2.gz_assembly | Haloarculaceae.3 | 75.8417 | 118 | 1248 |
| Haloarculaceae.3 | WSA2.gz_assembly | 75.8542 | 121 | 604 |
| Haloarculaceae.1 | Halobacteriales.2 | 75.8638 | 64 | 953 |
| WSA2.gz_assembly | Halovenus | 75.8747 | 67 | 1248 |
| Haloarculaceae.5 | Halobacteriales.2 | 75.8963 | 77 | 824 |
| Halovenus | Haloarculaceae.4 | 75.9099 | 64 | 605 |
| Haloferacaceae.3 | WSA2.gz_assembly | 75.9147 | 84 | 347 |
| WSA2.gz_assembly | Haloferacaceae.3 | 75.9204 | 93 | 1248 |
| PCN9.gz_assembly | Halovenus | 75.9496 | 75 | 981 |
| Halobacteriales.2 | Haloarculaceae.5 | 75.9543 | 87 | 686 |
| WSA2.gz_assembly | Halobacteriales.2 | 75.9709 | 104 | 1248 |
| Haloferacaceae.2 | Haloarculaceae.5 | 75.9789 | 57 | 416 |
| Haloarculaceae.1 | Haloferacaceae.2 | 75.9804 | 54 | 953 |
| Halobacteriales.2 | Haloferacaceae.1 | 75.9858 | 60 | 686 |
| PCN9.gz_assembly | Halobacteriales.2 | 75.996 | 70 | 981 |
| Haloferacaceae.2 | Haloarculaceae.4 | 76.0114 | 57 | 416 |
| Halobacteriales.2 | Haloarculaceae.4 | 76.0409 | 81 | 686 |

| | | | | |
|---|---|---|---|---|
| Haloferacaceae.1 | Halobacteriales.2 | 76.0496 | 53 | 713 |
| Haloarculaceae.4 | Halobacteriales.2 | 76.0591 | 70 | 678 |
| Halobacteriales.2 | PCN9.gz_assembly | 76.067 | 58 | 686 |
| Halobacteriales.2 | WSA2.gz_assembly | 76.0745 | 101 | 686 |
| Haloarculaceae.4 | Haloferacaceae.2 | 76.1068 | 60 | 678 |
| Haloarculaceae.3 | Haloarculaceae.4 | 76.1343 | 124 | 604 |
| PCN9.gz_assembly | Haloarculaceae.3 | 76.1655 | 107 | 981 |
| PCN9.gz_assembly | Haloarculaceae.2 | 76.2202 | 157 | 981 |
| Haloarculaceae.2 | Haloferacaceae.1 | 76.2264 | 100 | 663 |
| Haloarculaceae.5 | Haloferacaceae.2 | 76.2338 | 55 | 824 |
| Haloarculaceae.2 | PCN9.gz_assembly | 76.2443 | 159 | 663 |
| Haloferacaceae.1 | Haloarculaceae.1 | 76.2665 | 95 | 713 |
| WSH3.gz_assembly | Halobacteriales.2 | 76.2729 | 94 | 1077 |
| Haloferacaceae.1 | Haloarculaceae.2 | 76.2759 | 94 | 713 |
| PCN9.gz_assembly | Haloferacaceae.1 | 76.276 | 164 | 981 |
| Halovenus | PCN9.gz_assembly | 76.3011 | 66 | 605 |
| Haloarculaceae.3 | PCN9.gz_assembly | 76.3024 | 103 | 604 |
| Haloarculaceae.4 | Haloarculaceae.3 | 76.3097 | 119 | 678 |
| PCN9.gz_assembly | Haloferacaceae.2 | 76.3385 | 78 | 981 |
| Haloarculaceae.2 | WSA2.gz_assembly | 76.3494 | 195 | 663 |
| Halovenus | Haloarculaceae.5 | 76.3631 | 89 | 605 |
| Halobacteriales.2 | WSH3.gz_assembly | 76.3671 | 83 | 686 |
| Haloarculaceae.5 | Halovenus | 76.3789 | 92 | 824 |
| WSA2.gz_assembly | Haloarculaceae.2 | 76.4137 | 209 | 1248 |
| Haloarculaceae.5 | Haloferacaceae.1 | 76.4297 | 99 | 824 |
| Haloferacaceae.2 | PCN9.gz_assembly | 76.4777 | 61 | 416 |
| Haloferacaceae.1 | PCN9.gz_assembly | 76.5038 | 147 | 713 |
| PCN9.gz_assembly | Haloarculaceae.1 | 76.5093 | 226 | 981 |
| Halovenus | Haloarculaceae.1 | 76.5207 | 76 | 605 |
| Haloferacaceae.1 | Haloarculaceae.5 | 76.5286 | 91 | 713 |
| Haloferacaceae.1 | WSH3.gz_assembly | 76.5404 | 121 | 713 |
| WSH3.gz_assembly | Haloferacaceae.1 | 76.5728 | 136 | 1077 |
| WSH3.gz_assembly | Haloarculaceae.2 | 76.5878 | 217 | 1077 |
| Haloarculaceae.3 | Haloarculaceae.2 | 76.6097 | 124 | 604 |
| Haloarculaceae.2 | Haloarculaceae.3 | 76.6156 | 108 | 663 |
| Haloferacaceae.1 | Haloarculaceae.4 | 76.623 | 182 | 713 |

| | | | | |
|---|---|---|---|---|
| Haloarculaceae.1 | PCN9.gz_assembly | 76.6728 | 220 | 953 |
| PCN9.gz_assembly | WSH3.gz_assembly | 76.688 | 189 | 981 |
| Haloarculaceae.4 | Haloferacaceae.1 | 76.6991 | 166 | 678 |
| Haloarculaceae.1 | WSA2.gz_assembly | 76.7738 | 235 | 953 |
| Haloarculaceae.2 | WSH3.gz_assembly | 76.7878 | 201 | 663 |
| WSA2.gz_assembly | Haloarculaceae.1 | 76.8051 | 263 | 1248 |
| Haloarculaceae.1 | Haloarculaceae.3 | 76.8539 | 153 | 953 |
| WSA2.gz_assembly | Haloarculaceae.5 | 76.8577 | 265 | 1248 |
| WSH3.gz_assembly | PCN9.gz_assembly | 76.8622 | 178 | 1077 |
| Haloarculaceae.1 | Halovenus | 76.8764 | 81 | 953 |
| PCN9.gz_assembly | Haloarculaceae.5 | 76.9259 | 233 | 981 |
| Haloarculaceae.3 | Haloarculaceae.1 | 76.9321 | 154 | 604 |
| PCN9.gz_assembly | Haloarculaceae.4 | 76.9435 | 259 | 981 |
| Haloferacaceae.2 | WSA2.gz_assembly | 76.9708 | 128 | 416 |
| Haloferacaceae.1 | Haloferacaceae.3 | 76.9895 | 96 | 713 |
| Haloarculaceae.4 | PCN9.gz_assembly | 76.9934 | 249 | 678 |
| Haloarculaceae.2 | Haloarculaceae.4 | 77.0054 | 206 | 663 |
| WSA2.gz_assembly | WSH3.gz_assembly | 77.0281 | 287 | 1248 |
| Haloarculaceae.5 | WSA2.gz_assembly | 77.0404 | 254 | 824 |
| Haloarculaceae.4 | Haloarculaceae.2 | 77.0858 | 218 | 678 |
| WSA2.gz_assembly | Haloferacaceae.2 | 77.0998 | 144 | 1248 |
| WSH3.gz_assembly | WSA2.gz_assembly | 77.1226 | 281 | 1077 |
| Haloarculaceae.4 | Haloarculaceae.1 | 77.151 | 233 | 678 |
| WSH3.gz_assembly | Haloarculaceae.4 | 77.1535 | 256 | 1077 |
| Haloarculaceae.1 | Haloarculaceae.4 | 77.2056 | 233 | 953 |
| Haloarculaceae.4 | WSH3.gz_assembly | 77.2218 | 234 | 678 |
| Haloferacaceae.3 | Haloferacaceae.1 | 77.2904 | 98 | 347 |
| WSH3.gz_assembly | Haloarculaceae.3 | 77.3682 | 184 | 1077 |
| Haloarculaceae.1 | Haloarculaceae.2 | 77.374 | 175 | 953 |
| WSA2.gz_assembly | Haloferacaceae.1 | 77.3798 | 299 | 1248 |
| Haloarculaceae.2 | Haloarculaceae.1 | 77.4171 | 173 | 663 |
| Haloarculaceae.5 | PCN9.gz_assembly | 77.4383 | 202 | 824 |
| Haloarculaceae.3 | WSH3.gz_assembly | 77.4672 | 175 | 604 |
| Haloferacaceae.1 | WSA2.gz_assembly | 77.4823 | 282 | 713 |
| Haloarculaceae.5 | Haloarculaceae.3 | 77.5149 | 163 | 824 |
| WSA2.gz_assembly | Haloarculaceae.4 | 77.5401 | 334 | 1248 |

| | | | | |
|---|---|---|---|---|
| Haloarculaceae.4 | WSA2.gz_assembly | 77.5402 | 325 | 678 |
| WSH3.gz_assembly | Haloarculaceae.1 | 77.6034 | 272 | 1077 |
| WSH3.gz_assembly | Haloarculaceae.5 | 77.654 | 273 | 1077 |
| Haloarculaceae.4 | Haloarculaceae.5 | 77.6572 | 243 | 678 |
| Haloarculaceae.5 | Haloarculaceae.4 | 77.6658 | 245 | 824 |
| Haloarculaceae.3 | Haloarculaceae.5 | 77.6723 | 164 | 604 |
| Haloarculaceae.1 | WSH3.gz_assembly | 77.7218 | 250 | 953 |
| PCN9.gz_assembly | WSA2.gz_assembly | 77.7903 | 357 | 981 |
| Haloferacaceae.1 | Haloferacaceae.2 | 77.7996 | 147 | 713 |
| Haloferacaceae.2 | Haloferacaceae.1 | 77.8223 | 144 | 416 |
| Haloarculaceae.5 | WSH3.gz_assembly | 77.9163 | 272 | 824 |
| WSA2.gz_assembly | PCN9.gz_assembly | 77.93 | 348 | 1248 |
| Haloarculaceae.1 | Haloarculaceae.5 | 78.4522 | 267 | 953 |
| Haloarculaceae.5 | Haloarculaceae.1 | 78.6182 | 271 | 824 |
| Halovenus | WSH3.gz_assembly | 79.7953 | 340 | 605 |
| WSH3.gz_assembly | Halovenus | 79.933 | 331 | 1077 |
| Halovenus | Haloarculaceae.3 | 81.4079 | 105 | 605 |
| Haloarculaceae.3 | Halovenus | 81.5705 | 111 | 604 |
| Haloarculaceae.5 | Haloarculaceae.2 | 85.8727 | 359 | 824 |
| Haloarculaceae.2 | Haloarculaceae.5 | 85.9246 | 370 | 663 |

***Table S6***

| Gene ID | Gene | Best BLAST Hit | Acession # | % ID | E-value |
|---|---|---|---|---|---|
| *12B.1 Metagenome* | | | | | |
| c_000000339582_1 | *nifD* | *Chloroflexaceae bacterium* (stromatolite metagenome) | NJN65560.1 | 56.36 | 1.00E-138 |
| c_000000029304_3 | *nifD* | *Chloroflexaceae bacterium* (stromatolite metagenome) | NJN65560.1 | 57.2 | 3.00E-100 |
| c_000000286385_1 | *nifD* | *Chloroflexaceae bacterium* (stromatolite metagenome) | NJN65560.1 | 55.86 | 2.00E-129 |
| c_000000146019_1 | *nifH* | *Halorhodospira halochloris* | BAU57913.1 | 84.11 | 5.00E-89 |
| c_000000150855_1 | *nifH* | *Roseiflexus sp. RS-1* | WP_011955961.1 | 75.68 | 3.00E-99 |
| c_000000412048_2 | *nifH* | *Roseiflexus castenholzii* | WP_012122497.1 | 70.09 | 4.00E-44 |
| c_000000419371_3 | *nifH* | *Roseiflexus sp. RS-1* | WP_011955961.1 | 72.69 | 2.00E-142 |
| c_000000029304_2 | *nifK* | archaeon BMS3Bbin16 | GBE57063.1 | 54.3 | 0 |
| c_000000266741_1 | *nifK* | archaeon BMS3Abin16 | GBE18617.1 | 56.92 | 8.00E-50 |
| c_000000348638_1 | *nifk* | archaeon BMS3Abin16 | GBE18617.1 | 54.92 | 1.00E-40 |
| c_000000476483_1 | *nifk* | archaeon BMS3Bbin16 | GBE57063.1 | 56.07 | 2.00E-32 |
| *67B.1 Metagenome* | | | | | |
| c_000000142325_1 | *nifD* | *Rhodovibrio salinarum* | WP_027289759.1 | 90.7 | 1.00E-47 |
| *12B.2 Metagenome* | | | | | |
| c_000000324579_1 | *nifD* | *Rhodovibrio salinarum* | WP_027289759.1 | 84.58 | 0.00E+00 |
| c_000000324579_2 | *nifH* | *Rhodoplanes serenus* | WP_155481001.1 | 81.82 | 1.00E-16 |
| c_000000210854_1 | *nifH* | *Rhodovibrio salinarum* | WP_027289758.1 | 90.87 | 1.00E-154 |
| c_000000219157_1 | *nifk* | *Rhodovibrio salinarum* | WP_027289760.1 | 83.64 | 7.00E-134 |
| *33.2 Metagenome* | | | | | |
| c_000000151043_1 | *nifD* | *Deltaproteobacteria bacterium* (wastewater metagenome) | NCC26223.1 | 78 | 3.00E-143 |
| c_000000156957_1 | *nifD* | *Desulfovibrio brasiliensis* | WP_054649226.1 | 78.31 | 4.00E-38 |
| c_000000055755_1 | *nifH* | *Desulfobulbus sp. Tol-SR* | KGO35492.1 | 88.89 | 3.00E-49 |
| c_000000074817_2 | *nifH* | *Deltaproteobacteria bacterium* (wastewater metagenome) | NCC25938.1 | 82.78 | 2.00E-170 |
| c_000000066890_1 | *nifk* | *Desulfovibrio gilichinskyi* | WP_085099001.1 | 78.9 | 1.00E-59 |
| c_000000105010_1 | *nifk* | *Desulfocurvus vexinensis* | WP_028586618.1 | 82.76 | 2.00E-49 |
| *35.2 Metagenome* | | | | | |
| c_000000163780_1 | *nifD* | Candidatus *Electrothrix aarhusiensis (Desulfbulbaceae)* | RWX47654.1 | 71.17 | 9.00E-85 |
| c_000000144626_1 | *nifD* | *Rhodovibrio salinarum* | WP_027289759.1 | 83.33 | 2.00E-96 |
| c_000000242108_1 | *nifk* | *Halorhodospira halophila* | WP_011813089.1 | 76.42 | 2.00E-53 |

*Table S7*

| Gene ID | Gene Name | Best BLAST Hit | Acession # | % ID | E-value |
|---|---|---|---|---|---|
| *12B.2 Metagenome* | | | | | |
| c_000000000782_2 | *dsrA* | uncultured Desulfohalobiaceae bacterium (MgCl2-containing environments) | ABD38984.1 | 87.805 | 2.23E-20 |
| c_000000000782_1 | *dsrB* | Desulfovermiculus halophilus | WP_027370030.1 | 81.05 | 0 |
| c_000000307376_1 | *dsrB* | Desulfovermiculus halophilus | WP_027370031.1 | 79.487 | 1.72E-113 |
| *67B.2 Metagenome* | | | | | |
| c_000000052251_1 | *dsrA* | uncultured sulfate-reducing bacterium (Mediterranean solar saltern) | ACM45532.1 | 95.489 | 4.30E-89 |
| c_000000096034_1 | *dsrA* | Betaproteobacteria bacterium (groundwater environment) | MBI3042764.1 | 61.224 | 8.99E-34 |
| c_000000263801_1 | *dsrA* | uncultured sulfate-reducing bacterium (Mediterranean solar saltern) | ACM45532.1 | 91.71 | 3.68E-128 |
| c_000000328055_1 | *dsrA* | Desulfovermiculus halophilus | WP_027370030.1 | 84.516 | 5.50E-80 |
| c_000000431746_4 | *dsrA* | uncultured Desulfohalobiaceae bacterium (MgCl2-containing environments) | ABD38987.1 | 98 | 1.04E-25 |
| c_000000552184_1 | *dsrA* | Desulfovermiculus halophilus | WP_027370030.1 | 83.133 | 4.03E-144 |
| c_000000263801_2 | *dsrB* | Desulfovermiculus halophilus | WP_027370031.1 | 82.544 | 0 |
| c_000000423115_1 | *dsrB* | Gammaproteobacteria bacterium HGW-Gammaproteobacteria-1 (groundwater metagenome) | PKM45266.1 | 78.431 | 4.51E-55 |
| c_000000431746_3 | *dsrB* | Desulfovermiculus halophilus | WP_027370031.1 | 84.557 | 0 |
| c_000000652016_2 | *dsrB* | Candidatus Reidiella endopervernicosa | WP_078483318.1 | 74.638 | 5.54E-71 |
| *29.2 Metagenome* | | | | | |
| c_000000349271_2 | *dsrA* | Desulfovermiculus halophilus | WP_027370030.1 | 81.507 | 0 |
| c_000000158655_1 | *dsrB* | Desulfovermiculus halophilus | WP_027370031.1 | 78.824 | 6.07E-42 |
| c_000000349271_1 | *dsrB* | Desulfovermiculus halophilus | WP_027370031.1 | 83.578 | 0 |
| *33.2 Metagenome* | | | | | |
| c_000000095341_6 | *dsrA* | Desulfovermiculus halophilus | WP_027370030.1 | 81.735 | 0 |
| c_000000115348_1 | *dsrA* | uncultured sulfate-reducing bacterium (Mediterranean solar saltern) | ACM45532.1 | 98.013 | 1.01E-103 |
| c_000000154383_1 | *dsrA* | Desulfovermiculus halophilus | WP_027370030.1 | 84.131 | 0 |
| c_000000440998_1 | *dsrA* | uncultured sulfate-reducing bacterium (Mediterranean solar saltern) | ACM45532.1 | 94.841 | 7.52E-179 |
| c_000000470085_1 | *dsrA* | Desulfovermiculus halophilus | WP_027370030.1 | 83.784 | 2.90E-75 |
| c_000000095341_7 | *dsrB* | Desulfovermiculus halophilus | WP_027370031.1 | 82.544 | 0 |
| c_000000099172_1 | *dsrB* | Desulfovibrionales bacterium (anaerobic digester metagenome) | NLW80111.1 | 84.706 | 1.53E-50 |
| c_000000115348_2 | *dsrB* | uncultured sulfate-reducing bacterium (Mediterranean solar saltern) | ACM45533.1 | 94.924 | 2.14E-142 |
| c_000000138427_1 | *dsrB* | Desulfovermiculus halophilus | WP_027370031.1 | 83.465 | 2.03E-73 |
| c_000000182614_1 | *dsrB* | Desulfovermiculus halophilus | WP_027370031.1 | 84.255 | 1.03E-150 |
| *35.1 Metagenome* | | | | | |
| c_000000234629_1 | *dsrA* | Halorhodospira halophila | WP_201171943.1 | 90.323 | 0 |
| c_000000667957_1 | *dsrA* | Thiobacillus denitrificans | WP_011311881.1 | 70.27 | 1.11E-29 |
| c_000000698539_6 | *dsrA* | Desulfovermiculus halophilus | WP_027370030.1 | 81.735 | 0 |
| c_000000698539_5 | *dsrB* | Desulfovermiculus halophilus | WP_027370031.1 | 82.045 | 0 |
| *35.2 Metagenome* | | | | | |
| c_000000146673_4 | *dsrA* | uncultured sulfate-reducing bacterium (Mediterranean solar saltern) | ACM45532.1 | 93.214 | 0 |
| c_000000196400_16 | *dsrA* | Desulfovermiculus halophilus | WP_027370030.1 | 81.34 | 1.94E-113 |
| c_000000146673_3 | *dsrB* | Desulfovermiculus halophilus | WP_027370031.1 | 82.294 | 0 |
| c_000000507418_1 | *dsrB* | uncultured sulfate-reducing bacterium (Mediterranean solar saltern) | ACM45525.1 | 94.624 | 1.53E-129 |
| *35.3 Metagenome* | | | | | |
| c_000000154008_1 | *dsrA* | Desulfovermiculus halophilus | WP_027370030.1 | 82.609 | 3.86E-146 |
| c_000000154017_1 | *dsrA* | Desulfovermiculus halophilus | WP_027370030.1 | 86.611 | 7.69E-144 |
| c_000000699647_1 | *dsrA* | uncultured sulfate-reducing bacterium (Mediterranean solar saltern) | ACM45524.1 | 93.798 | 6.79E-83 |
| c_000000614952_1 | *dsrB* | uncultured sulfate-reducing bacterium (Mediterranean solar saltern) | ACM45543.1 | 85.143 | 2.36E-110 |
| c_000000743744_3 | *dsrB* | Desulfovermiculus halophilus | WP_027370031.1 | 83.399 | 6.52E-160 |
| *56.2 Metagenome* | | | | | |
| c_000000223435_2 | *dsrA* | Desulfohalobium retbaense | WP_015750705.1 | 80.899 | 6.03E-90 |
| c_000000481738_1 | *dsrA* | uncultured sulfate-reducing bacterium (Mediterranean solar saltern) | ACM45532.1 | 90.526 | 1.36E-55 |
| c_000000296478_1 | *dsrB* | Desulfovibrio sp. (anaerobic digester metagenome) | NMC50171.1 | 82 | 5.35E-58 |
| c_000000481738_2 | *dsrB* | uncultured sulfate-reducing bacterium (Mediterranean solar saltern) | ACM45535.1 | 86.512 | 1.81E-142 |
| *56.3 Metagenome* | | | | | |
| c_000000266214_1 | *dsrA* | Desulfonatronovibrio magnus | WP_045212625.1 | 82.639 | 5.23E-68 |
| c_000000310384_1 | *dsrA* | uncultured sulfate-reducing bacterium (Mediterranean solar saltern) | ACM45532.1 | 90 | 6.89E-73 |
| c_000000083255_1 | *dsrB* | Candidatus Reidiella endopervernicosa | WP_078483318.1 | 75 | 8.36E-34 |
| c_000000129612_1 | *dsrB* | Sulfurifustis variabilis | WP_096457663.1 | 77.348 | 2.88E-103 |
| c_000000310384_2 | *dsrB* | uncultured sulfate-reducing bacterium (Mediterranean solar saltern) | ACM45533.1 | 89.706 | 1.46E-41 |
| *46.2 Metagenome* | | | | | |
| c_000000109617_1 | *dsrA* | uncultured prokaryote (GSL sediments) | ABD46878.1 | 94.958 | 8.99E-78 |
| c_000000109642_9 | *dsrA* | Desulfovermiculus halophilus | WP_027370030.1 | 81.579 | 3.78E-154 |
| c_000000123025_2 | *dsrA* | Desulfomicrobium norvegicum | WP_092194128.1 | 74.576 | 2.38E-23 |
| c_000000169346_1 | *dsrA* | Gammaproteobacteria bacterium (hydrothermal sediment) | RKZ39678.1 | 72.678 | 8.41E-96 |
| c_000000196300_1 | *dsrA* | uncultured sulfate-reducing bacterium (Mediterranean solar saltern) | ACM45532.1 | 97.143 | 1.23E-94 |
| c_000000206341_1 | *dsrA* | Desulfonatronovibrio magnus | WP_045212625.1 | 83.333 | 6.74E-35 |
| c_000000225585_8 | *dsrA* | uncultured sulfate-reducing bacterium (Mediterranean solar saltern) | ACM45532.1 | 91.365 | 0 |
| c_000000388521_2 | *dsrA* | uncultured sulfate-reducing bacterium (Mediterranean solar saltern) | ACM45532.1 | 95.652 | 4.07E-39 |
| c_000000388564_2 | *dsrA* | Desulfovermiculus halophilus | WP_027370030.1 | 82.877 | 0 |
| c_000000442685_1 | *dsrA* | uncultured sulfate-reducing bacterium (Mediterranean solar saltern) | ACM45532.1 | 99.057 | 2.05E-70 |
| c_000000483844_2 | *dsrA* | uncultured microorganism (hydrothermal fluid) | AJA06229.1 | 62.903 | 2.71E-20 |
| c_000000537023_1 | *dsrA* | Archaeoglobaceae archaeon (Mid-Cayman Rise Vent Fluids) | HID42311.1 | 68.77 | 6.00E-168 |
| c_000000544283_1 | *dsrA* | uncultured prokaryote (GSL sediments) | ABD46878.1 | 93.45 | 3.67E-161 |
| c_000000544595_1 | *dsrA* | uncultured prokaryote (GSL sediments) | ABD46870.1 | 93.023 | 2.23E-81 |
| c_000000655084_1 | *dsrA* | uncultured prokaryote (GSL sediments) | ABD46878.1 | 91.818 | 6.03E-69 |
| c_000000069099_1 | *dsrB* | Archaeoglobus sp. (hot spring sediment) | HGQ78171.1 | 76.033 | 5.44E-63 |
| c_000000122840_1 | *dsrB* | Betaproteobacteria bacterium (soil metagenome) | HAN56732.1 | 75.806 | 3.70E-26 |
| c_000000225585_7 | *dsrB* | Desulfovermiculus halophilus | WP_027370031.1 | 82.544 | 0 |
| c_000000276611_1 | *dsrB* | Magnetococcales bacterium (lake sediments) | MBF0613955.1 | 73.786 | 2.98E-48 |
| c_000000312408_3 | *dsrB* | Desulfobacteraceae bacterium SEEP-SAG9 (marine sediment) | PQP34394.1 | 87.273 | 5.54E-31 |
| c_000000388521_1 | *dsrB* | uncultured sulfate-reducing bacterium (Mediterranean solar saltern) | ACM45525.1 | 94.286 | 6.10E-96 |
| c_000000388564_1 | *dsrB* | uncultured sulfate-reducing bacterium (Mediterranean solar saltern) | ACM45533.1 | 95 | 4.59E-96 |
| c_000000483844_1 | *dsrB* | uncultured microorganism (hydrothermal fluid) | AJA06230.1 | 61.224 | 2.29E-64 |
| c_000000539054_3 | *dsrB* | Desulfovermiculus halophilus | WP_027370031.1 | 86.093 | 0 |

| | | | | | |
|---|---|---|---|---|---|
| c_000000545959_1 | *dsrB* | Desulfovibrio ferrophilus | WP_126380811.1 | 84.685 | 6.96E-65 |
| c_000000658443_1 | *dsrB* | Desulfohalophilus alkaliarsenatis | AFJ04155.1 | 88.462 | 3.42E-63 |
